# Long-read transcriptomics of purified human cortical cell types exposes glial isoform complexity and disease-relevant transcript architecture

**DOI:** 10.1101/2025.11.25.690524

**Authors:** Andy Yang, Miguel Rodriguez de los Santos, Alexey Kozlenkov, Ramu Vadukapuram, Yasmin Hurd, Stella Dracheva, Jack Humphrey, Michael S. Breen

## Abstract

Alternative splicing generates extraordinary transcriptomic complexity in the human brain, yet the full-length isoform landscape across human cortical cell types remains uncharted. Combining fluorescence-activated nuclei sorting with long- and short-read RNA sequencing, we generated isoform-resolved transcriptomes for five major lineages of the adult human prefrontal and orbitofrontal cortex: GABAergic neurons, glutamatergic neurons, oligodendrocytes, astrocytes, and microglia. We cataloged over 220,000 full-length isoforms, ∼35-56% previously unannotated; novel transcripts were longer, more exon-rich, and predominantly protein-coding. Contrary to the neuron-centric view of cortical complexity, glial lineages, particularly oligodendrocytes and microglia, emerged as the most isoform-diverse populations in the cortex. Differential transcript usage and dominant isoform switching defined cell identity, with ∼59-62% of differentially regulated transcripts absent from current annotations. Critically, pathogenic variants were enriched >2-fold at novel splice boundaries within disease genes including *POGZ*, *TARDBP*, and *PLP1*, establishing isoform selection as a primary axis of cortical identity and exposing a layer of pathogenic variation invisible to canonical gene annotations.

## INTRODUCTION

Alternative splicing (AS) is a fundamental mechanism of post-transcriptional regulation, particularly expanded in the brain, where it governs neuronal differentiation, synaptic plasticity, and circuit formation^1,2^. Splicing patterns exhibit remarkable specificity across developmental stages, cortical regions, and cell types^3–5^, enabling precise control of neuronal and glial gene expression programs essential for normal brain function^6,7^. Disruption of AS is increasingly recognized as a molecular hallmark of neuropsychiatric and neurodegenerative disorders, including autism spectrum disorder (ASD)^8^, schizophrenia (SZ)^9^, and Alzheimer’s disease (AD)^10–11^. Resolving the full repertoire of AS events across defined brain cell types is therefore essential for identifying disease-relevant isoforms and potential therapeutic targets.

Recent advances in long-read sequencing have transformed transcriptome annotation by capturing full-length isoforms, resolving complex exon connectivity, and accurately identifying AS events^12–14^. In animal models, long-read sequencing has revealed tens of thousands of previously unannotated isoforms and extensive region- and cell type-specific splicing patterns in key brain areas^15–17^. Comparative studies further underscore the functional relevance of these isoforms, identifying human-enriched splicing events that may underlie higher-order cognitive function^18–20^. In humans, long-read approaches have mapped over 200,000 unique isoforms during mid-gestation^21^, delineated cell-specific splicing programs in the adult frontal cortex^22^, and linked isoform-level changes directly to Alzheimer’s pathology^10–11^. Despite these advances, existing human datasets remain dominated by bulk tissue or developmental samples, limiting resolution of cell type-specific isoform architecture in the adult brain. Single-nucleus long-read studies capture only a few hundred thousand nuclei across limited cell populations^5,15,17^, constraining both isoform discovery depth and cell type resolution.

Here, we show that cortical cell identity is encoded as much by which isoforms of a gene are used as by which genes are expressed, and that this isoform-level encoding directly shapes the genomic interpretation of disease risk. By integrating fluorescence-activated nuclei sorting (FANS) with PacBio Iso-Seq long-read sequencing and matched short-read RNA-seq across the adult human dorsolateral (DLPFC) and orbitofrontal (OFC) cortex, we generated a cell type-resolved reference of full-length transcripts spanning five major cortical lineages: MGE-derived GABAergic neurons, glutamatergic neurons, oligodendrocytes, astrocytes, and microglia. Transcript specialization, not gene expression alone, emerges as a defining feature of cortical cell identity. Critically, pathogenic variants are concentrated within the splice-boundary classes that drive transcript novelty, exposing a hidden layer of disease-relevant exonic sequences that canonical gene annotations miss entirely, with implications for how variants in established neurological disease genes should be interpreted.

## RESULTS

### Comprehensive long-read transcriptome profiling and cross-platform validation across human cortical cell types

To define the full-length isoform landscape across major cortical cell types, we generated a cell type-resolved, full-length transcriptome atlas of the adult human cortex. Long-read transcriptomes were generated from fluorescence-activated nuclei sorting (FANS)-purified populations representing five principal cortical lineages: MGE-derived GABAergic neurons (GABA), glutamatergic neurons (GLU), oligodendrocytes (OLIG), astrocytes (AST), and microglia (MG) isolated from DLPFC (n = 3 donors) and OFC (n = 4 donors) (**Figure 1**). Within each region, long-read sequencing was performed for GABA, GLU, and OLIG nuclei, while AST and MG were profiled exclusively in the OFC cohort to complete representation of the five major cortical cell classes. Because cohorts were generated in separate experimental batches, all primary analyses were performed within-region, with cross-region comparisons serving as orthogonal validation of shared biological patterns.

**Figure 1.**
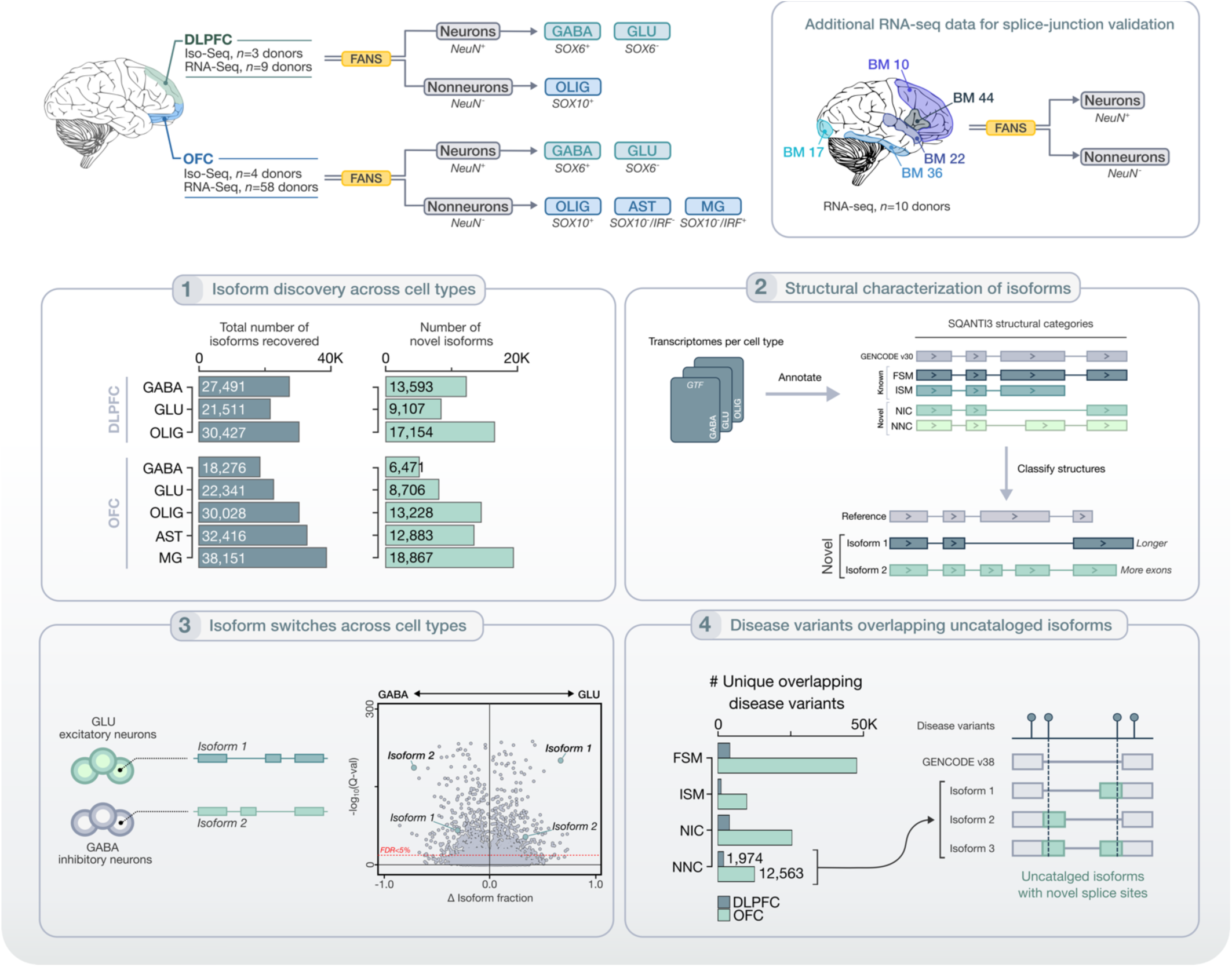
Experimental workflow for cell type-resolved long-read isoform profiling in human cortex. Postmortem human dorsolateral prefrontal cortex (DLPFC) and orbitofrontal cortex (OFC) were processed using fluorescence-activated nuclei sorting (FANS) to isolate five major cortical lineages. Neuronal nuclei (NeuN⁺) were subdivided into MGE-derived inhibitory neurons (GABA; SOX6⁺) and excitatory neurons (GLU; SOX6⁻). Non-neuronal nuclei (NeuN⁻) were sorted into oligodendrocytes (OLIG; SOX10⁺) in both regions, and additionally into astrocytes (AST; SOX10⁻/IRF5⁻) and microglia (MG; SOX10⁻/IRF5⁺) in OFC. DLPFC: n = 3 donors (Iso-Seq), n = 9 donors (RNA-seq); OFC: n = 4 donors (Iso-Seq), n = 58 donors (RNA-seq). An independent cohort of NeuN⁺ and NeuN⁻ nuclei from five cortical regions (BM10, BM17, BM22, BM36, BM44; n = 10 donors) provided orthogonal splice-junction support. PacBio HiFi Iso-Seq libraries were prepared from each sorted population. Full-length transcript models were processed through SQANTI3 for structural classification and splice-junction validation against GENCODE v38. Inset panels summarize four key analytical outputs: (1) isoform discovery across cell types, (2) structural characterization of novel isoforms, (3) differential isoform switching, and (4) intersection of novel isoforms with HGMD disease-causing variants.

Nuclei were isolated using two previously established FANS strategies^23,24^ (**Figure S1A-B**). The first employed antibodies against NeuN (RBFOX3; neuronal marker), SOX10 (oligodendroglial marker), and SOX6 (marker of MGE-derived GABAergic neurons) to enrich GABA, GLU, and OLIG populations. The second used antibodies against NeuN, SOX10, and IRF5 (microglial marker) to separate microglia from other glial nuclei and enable recovery of astrocyte-enriched populations. Population purity was previously validated using bulk RNA-seq and ChIP-seq^23,24^. For long-read sequencing, ∼40,000 nuclei were profiled per sample using PacBio Iso-Seq. Across all samples, Iso-Seq libraries generated 23.1 million and 50.5 million circular consensus sequence (CCS) reads from DLPFC and OFC, respectively, corresponding to an average of 7.71 ± 1.05 million CCS reads per cell type in DLPFC and 10.1 ± 1.70 million CCS reads per cell type in OFC (**Table S1**). Mean read lengths were 3.33 kb (DLPFC) and 2.04 kb (OFC) (**Figure S2**) and correlated modestly with RNA integrity number (**Figure S3**), consistent with prior reports^25,26^. Read yield, splice-junction support, and transcript completeness were robust across all samples and cell types (**Tables S2-S3**).

To validate splice-junction architecture, long-read data were paired with matched short-read Illumina RNA-seq from FANS-derived GABA, GLU, and OLIG nuclei in DLPFC (n = 9 donors; ∼125M reads/sample)^23,27^ and OFC (n = 58 donors; ∼131M reads/sample)^28^ (**Table S3**). Cell type deconvolution confirmed strong enrichment for target populations and minimal non-target contamination, independently validating sort purity (**Figure S1C-D**). For AST and MG, which lacked matched short-read data, we incorporated an independent deeply sequenced reference: Illumina RNA-seq from 100 NeuN⁺ and NeuN⁻ nuclei spanning five cortical regions (n = 10 donors; ∼182M reads/sample)^29^ (**Table S3**), providing orthogonal splice-junction support across all five lineages.

This integrated framework combines the structural precision of long-read sequencing with the quantitative depth of short-read RNA-seq across two cortical regions and five cell types. The long-read data contribute isoform structure and splice-junction architecture, while short-read data provide the statistical power for all quantification and lineage-level inferences.

### Quantification of high-confidence long-read isoform annotations

Isoform reconstruction followed a stringent, multi-stage filtering framework (*see* **Figure S4** *for full details*). After quality control, 76% of DLPFC and 60% of OFC circular consensus sequence (CCS) reads achieved Q50 (99.9%) accuracy. Alignment to the GRCh38 genome yielded ∼1.0 million and ∼2.7 million full-length non-chimeric (FLNC) reads in the DLPFC and OFC, respectively (**Figure S4**, **Table S1**). Isoforms were then reconstructed using TAMA and refined with matched short-read RNA-seq through SQANTI3, which removed transcripts with weak junction support or signatures of technical artifact, including poly(A) intra-priming and reverse transcriptase switching (**Figure S5**). All downstream analyses were restricted to these high-confidence isoforms.

### Glial cells harbor the greatest isoform diversity of any major cortical lineage

Sequencing saturation analysis revealed a critical asymmetry (**Figure S6**). Gene detection approached saturation in every lineage, whereas isoform detection increased nearly linearly with depth and showed no evidence of plateauing. Differences in isoform diversity therefore reflect genuine combinatorial complexity of splice-junction usage and transcript boundaries rather than sampling variation, consistent with prior long-read studies of human brain^22^ and the known abundance of lowly expressed and cell-state-restricted transcripts. Gene-level detection was highly reproducible across donors, with 51–55% of genes identified in every donor within each region (**Figure S7**).

All cell type comparisons were performed within-region to eliminate batch effects. In the DLPFC, we identified 79,429 unique isoforms corresponding to 10,135 annotated genes and 140 unknown gene loci (**Table 1, Figure 2**). Isoform diversity was highest in OLIG (30,427), followed by GABA (27,491) and GLU (21,511) (**Figure 2A**). Of these, 32,954 isoforms (∼41%) were restricted to a single cell type, whereas only 9,023 (∼11%) were shared across all three populations (**Figure 2C**), indicating that most isoform expression is lineage-specific rather than constitutive, consistent with prior long-read studies of the human brain^5,21^. A similar pattern was observed at the gene level, where ∼10% of expressed genes exhibited cell type-restricted detection (**Figure 2D, Table S4**), including canonical lineage markers for GABA (*GAD1*, *GAD2*, *SLC32A1*), GLU (*SLC17A7*, *SLC17A6*, *SATB2*), and OLIG (*OLIG2*, *MBP*, *PLP1*).

**Figure 2.**
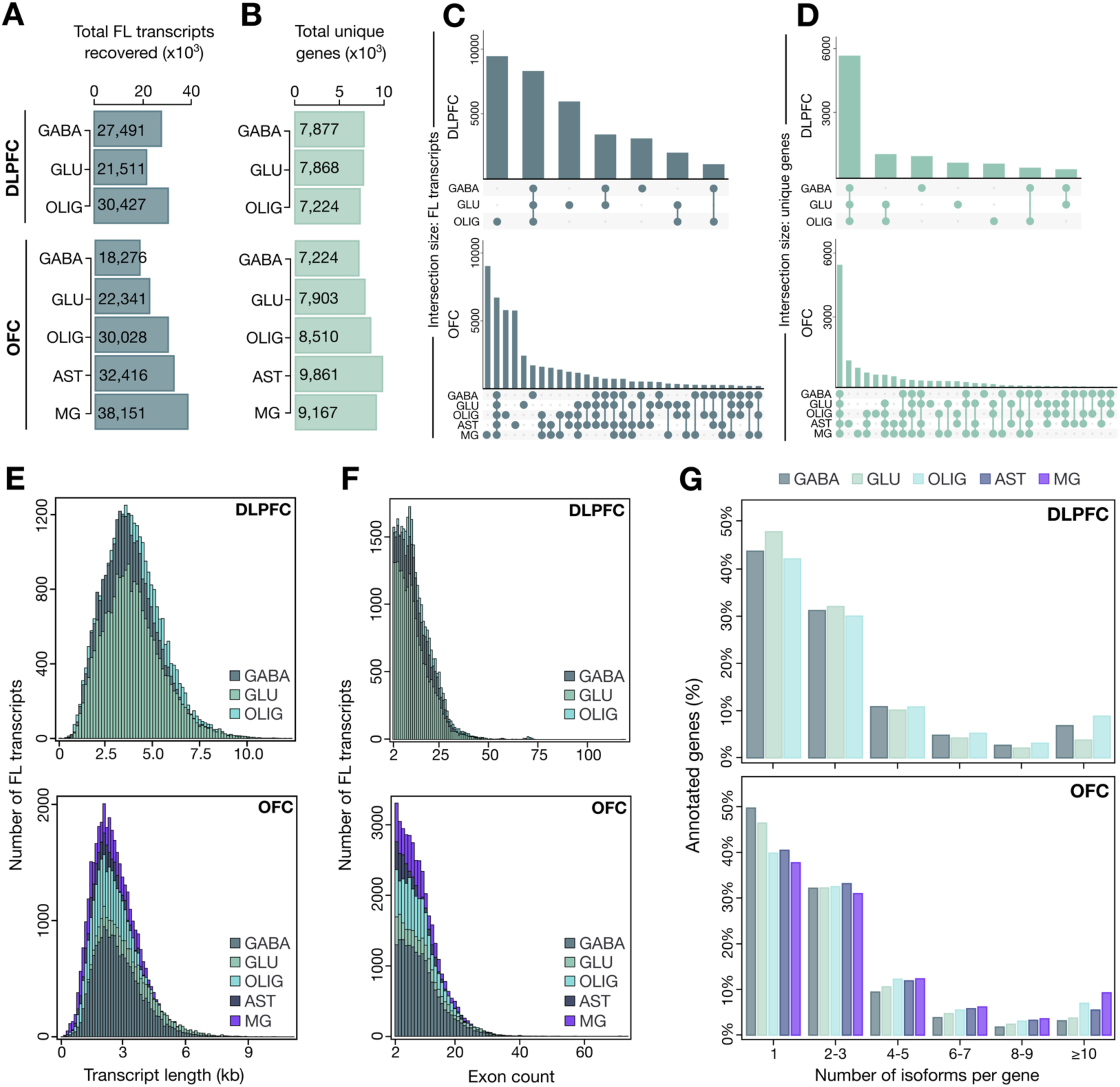
Glial populations harbor the greatest isoform diversity of any major cortical lineage. (**A**) Total full-length (FL) transcripts recovered per cell type in DLPFC and OFC. (**B**) Total unique annotated genes detected per cell type. UpSet plots showing overlap of (**C**) FL isoforms and (**D**) annotated genes across cell types within each region. (**E**) Transcript length distributions (kb) per cell type in DLPFC and OFC. (**F**) Exon count distributions per transcript across cell types. (**G**) Distribution of isoform number per gene across cell types.

**Table 1.** Summary of full-length isoform discovery, annotation status, and coding potential across cortical cell types.

|  |  |  |  |  |  |  |  |  |
| --- | --- | --- | --- | --- | --- | --- | --- | --- |
|  | **Dorsolateral Prefrontal Cortex** |  |  | **Orbitofrontal Cortex** |  |  |  |  |
|  | **GABA** | **GLU** | **OLIG** | **GABA** | **GLU** | **OLIG** | **AST** | **MG** |
| Total isoforms | 27491 | 21511 | 30427 | 18276 | 22341 | 30028 | 32416 | 38151 |
| # FSM | 12460 | 11292 | 11528 | 10967 | 12436 | 15143 | 17579 | 16639 |
| # ISM | 908 | 768 | 1009 | 698 | 928 | 1263 | 1439 | 1867 |
| # NIC | 10979 | 7898 | 13155 | 5541 | 7389 | 10594 | 10560 | 15097 |
| # NNC | 2614 | 1209 | 3999 | 930 | 1317 | 2634 | 2323 | 3770 |
| # Other | 530 | 344 | 736 | 140 | 271 | 394 | 515 | 778 |
| # Protein coding | 26299 | 20599 | 29212 | 17088 | 20807 | 27714 | 29791 | 34843 |
| # Non-protein coding | 1192 | 912 | 1215 | 1188 | 1534 | 2314 | 2625 | 3308 |
| Annotated genes | 7877 | 7868 | 7433 | 7224 | 7903 | 8510 | 9861 | 9167 |
| Novel genes | 58 | 45 | 51 | 22 | 45 | 64 | 89 | 143 |
| FSM = Full splice match; ISM = Incomplete splice match; NIC = Novel in catalog; NNC = Novel not in catalog |  |  |  |  |  |  |  |  |

In the OFC, with a broader cell type representation, we identified 141,212 isoforms corresponding to 13,040 annotated genes and 323 novel loci, which was nearly twice the isoform count detected in the DLPFC (**Figure 2A-B**). Glial populations again exhibited the greatest isoform diversity, with MG (38,151), AST (32,416), and OLIG (30,028) exceeding both GABA (18,276) and GLU neurons (22,341) (**Figure 2A**). The OFC displayed pronounced transcriptomic specialization, with 46,451 isoforms (∼32%) cell type-restricted, including 18,290 unique to MG and 10,365 unique to OLIG, and only 6,704 isoforms (∼4.7%) shared across all five lineages (**Figure 2C**). At the gene level, ∼7% of genes showed robust cell type specificity (**Figure 2D, Table S4**), with profiles that replicated across regions and aligned with canonical lineage markers (as above), indicating high biological fidelity of the sorted populations. Cross-region reproducibility was high, with 42-47% of isoforms and 73-79% of genes shared between matched cell types (**Figure S8**).

Despite differences in isoform number, transcript structure was comparable across lineages within each region. Mean isoform length and exon count were comparable among DLPFC cell types (3,937 ± 1,530 bp; 12.4 ± 8.2 exons) and across the five OFC lineages (2,746 ± 1,123 bp; 9.4 ± 6.1 exons) (**Figure 2E-F**). Isoform counts per gene ranged from 1-329 in DLPFC and 1-271 in OFC (median = 2) and correlated positively with gene expression level (r = 0.27-0.33, *p* = 1×10⁻⁴⁸) and exon count (r = 0.13-0.31, *p* = 1×10⁻¹⁶), but not gene length (**Figure S9**). OLIG and MG exhibited the highest isoform complexity, with more isoforms per gene on average than all other cortical cell populations (**Figure 2G**). Approximately 70% of genes produced two or more isoforms and ∼6% produced ten or more (**Figure 2G**; **Table S5**), with highly spliced genes enriched for catalytic activity and RNA processing functions (**Table S6**). Collectively, these results show that glia surpass neuronal lineages in isoform complexity, a finding that overturns the neuron-centric view of cortical transcriptomics.

### Novel full-length isoforms are longer, more complex, and expand coding and structural landscape of the cortex

Using SQANTI3 classification against GENCODE v38 (**Figure 3A**), isoforms were grouped as known (full-splice match [FSM] or incomplete-splice match [ISM]), novel (novel in catalog [NIC] or novel not in catalog [NNC]), or other if the isoform were antisense, intergenic, genic, or fusion. Novel isoforms were pervasive across all lineages, and glial populations consistently harbored greater proportions than neurons (**Figure 3B-D, Table 1**). Between one-third and one-half of all detected transcripts lacked prior annotation. In the DLPFC, novel isoforms comprised ∼49% of GABA, 42% of GLU, and up to 56% of OLIG transcripts. In the OFC, novelty ranged from 35% in GABA to 50% in MG. This pervasive, cell-type-patterned annotation gap reflects a systematic expansion of the cortical transcriptome absent from current reference databases.

**Figure 3.**
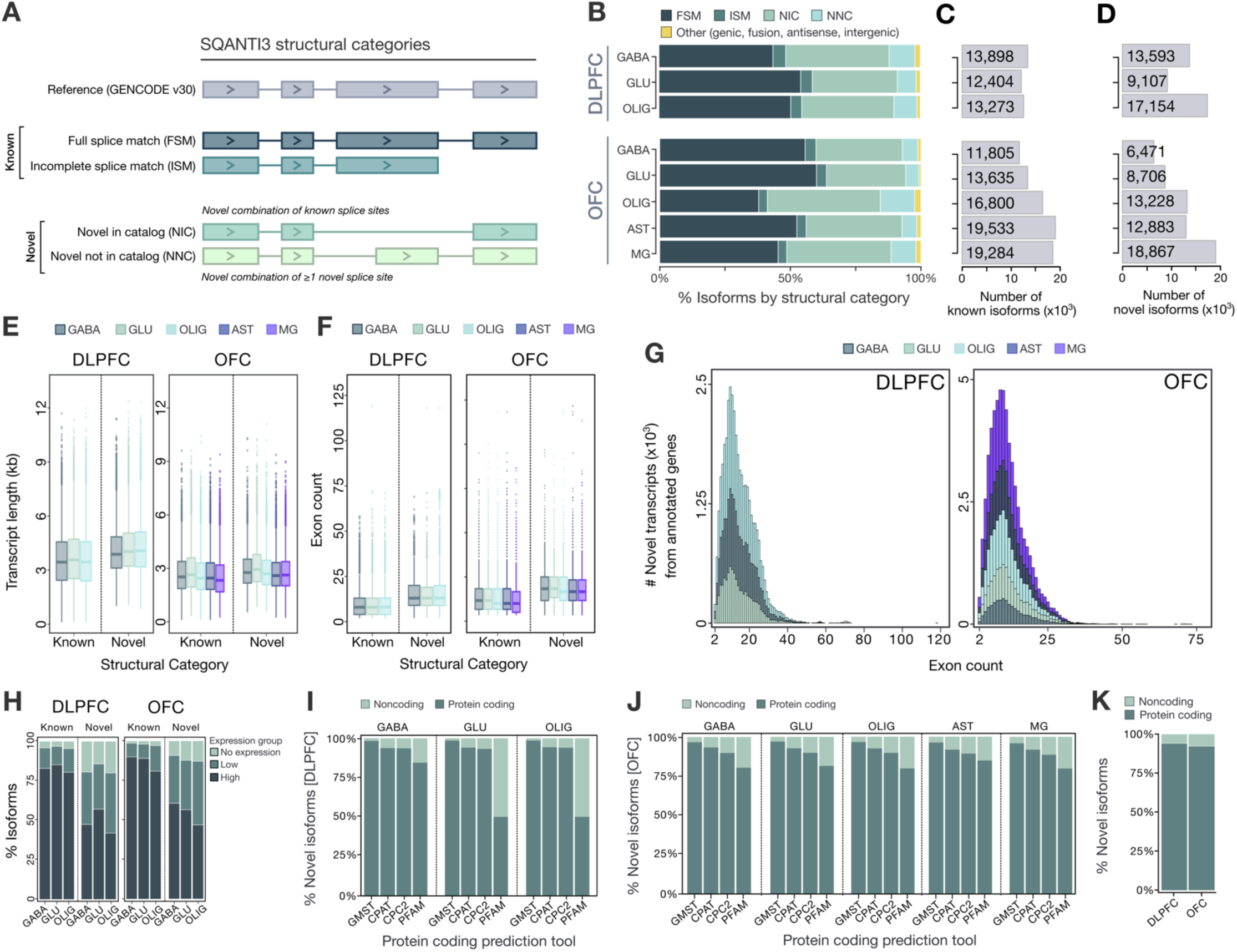
Novel full-length isoforms are structurally complex, stably expressed, and predominantly protein-coding. **(A**) SQANTI3 structural categories used to classify full-length isoforms relative to GENCODE v38: full-splice match (FSM), incomplete-splice match (ISM), novel in catalog (NIC), and novel not in catalog (NNC). (**B**) Proportional distribution of isoforms across structural classes per cell type in DLPFC and OFC. (**C-D**) Counts of known and novel isoforms per cell type in DLPFC and OFC. (**E**) Transcript length and (**F**) exon count distributions for known and novel isoforms across all cell types and regions. (**G**) Exon count distributions for novel isoforms arising within annotated gene loci, stratified by cell type. (**H**) Expression-level distributions (TPM category: no expression, low, high) for known and novel isoforms across cell types. (**I-J**) Protein-coding potential of novel isoforms in (**I**) DLPFC and (**J**) OFC, assessed by four independent frameworks (CPAT, CPC2, GMST, PFAM). (**K**) Protein-coding potential for known and novel isoforms across DLPFC and OFC.

Novel isoforms arising within annotated genes were structurally more complex than known transcripts across every cell type (**Figure 3E-F, Figure S10**). Novel isoforms were consistently longer and more multi-exonic than their annotated counterparts (e.g., in the DLPFC: novel isoforms, 4,175 ± 1,449 bp and 14.8 ± 8.2 exons; known isoforms, 3,659 ± 1,570 bp and 9.6 ± 7.3 exons). This effect was most pronounced in glial populations, particularly OLIG and MG, which contributed disproportionately to highly complex transcripts (**Figure 3G**). Although novel isoforms were on average lower in abundance, with ∼50% reaching moderate-to-high expression levels (**Figure 3H, Table S7**), they were consistent with regulated, context-dependent expression rather than transcriptional noise. These isoforms were enriched in genes central to glial biology, including cytoskeletal remodeling and membrane dynamics (*FNBP1*, *PLEKHH1*, *MAP4K4*, *ANLN*, *TTC3*), membrane transport and endolysosomal trafficking (*MYO6*, *STXBP5L*, *TMEM165*), and extracellular matrix organization (*EDIL3*, *SPARCL1*, *ERBIN*), indicating that isoform diversification meaningfully expands functional output in key cellular pathways.

More than 90% of novel isoforms were classified as protein-coding by four independent frameworks (CPAT, CPC2, GMST, PFAM; **Figure 3I-J, Figure S11**), indicating that this unannotated diversity has the potential to expand proteomic output. Novel isoforms also exhibited significantly lower minimum free energy (MFE) values than known transcripts across all cell types and regions (**Figure S12A-B**), reflecting more thermodynamically stable predicted secondary structures. This effect scaled with transcript length (|*r*| ≈ 0.86-0.87; **Figure S12C-D**) and was further modulated by expression level (**Figure S12E-F**). Novel cortical isoforms are thus structurally robust and predominantly protein-coding, expanding the functional transcriptome well beyond current annotations.

Previously unannotated genomic loci contributed only a minor fraction of overall transcript diversity (**Figure S13A**). Most were antisense to annotated genes rather than intergenic (**Figure S13B**), consistent with prior long-read RNA-sequencing reports^11,19,21^. Across all cell types, these loci were typically represented by a single isoform (**Figure S13C-D**) and were structurally compact, most commonly two-exon antisense transcripts (**Figure S13E-H**). These novel loci were also highly cell type- and region-specific, with minimal overlap across lineages or between cortical regions (**Figure S13I-L**; **Tables S8-S9**).

### Five orthogonal validation layers confirm the structural integrity and biological relevance of novel isoforms

To guard against the possibility that novel isoforms reflect technical artifacts rather than genuine biology, we implemented a five-layer orthogonal validation framework covering transcription start site (TSS) support, transcription termination site (TTS) support, full-length read confirmation, cross-dataset replication, and evolutionary constraint (**Figure 4**). Five-prime boundaries were evaluated against open chromatin peaks from four independent brain ATAC-seq datasets^30–33^ as well as CAGE peaks from FANTOM5^34–35^, while three-prime boundaries were assessed using 3′-seq polyadenylation sites and canonical polyA motifs^36^ (**Figure 4A**). Across all structural classes, >80% of transcripts received independent 5′-end support and 70-95% received 3′-end support (**Figure 4B**). Novel isoforms (NIC/NNC) matched or exceeded these rates relative to annotated transcripts, and NNC isoforms, the most structurally divergent class, showed particularly strong bilateral support. Full-length support was defined as at least one independent validation at each transcript end (Tier 2). More than 96% of NIC and NNC isoforms achieved Tier 2 status in both regions (**Figure 4C-F**), and novel isoforms achieved significantly higher Tier 2 support rates than annotated transcripts in both DLPFC (97.7% vs 88.2%; p = 2.2×10^-8^) and OFC (96.1% vs 84.1%; p = 2.3×10^-12^). Fewer than 0.03% of novel isoforms completely lacked end support at both transcript boundaries, collectively arguing against artifactual reconstruction and confirming reliable full-length recovery across known and novel structural classes.

**Figure 4.**
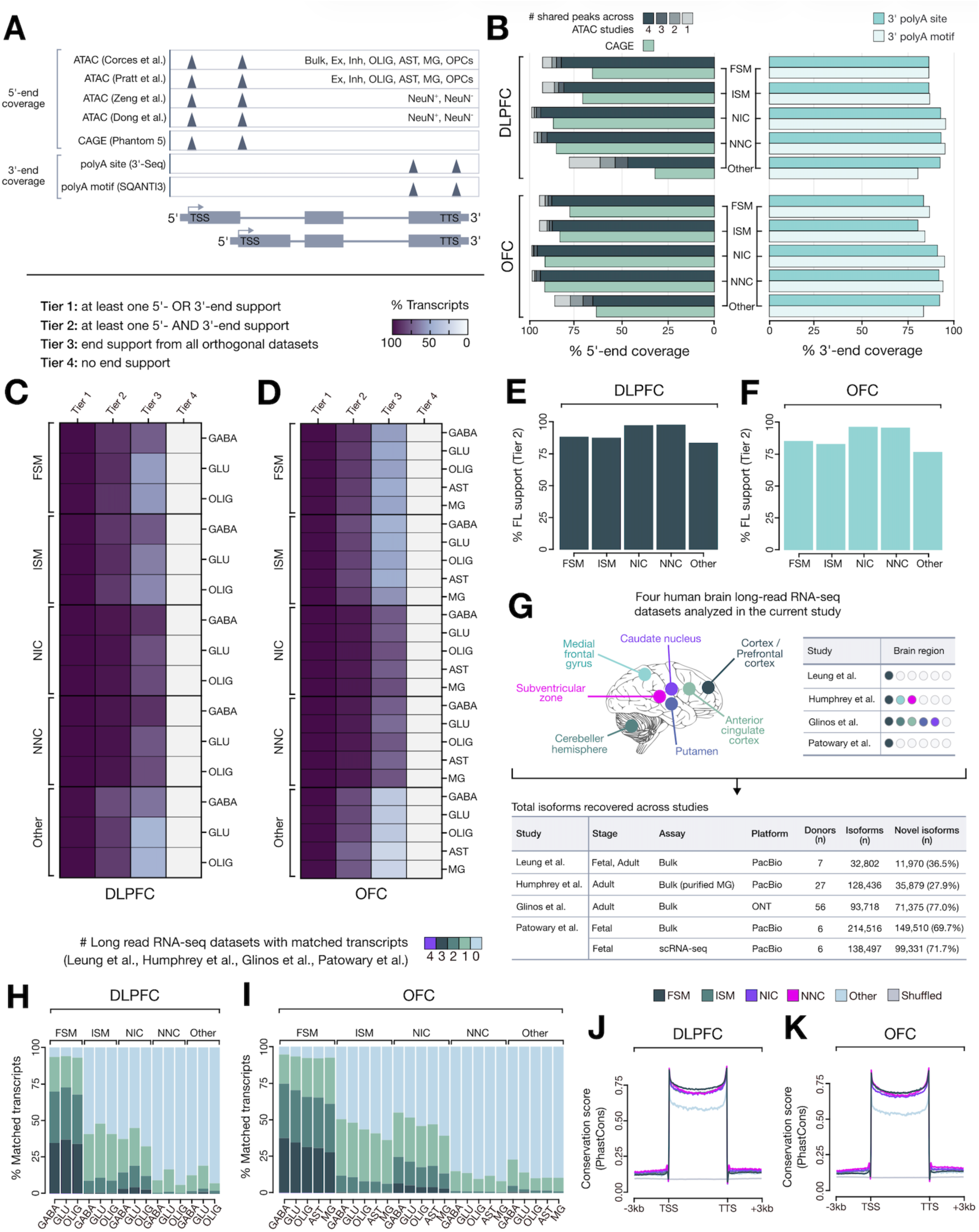
Five orthogonal validation layers confirm the structural accuracy and biological relevance of novel cortical isoforms. (**A**) Schematic of the multi-layer validation framework. Five-prime boundaries were evaluated using open chromatin peaks from four independent human brain ATAC-seq datasets and CAGE peaks from FANTOM5. Three-prime boundaries were evaluated using polyadenylation sites from 3′-seq and canonical polyA motifs identified by SQANTI3. Isoforms were assigned to four validation tiers based on support at transcript start sites (TSS) and transcript termination sites (TTS). (**B**) Percentage of isoforms with 5′-end support (left) and 3′-end support (right) across SQANTI3 structural classes in DLPFC and OFC. Bar shading indicates the number of overlapping ATAC studies; green bars indicate CAGE support; blue bars indicate polyA motif and 3′-seq support. (**C-D**) Distribution of isoforms across validation tiers per cell type and structural class in DLPFC(C) and OFC (D). (**E-F**) Percentage of isoforms achieving Tier 2 full-length support across structural classes in (**E**) DLPFC and (**F**) OFC. (**G**) Four independent human brain long-read transcriptome datasets used for cross-dataset validation, spanning fetal and adult cortex, caudate nucleus, cerebellum, and subventricular zone on PacBio and Oxford Nanopore platforms. Table summarizes study metadata, platform, sample size, and isoform recovery. (**H–I**) Proportion of transcripts with matched intron chains in independent datasets across cell types and structural classes in (**H**) DLPFC and (**I**) OFC. (**J-K**) PhastCons conservation profiles (100-way vertebrate alignment) surrounding TSS and TTS for each isoform class relative to shuffled controls in (**J**) DLPFC and (**K**) OFC.

Cross-dataset replication was tested against four independent human brain long-read transcriptome resources^11,19,21,37^ spanning fetal and adult cortex, caudate nucleus, cerebellum, and subventricular zone, generated on both PacBio and Oxford Nanopore platforms (**Figure 4G**). As expected, FSM and ISM transcripts showed the highest replication rates, with ∼92–95% recovered in at least one external dataset and ∼65–75% matched across two or more independent studies in both regions. NIC isoforms showed substantial reproducibility (∼32–55% in at least one dataset; ∼12–25% across two or more studies), while NNC isoforms, which reflect greater structural divergence and cell type specificity, showed lower but consistent recovery (∼6–17%). “Other” transcripts showed the lowest replication rates, consistent with their structural heterogeneity (**Figure 4H-I**). Replication rates for novel isoforms were comparable to those reported in prior large-scale human brain long-read studies^11,21^, arguing against technical or dataset-specific origin.

Finally, both known and novel isoforms showed marked enrichment of sequence conservation surrounding TSS and TTS relative to shuffled controls (**Figure 4J-K**). NIC and NNC isoforms displayed conservation profiles closely matching those of fully annotated FSM and ISM transcripts, indicating that previously unannotated transcript boundaries fall within evolutionarily constrained genomic regions and are unlikely to reflect stochastic transcription. The convergence of transcript-end support, full-length read confirmation, multi-platform replication, and evolutionary constraint establishes the structural accuracy and biological relevance of these novel isoforms.

### Intron retention is a pervasive, regulated feature of cortical transcriptomes and is tightly coupled to nonsense-mediated decay

Using SUPPA2, we classified seven canonical AS event types: skipping exon (SE), mutually exclusive exons (MX), alternative 5′ splice site (A5), alternative 3′ splice site (A3), retained intron (RI), alternative first exon (AF), and alternative last exon (AL) (**Figure 5A**). Retained introns (RI; ∼30-35%) and exon skipping (SE; ∼20-25%) together accounted for more than half of all detected alternative splicing events across cortical cell types (**Figure 5B-C, Figure S14A-B**). RI- and SE-containing transcripts were predominantly lowly expressed or undetectable, whereas AF- and AL-containing isoforms showed the greatest proportions of highly expressed transcripts in both regions **(Figure 5D**; **Figure S14C**). Structurally, RI isoforms were the most novel of any AS class, with the overwhelming majority classified as NIC or NNC across all cell types and regions, identifying intron retention as a primary driver of unannotated isoform diversity (**Figure 5E**; **Figure S14D**).

**Figure 5.**
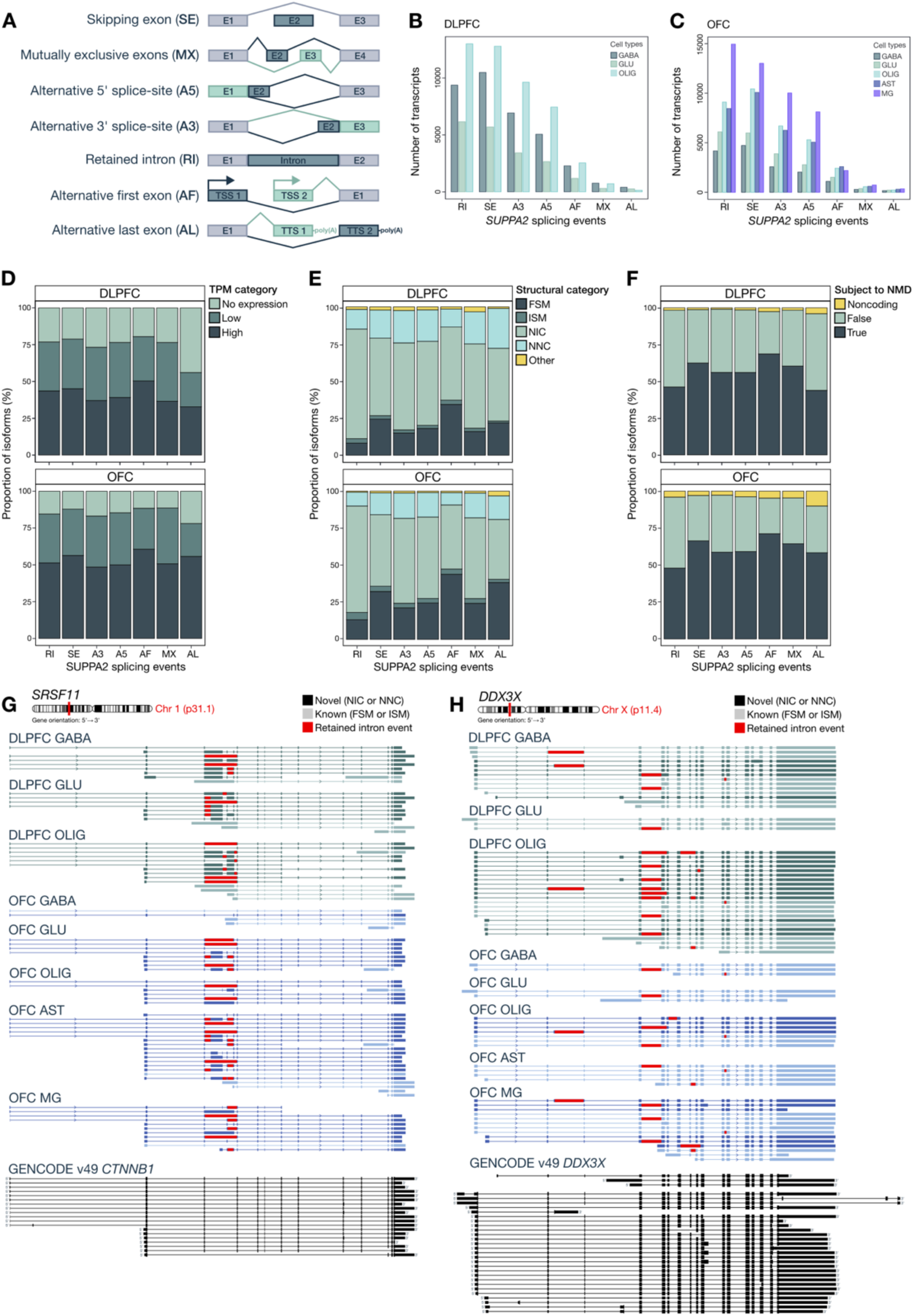
Intron retention is a dominant alternative splicing class across cortical lineages and is tightly coupled to nonsense-mediated decay. **(A)** Schematic of the seven alternative splicing event types classified by SUPPA2: retained intron (RI), skipping exon (SE), alternative 3′ splice site (A3), alternative 5′ splice site (A5), alternative first exon (AF), alternative last exon (AL), and mutually exclusive exons (MX). (**B-C**) Number of transcripts per SUPPA2 event type across cell types in (**B**) DLPFC and (**C**) OFC. (**D**) Proportion of isoforms with high, low, or undetectable expression (log₂[TPM+1]) across SUPPA2 event types in DLPFC and OFC. (**E**) Distribution of isoforms across SQANTI3 structural categories for each SUPPA2 event type, indicating that many AS events generate novel splice-junction combinations. (**F**) Proportion of isoforms predicted to undergo nonsense-mediated decay (NMD) across SUPPA2 event types in DLPFC and OFC. (**G**) Cell type-specific alternative splicing at *SRSF11* across cortical lineages. (**H**) Cell type-specific alternative splicing at *DDX3X*, with intron retention events observed alongside both NMD-predicted and stably expressed isoforms. Color coding: black, novel (NIC or NNC) isoforms; gray, known (FSM or ISM) isoforms; red, retained intron events. GENCODE v38 reference shown below for comparison.

The regulatory implications of intron retention are substantial. Across all cortical lineages, ∼54% of AS-derived transcripts were predicted NMD targets, with RI isoforms showing the strongest enrichment (**Figure 5F**). Approximately half of all RI transcripts in GABA, GLU, and OLIG populations were predicted to undergo NMD, and among protein-coding RI isoforms, a substantial fraction also remained NMD-sensitive (DLPFC: 31,652 RI isoforms [58%] from 2,086 genes; OFC: 38,968 RI isoforms [54%] from 2,605 genes) (**Table S9**). NMD-targeted transcripts were expressed at lower levels than non-NMD isoforms across all lineages **(Figure S15),** consistent with active selective degradation. Only a minority of RI-containing genes showed mutually exclusive RI and NMD states (DLPFC: 28%; OFC: 30%), indicating retention and decay are tightly coupled at the majority of loci. *MEG3* ranked among the genes with the highest number of retained-intron isoforms (**Table S10**), consistent with prior reports and its established role in RNA stability and nuclear retention^19^. Although elevated RI may be expected in nuclear RNA datasets, several converging observations support a regulated rather than purely technical origin: the RI fraction observed here (∼31%) is only modestly higher than in prior long-read studies of bulk human brain (∼23-27%)^19,37^. RI events are cell type-specific, tightly coupled with NMD, a substantial subset of RI transcripts is stably expressed and predicted to be protein-coding.

Representative loci illustrate how these mechanisms diversify gene output (**Figure 5G-H**). In *SRSF11*, a core splicing regulator, we observed multiple cell-type-specific exon-skipping and intron-retention events, generating distinct neuronal and glial isoforms (**Figure 5G**), consistent with altered autoregulatory control of splicing efficiency. *CTNNB1* showed coordinated exon-skipping and cell-type-restricted intron-retention events that generated distinct isoform repertoires across neuronal and glial lineages (**Figure S16**). In *DDX3X*, pervasive RI across OLIG and MG simultaneously generated NMD-sensitive isoforms alongside stably expressed variants (**Figure 5H**), illustrating how a single locus can deploy intron retention for both transcript surveillance and functional diversification in a cell type-dependent manner.

### Isoform-level regulation defines cortical cell identity beyond gene expression

A central question is how cell type-specific identity is encoded and whether these differences arise at the level of gene expression alone or are fundamentally driven by isoform-specific regulation. To address this with appropriate statistical power, we leveraged the large matched short-read cohorts as the quantitative backbone for all differential analyses, with long-read isoform models providing the structural reference against which transcript-level changes were measured. We performed integrated differential gene expression (DGE), differential transcript expression (DTE), and differential transcript usage (DTU) analyses on merged, region-specific transcriptome annotations incorporating cell type-specific isoform models (**Figure S17**), with donor modeled as a repeated measure (**Tables S11-S13**). Analyses were restricted to GABA, GLU, and OLIG, the three lineages with matched long- and short-read data in both regions (DLPFC, n = 9; OFC, n = 58 donors) to ensure robust splice-junction validation and directly comparable quantification across regions.

At the gene level, transcriptional divergence was greatest between neurons and oligodendrocytes, whereas GABA and GLU neurons remained comparatively similar. In the DLPFC, most DEGs contained at least one novel isoform, with a substantial fraction harboring multiple novel isoforms (**Figure 6A**). We identified 481 DEGs (FDR < 5%, |log₂FC| > 2) between GABA and GLU neurons and 1,758 DEGs between neurons and OLIG (**Figure S18A-B, Table S11**). The OFC yielded 476 and 2,188 DEGs, respectively (**Figure S18C-D**), with a similar distribution of novel isoform content across DEGs (**Figure 6B**). Cross-region conservation was substantial, indicating conserved lineage-specific programs. Notably, a large proportion of DEGs harbored novel isoforms absent from GENCODE annotations, underscoring that gene-level counts underestimate the transcriptional complexity revealed by long-read sequencing. Functional enrichment highlighted neuronal pathways related to synaptic signaling, neurotransmitter transport, and calcium-dependent processes, while OLIG-enriched genes were dominated by myelination, lipid metabolism, and RNA processing pathways (**Figure S19, Tables S14-S15**).

**Figure 6.**
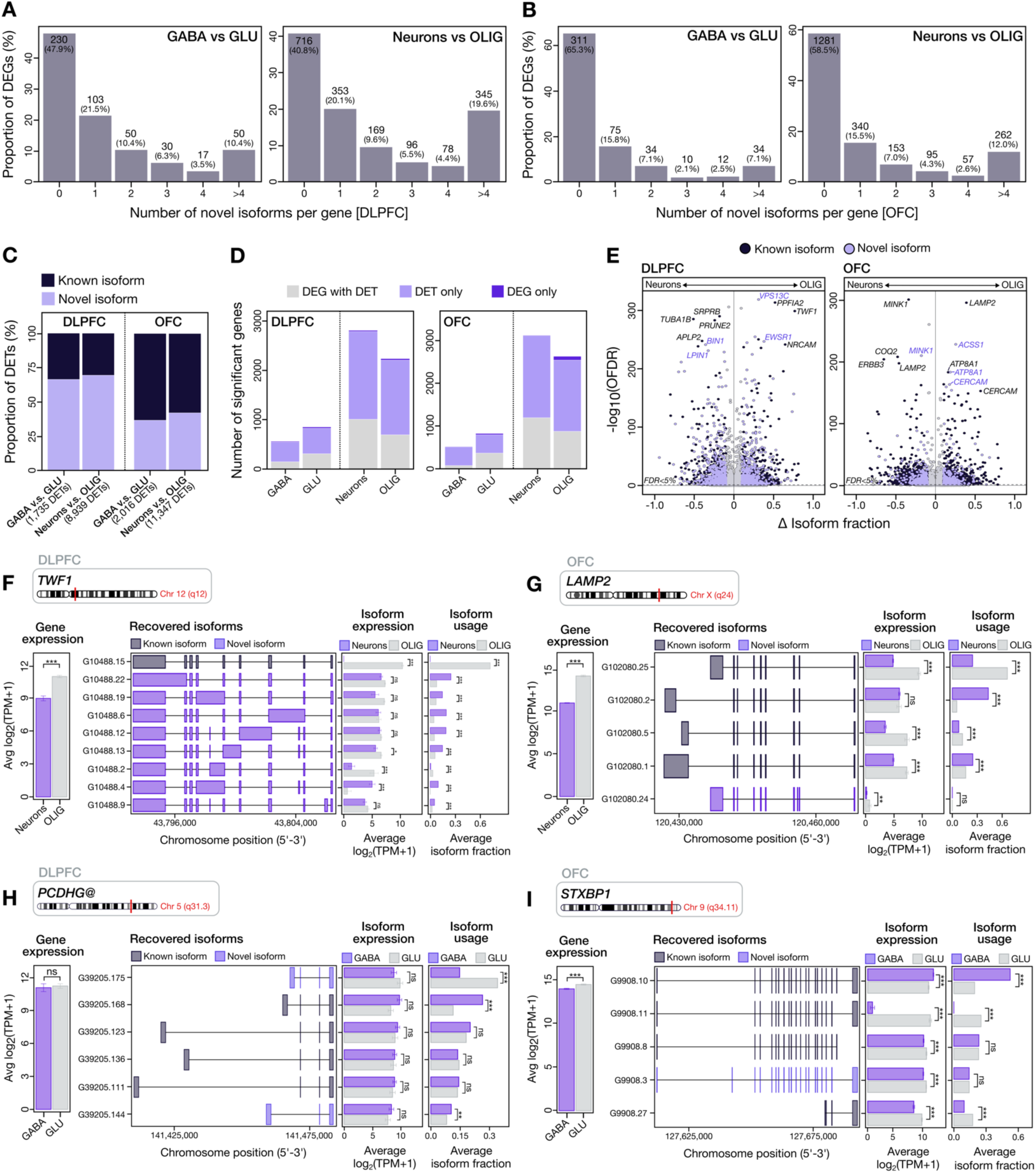
Isoform-level regulation defines cortical cell identity independently of differential gene expression. (**A-B**) Distribution of the number of novel isoforms per differentially expressed gene (DEG; FDR < 5%, |log₂FC| > 2) for GABA versus GLU and neurons versus OLIG in (**A**) DLPFC and (**B**) OFC. Bars indicate the proportion of DEGs harboring 0, 1, 2, 3, 4, or >4 novel isoforms; counts are shown above each bar. (**C**) Proportion of differentially expressed transcripts (DETs) classified as known or novel isoforms across all comparisons in DLPFC and OFC; the majority of DETs correspond to novel isoforms. (**D**) Concordance between differential gene expression (DGE) and differential transcript expression (DTE) per cell type. Stacked bars indicate genes significant in both analyses, DTE only, or DGE only (DTE summarized at the gene level). (**E**) Volcano plots of differential transcript usage (DTU) for GABA versus GLU and neurons versus OLIG in DLPFC and OFC. Each point represents one isoform, colored by novelty status (known, dark; novel, light). Significance threshold: FDR-adjusted q < 5% and |ΔIF| > 0.1. (**F-I**) Representative loci illustrating isoform-level regulation across cortical cell types. Shown are gene-level expression, recovered isoform structures (dark gray, known; purple, novel), average isoform expression, and isoform usage (fraction) per cell type for (**F**) *TWF1*, (**G**) *LAMP2*, (**H**) *PCDHG*, and (**I**) *STXBP1*.

DTE revealed that regulatory divergence is substantially more extensive at the isoform level. In the DLPFC, we identified 1,937 differentially expressed transcripts (DETs) across 1,242 genes between GABA and GLU neurons and 9,930 DETs across 4,060 genes between neurons and OLIG. The OFC showed a similar pattern, with 2,052 DETs across 1,299 genes and 11,651 DETs across 4,913 genes, respectively (**Figure 6C, Figure S18E-H, Table S12**). Approximately 59%-62% DETs corresponded to novel isoforms, indicating that the most differentially regulated transcripts across cortical lineages are largely absent from current reference annotations. Most DEGs contained at least one DET (**Figure 6D**). Critically, many DETs arose from genes not classified as DEGs, revealing a layer of regulatory divergence invisible to gene-centric analyses.

### Dominant isoform switching encodes cell type identity and exposes disease-relevant exonic sequence

We next performed differential transcript usage (DTU) to identify shifts in isoform proportions independent of total gene expression (FDR < 5%, Δisoform fraction > 10%). DTU was widespread across all comparisons and, again, was strongest between neurons and OLIG. In the DLPFC, 1,108 isoforms showed significant DTU between GABA and GLU neurons and 3,577 isoforms between neurons and OLIG (**Table S13**); the OFC identified 661 and 4,163 events, respectively (**Figure 6E**). DTU was highly concordant across regions (*r* = 0.47-0.54; **Figure S20A-C**), confirming reproducible cell type-specific isoform usage across independent cortical areas.

Integration of DGE, DTE, and DTU demonstrated that these regulatory layers are largely non-redundant. Most DTU events occurred independently of DGE, and only ∼9% of transcripts showed both DTE and DTU between GABA and GLU neurons, rising to 14-17% at the neuron-OLIG boundary (**Figure S20D-E**), consistent with progressively deeper transcriptome remodeling at neuronal-glial boundaries. Isoform composition is regulated independently of gene expression.

Multi-isoform rebalancing within single loci was extensive. In the DLPFC, 45 genes showed multiple DTU isoforms in the GABA-GLU comparison and 331 in the neuron-OLIG contrast; the OFC showed 28 and 367, respectively (**Figure S20D-E**). *TWF1*, an actin-binding regulator of cytoskeletal remodeling, showed both differential expression and DTU between neurons and OLIG, with one OLIG-enriched isoform and several neuron-specific novel isoforms (**Figure 6F**). *LAMP2* displayed elevated expression and multiple OLIG-enriched isoforms across both regions (**Figure 6G**). In contrast, gene cluster *PCDHG* showed comparable gene-level abundance between GABA and GLU neurons but marked differences in isoform usage, with a GLU-dominant isoform contributing ∼30% of total output versus ∼15% in GABA (**Figure 6H**). Similarly, *STXBP1* showed subtype-specific isoform selection despite minimal expression differences (**Figure 6I**), indicating that the functional output of disease-relevant synaptic genes is set at the isoform level, not the gene level.

Dominant isoform switching, where the major expressed transcript of a gene differs between cell types (|ΔIF| > 0.5, FDR < 5%), was most extensive between neurons and OLIG (DLPFC: 1,816 events; OFC: 1,547), with far fewer events between GABA and GLU (DLPFC: 317; OFC: 185) (**Figure 7A**). A representative example, *SLC5A6*, encoding the sodium-dependent multivitamin transporter, showed distinct dominant isoforms across GABA, GLU, and OLIG populations in both cortical regions, nearly all of which were novel and absent from current reference annotations (**Figure 7B**). Because *SLC5A6* regulates transport of pantothenate, biotin, and lipoate, these isoform switches may alter metabolic capacity across cortical cell types. Three disease-causing variants overlapped exons unique to these dominant isoforms, linking cell type-specific isoform switching to neurodevelopmental disease risk.

**Figure 7.**
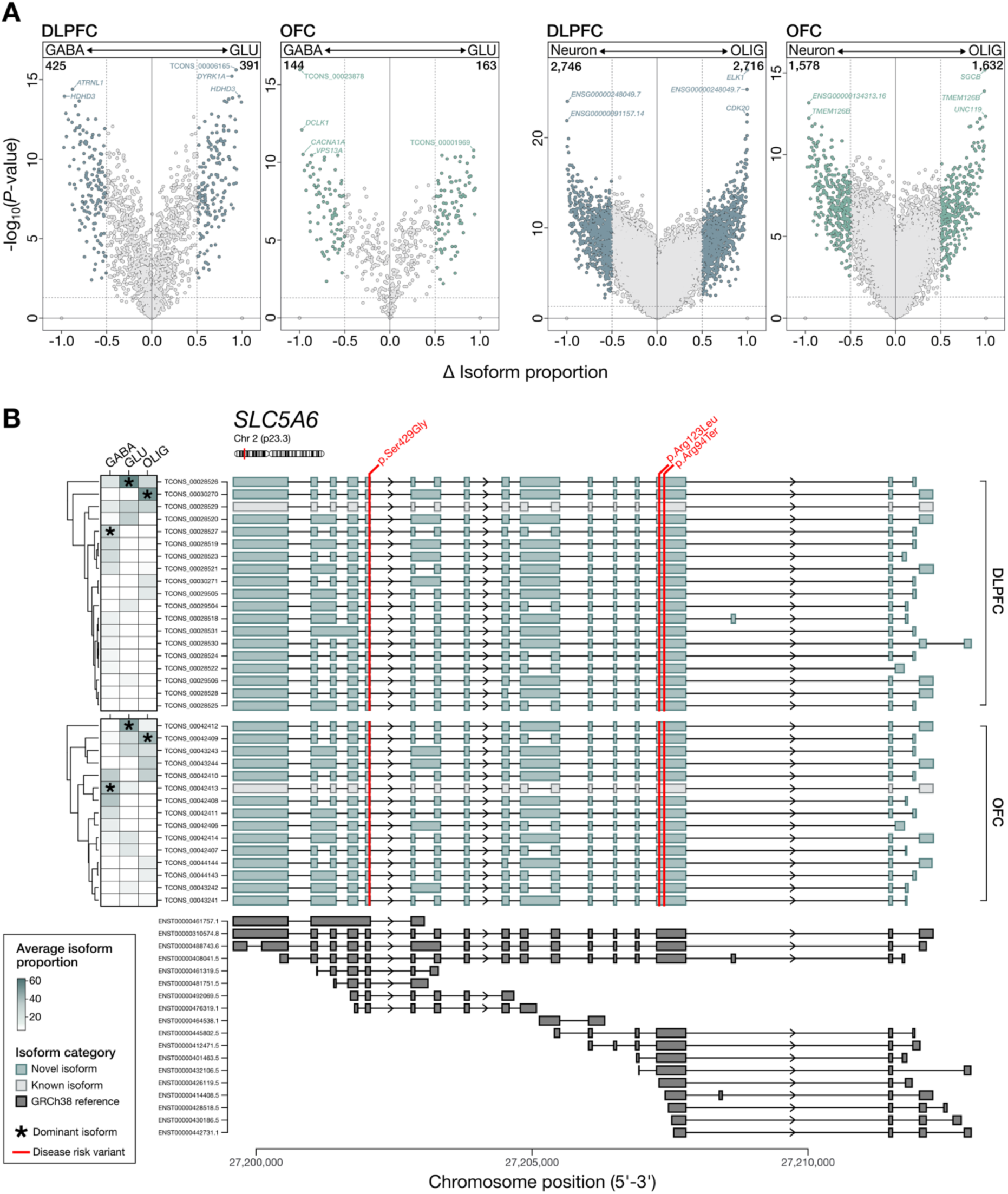
Dominant isoform switching encodes cortical cell type identity and intersects disease-causing variants. **(A)** Volcano plots of differential transcript usage (DTU) for GABA versus GLU neurons and neurons versus OLIG in DLPFC and OFC. x-axis, change in isoform fraction (ΔIF); y-axis, -log10 (p-value). Each point represents one isoform. Dashed lines indicate significance thresholds (|ΔIF| > 0.5 and *p* < 0.05). Numbers denote the total number of isoforms meeting these criteria in each comparison. **(B)** Isoform usage at *SLC5A6* across cortical cell types in DLPFC and OFC. Left, heatmap showing average isoform fraction per cell type, with asterisks marking dominant isoforms. Right, corresponding isoform structures, with novel isoforms (teal), known isoforms (gray), and GENCODE v38 annotations (dark gray). Red dashed lines indicate HGMD disease-causing variants mapping within exons unique to cell type-specific dominant isoforms.

### Isoform co-expression networks reveal cell type-structured regulatory architecture and converge on neurodevelopmental disease risk

Isoform co-expression network analysis resolved 11 modules in the DLPFC and 7 in the OFC, revealing a hierarchical, cell type-structured organization of transcript regulation with strongly conserved functional architecture across regions (**Figures S21-S22**; **Tables S17-S19**). Approximately 22-30% of all genes, and nearly a third of multi-isoform genes, distributed their transcripts across two or more co-expression modules (DLPFC: 30.4%; OFC: 21.9%) (**Figure S22**), indicating that isoforms from the same gene frequently participate in distinct regulatory programs that are obscured by gene-level analysis. Neuronally enriched modules showed the strongest and most reproducible enrichment for bona fide neurodevelopmental risk genes^38–40^ (**Figure S23**), and 38 autism linked genes containing novel isoforms directly intersected bona fide pathogenic variants within isoform-specific exonic regions, including recurrent risk genes *POGZ*, *CHD8*, *FOXP1*, *DYRK1A*, and *SMARCC2* (Figure 8, **Tables S20-S21**; *see **Supplementary Results** for full details*).These findings link previously unannotated transcript structures to genetic disease risk at single-exon resolution. These analyses were, however, restricted to a focused set of ASD-linked loci and co-expression networks; whether this principle generalizes more broadly is addressed in the following section.

**Figure 8.**
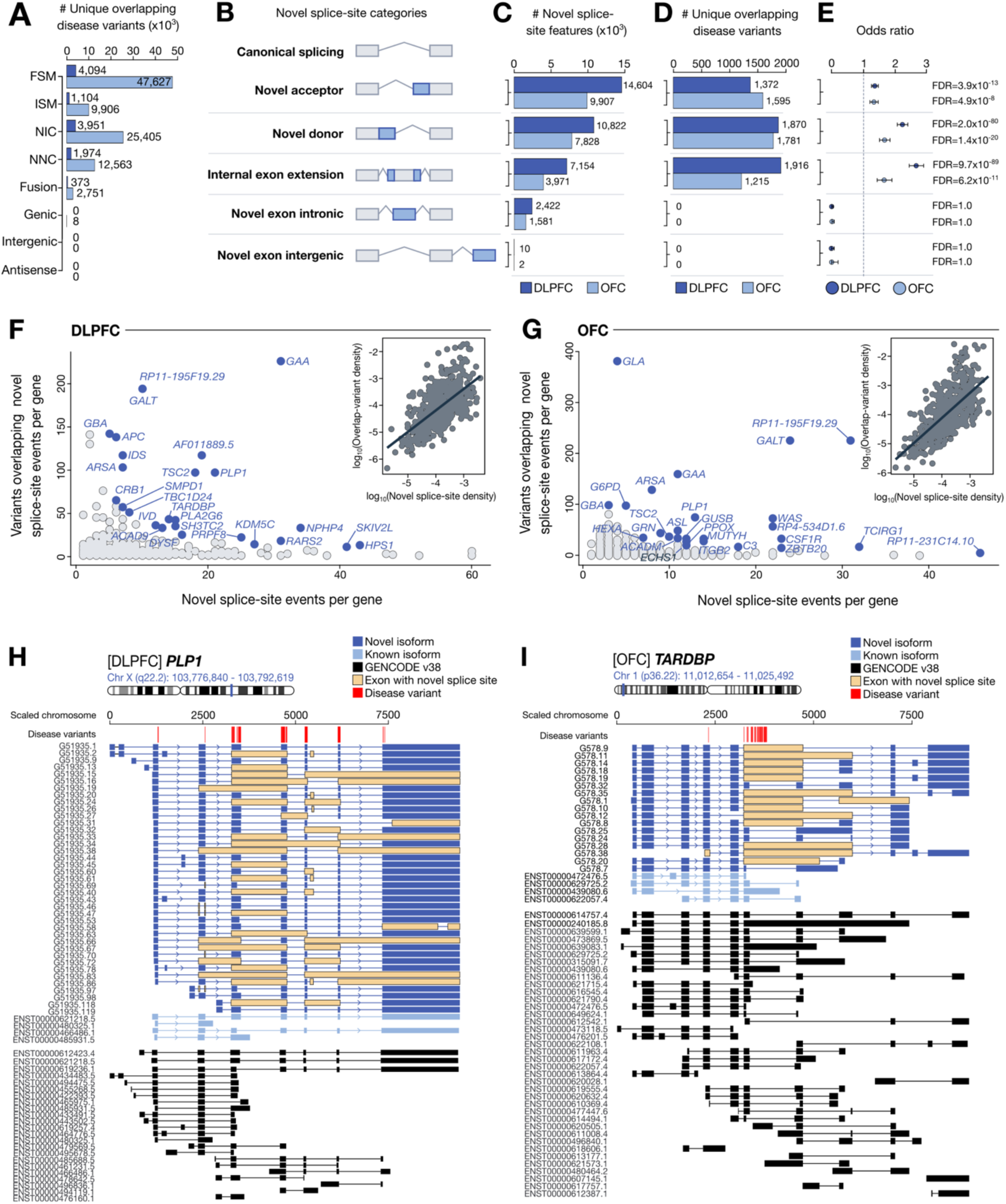
Novel splice-site architecture exposes disease-causing variants within previously unannotated exonic sequence. (**A**) Number of unique HGMD disease-causing variants (DM class) overlapping isoform exons across SQANTI3 structural categories in DLPFC and OFC. (**B**) Schematic of novel splice-site feature classes within NNC isoforms relative to canonical splice architecture, including novel acceptors, novel donors, internal exon extensions, and novel exons. (**C**) Number of novel splice-site features per class in DLPFC and OFC. (**D**) Number of unique HGMD disease-causing variants overlapping each novel splice-site class in DLPFC and OFC. (**E**) Enrichment of disease-causing variants within novel splice-site classes relative to annotated GENCODE v38 exons, shown as odds ratios with 95% confidence intervals and FDR-adjusted p-values. (**F-G**) Relationship between the number of novel splice-site events per gene and the number of overlapping HGMD variants per gene in (**F**) DLPFC and (**G**) OFC; insets show analyses normalized for gene length. Selected loci are labeled. (**H-I**) Locus plots for (**H**) *PLP1* in DLPFC and (**I**) *TARDBP* in OFC. Blue, novel isoforms; light blue, known isoforms; black, GENCODE v38 reference; orange shading, exons containing novel splice sites; red bars, HGMD disease-causing variants.

### Long-read isoform architecture exposes a hidden layer of disease-relevant exonic sequence in the human cortex

The ultimate test of this isoform atlas is whether it changes how we interpret genetic disease risk. We addressed this by intersecting our full-length isoform annotations with high-confidence disease-causing mutations (DM class) from the Human Gene Mutation Database (HGMD)^41^. Pathogenic variants were distributed across the transcriptome, spanning FSM, NIC, and NNC isoforms, confirming that disease-associated variation is embedded within both reference-consistent and previously unannotated transcript structures (**Figure 8A**).

We next reclassified NNC isoforms by splice-site architecture, including novel acceptors, novel donors, internal exon extensions, novel intronic exons, and novel intergenic exons (**Figure 8B**). Across both cortical regions, the predominant classes were splice-boundary alterations rather than de novo exon creation. Novel acceptors represented the largest category (DLPFC: 14,604 [41.7%]; OFC: 9,907 [42.5%]), followed by novel donors (10,822 [30.9%]; OFC: 7,828 [33.6%]) and internal exon extensions (7,154 [20.4%]; OFC: 3,971 [17.1%]), whereas fully novel intronic and intergenic exons were comparatively rare (**Figure 8C**). Splice-site plasticity, not de novo exon creation, is the primary mechanism of transcript novelty in the adult human cortex.

Intersecting these features with HGMD disease variants revealed a strikingly selective pattern: pathogenic variation was strongly concentrated within splice-boundary classes, particularly novel donors, novel acceptors, and internal exon extensions (**Figure 8D**). In the DLPFC, 1,916 variants overlapped internal exon extensions, 1,870 overlapped novel donors, and 1,372 overlapped novel acceptors; in the OFC, the corresponding counts were 1,215, 1,781, and 1,595, respectively (**Figure 8D**). Enrichment testing against annotated GENCODE exons confirmed significant overrepresentation of disease-associated variants within these splice-boundary classes in both cortical regions, with the strongest effects observed for internal exon extensions (DLPFC OR = 2.67; OFC OR = 1.65) and novel donor events (DLPFC OR = 2.22; OFC OR = 1.66) (**Figure 8E**). By contrast, intronic and intergenic novel exons showed no enrichment. Disease variants clustered non-randomly near splice junctions: in DLPFC, 1,331 variants fell within ±50 bp of novel splice sites (∼12% within ±5 bp); OFC showed a near-identical pattern (1,029 variants; ∼11% within ±5 bp) (**Figure S24A**). Genes with denser novel splice-site architectures showed significantly greater disease variant overlap in both regions (**Figure 8F-G**), including *GAA*, *PLP1*, *TSC2*, *GBA*, and *TARDBP* in DLPFC, and *GALT*, *GAA*, *GLA*, *CSF1R*, *ARSA*, and *PLP1* in OFC. This relationship held after normalization for gene length (**Figure 8F-G**, insets), confirming that complex isoform architecture predicts hidden disease variant burden (**Table S22**). Importantly, these genes also showed high cell type specificity of expression, indicating that disease-associated splice architecture is often embedded within genes preferentially expressed in specific cortical lineages rather than uniformly across cell types (**Figure S24B-C**).

Representative loci illustrated how these hidden transcript structures reshape disease interpretation. In *PLP1*, multiple disease-associated variants mapped within coding exonic regions newly exposed by splice-boundary extension that were absent from canonical reference annotations (**Figure 8H**). Similarly, in *TARDBP*, novel donor and acceptor usage created previously unannotated coding structures that directly overlapped pathogenic variants linked to neurodevelopmental phenotypes (**Figure 8I**). These examples illustrate that variant interpretation based solely on canonical transcript models can misclassify pathogenic sequence as intronic or non-coding when relevant isoform structures are absent from the reference. Long-read transcriptomics of defined cell populations exposes a layer of pathogenic variation invisible to canonical gene models and supports a model in which cortical cell identity and disease risk are shaped at the isoform level.

## DISCUSSION

In the adult human cortex, cell identity is defined not only by which genes are expressed, but by which isoforms of those genes are preferentially used. The full-length isoform atlas generated here demonstrates this across five major cortical lineages, revealing more than 220,000 unique isoforms, ∼35-56% of which lack representation in current GENCODE annotations across the adult DLPFC and OFC. This scale of unannotated transcript diversity, resolved at the level of purified cell populations, demonstrates that the human brain transcriptome remains substantially incomplete, and that isoform-level resolution is not a refinement but a requirement for understanding cortical gene regulation. Building on prior long-read studies of the human brain, our cell type-resolved approach uncovers regulatory features, including lineage-restricted splice-junction usage, pervasive isoform switching, and widespread intron retention coupled to NMD that are largely inaccessible to short-read or bulk-tissue profiling. Novel isoforms were consistently longer, contained more exons, and were predominantly protein-coding, with strong orthogonal support from ATAC-seq^30–33^, CAGE^34–35^, 3′-seq^36^, independent long-read RNA-seq datasets^21,22,36,37^, and evolutionary constraint profiles comparable to fully annotated transcripts. These properties collectively argue that the unannotated isoforms catalogued here are functional components of cortical gene regulation whose omission from reference annotations can substantially limit interpretation of both cell identity and disease risk.

The most prominent finding is that glial populations, not neurons, harbor greater isoform diversity. Across both cortical regions, OLIG and MG consistently surpassed GABA and GLU neurons in isoform count, transcript novelty, and the proportion of cell type-restricted isoforms. This finding inverts the neuron-centric framing that has dominated brain transcriptomics and reflects a fundamental limitation of prior bulk-tissue studies^42–44^, in which glial signals are diluted by neuronal abundance. By profiling purified populations, we show that glial lineages harbor some of the most elaborately spliced transcriptomes in the adult cortex, extending emerging observations from mouse and developmental studies^19–21^. The predominance of novel, multi-exonic isoforms in OLIG aligns with the substantial biosynthetic and cytoskeletal demands of myelination^45^, while MG isoform diversity is concentrated in genes governing immune signaling and endolysosomal trafficking, consistent with the functional adaptability required of resident brain immune cells^46^. Unlike neurons, which achieve functional specialization through discrete cellular subtypes, microglia diversify primarily through transcriptional state transitions, including surveillance, phagocytic, and disease-associated states, requiring rapid adaptation to signals such as infection, ischemia, and synaptic activity^47^. The breadth of MG isoform diversity identified here may therefore reflect the transcriptomic repertoire underlying this state-dependent plasticity across heterogeneous brain environments. Moreover, the enrichment of extended coding regions and divergent exon combinations in glial isoforms raises the possibility of altered protein domain composition with direct functional consequences for myelination, phagocytosis, and inflammatory signaling, effects invisible at the gene level^48^. Saturation analyses and five-layer orthogonal validation confirm that these patterns reflect biology rather than technical artifact. These results reframe glial populations as major contributors to cortical isoform complexity and establish a foundation for understanding how glial splice dysregulation may contribute to neurological and neuropsychiatric disease.

Intron retention was the dominant AS event type across all five cortical lineages and was tightly and reproducibly coupled to nonsense-mediated decay. Prior long-read studies of fetal cortex^21^, bulk adult brain^22^, and mouse hippocampus^15^ have each highlighted the prominence of RI, but our dataset extends these observations to purified human cortical populations and directly links RI to degradation programs at scale. Because profiling was performed on isolated nuclei, where unspliced and partially spliced transcripts are naturally enriched^49^, the overall prevalence of RI may be modestly higher than in whole-cell preparations. However, several observations argue against a purely technical explanation. The RI fraction observed here (∼31%) is only modestly higher than prior long-read studies of bulk human brain (∼23-27%)^19,37^. RI isoforms were flanked by canonical splice sites, enriched for nonsense-mediated decay architecture, and showed lower expression than non-NMD transcripts. Established loci such as *MEG3* showed concordant patterns with prior reports^19^. RI events displayed robust cell type specificity and reproducibility across donors and cortical regions. And critically, >96% of our novel isoforms acheive SQANTI3 Tier 2 orthogonal support, arguing strongly against artifactual reconstruction. Indeed, the balance between NMD-targeted RI isoforms and the substantial subset that escapes predicted decay and persists at stable expression levels suggests that intron retention operates as a regulatory mechanism to tune transcript availability in a cell type-specific manner^50–52^. Loci such as *SRSF11* and *DDX3X* exemplify this flexibility, simultaneously deploying RI for transcript clearance and functional isoform diversification across neuronal and glial lineages^53^. These results support an emerging view of intron retention as a substantive regulatory axis in the brain^52,53^, with potential roles in nuclear retention, cytoplasmic export, and the timing of transcript availability for translation^54^.

Isoform switching was widespread across all lineage comparisons and occurred largely independently of gene-level expression changes, demonstrating that transcript usage is an independent axis of cortical cell identity rather than a downstream consequence of differential gene expression. While prior long-read studies have documented isoform switching during corticogenesis and across bulk brain regions^5,22,55^, our data define a more extensive and cell type-resolved landscape in the mature human cortex. The decoupling of DTU from DGE was most pronounced at the neuron-OLIG boundary, where isoform-level divergence far exceeds gene-level differences, indicating that transcript selection contributes to regulatory outcomes distinct from those captured by expression abundance alone^56,57^. Representative loci *SLC5A6*, *TWF1*, *LAMP2*, *PCDHG*, and *STXBP1* each display distinct dominant isoforms despite stable or modestly changing gene expression, illustrating how functional diversification is encoded at the isoform rather than the gene level. Nearly all dominant isoforms involved in these switches were novel and absent from current annotations, meaning the functionally dominant transcript for a given gene in a given cell type is, in most cases, not captured by reference-based analyses^58^. Extended or alternative UTR architectures present in many switched isoforms introduce regulatory sequences that may influence mRNA stability, localization, or translational efficiency^59,60^, providing a mechanistic basis for cell type-specific isoform selection that operates independently of transcriptional control. The direct overlap of disease-causing variants with exons unique to dominant isoforms, illustrated by *SLC5A6* and confirmed more broadly across the HGMD intersection, further demonstrates that isoform switching is not only a feature of cell identity, but a determinant of how genetic risk is read out in specific cortical lineages.

Isoform-level co-expression analysis extended these findings from pairwise comparisons to network-level organization, revealing that transcript regulation in the adult cortex is structured along cell type- and region-specific axes that are invisible at the gene level. Neuronal and oligodendrocyte modules showed reproducible functional coherence (i.e., synaptic signaling and cytoskeletal organization in neurons; RNA processing, ribonucleoprotein assembly, and mRNA metabolism in OLIG) consistent with the known demands of these lineages and with prior evidence that transcript-level networks capture regulatory programs missed by gene-level co-expression^56,61^. Approximately 22-30% of all genes, and nearly half of multi-isoform genes, distributed their transcripts across two or more co-expression modules, indicating that isoforms from the same gene frequently participate in functionally distinct regulatory programs^62,63^. This architecture also converged on neurodevelopmental disease risk: neuronally enriched modules showed the strongest enrichment for autism-associated genes, and high-confidence loci such as *POGZ*, *CHD8*, *FOXP1*, *DYRK1A*, and *SMARCC2* harbored numerous novel isoforms directly intersecting pathogenic variants, consistent with recent evidence that isoform-resolved annotation substantially improves interpretation of pathogenic variation in neurodevelopmental disorders^64,65^. Together, these findings support isoform-level organization as a fundamental regulatory feature of cortical biology and suggest that alternative splicing provides an additional layer through which disease-associated variants exert cell type-specific effects.

More broadly, the intersection of novel splice-site architecture with HGMD disease variants demonstrates that pathogenic variation is selectively concentrated within the splice-boundary classes that drive transcript novelty in the cortex: novel donors, novel acceptors, and internal exon extensions. This enrichment was significant in both regions, strongest for internal exon extensions and novel donor events, and persisted after normalization for gene length, confirming that complex isoform architecture at specific loci, not gene size, predicts hidden disease variant burden. In *PLP1* and *TARDBP*, disease-associated variants mapped within coding exonic sequence exposed by splice-boundary extension that is entirely absent from canonical annotations. In *PLP1*, splice-site variants overlapping the exon boundaries identified here have been shown in independent minigene and patient-derived cell assays to alter exon inclusion and produce aberrant transcripts associated with Pelizaeus-Merzbacher disease^66,67^. In *TARDBP*, deep intronic and exon-boundary variants have been functionally validated to activate cryptic splice sites and generate truncated or aberrant TDP-43 isoforms in amyotrophic lateral sclerosis and frontotemporal dementia patient samples^68,69^. The convergence of our isoform-resolved mapping with these experimentally characterized splice events supports the interpretation that novel transcript structures identified here define functionally relevant exonic territory and that variants falling within them warrant experimental prioritization. Systematic functional validation of the broader variant-isoform intersections identified here, through splicing reporter assays, patient-derived iPSC models, or CRISPR perturbation, remains an important priority for future work.

Together, these findings argue that cortical cell identity and disease risk are written into transcript structure, not just gene-level expression. By pairing FANS-based long-read sequencing with extensive short-read validation across two cortical regions and five lineages, we generated a resource that substantially expands current transcript models, repositions glial populations as the major contributors to cortical isoform complexity, and directly links novel transcript structures to neurodevelopmental genetic risk. As single-cell long-read sequencing matures and proteomic and epigenomic integration becomes tractable, the principles defined here, that isoform selection is a primary axis of cortical cell identity, and that disease risk is embedded in transcript structures absent from canonical annotation, will anchor future mechanistic and translational studies of the human brain. The full atlas, including isoform structures, expression, and disease-variant intersections, is available through an interactive web application (https://andyyang-isoseq.share.connect.posit.cloud/).

### Limitations of the current study

This study has notable limitations. First, we performed long-read sequencing across a modest sample size: four biological replicates in the OFC and three in the DLPFC, profiling one sample per major cell type per donor (total n=12 OFC; n=9 DLPFC). While this scale is consistent with current long-read studies^19–22^ and our major findings were supported by strong cross-donor consistency and orthogonal short-read validation, the limited number of donors constrains population-level inference and reduces power to detect rarer or more subtle sources of inter-individual variation. In addition, astrocytes and microglia were profiled only in the OFC. While oligodendrocytes harbored the greatest isoform diversity of any cell type with replication across both regions, the corresponding claims for astrocytes and microglia rest on a single cortical region and should be considered provisional until cross-region replication is performed. Confirming the breadth of glial isoform diversity in additional cortical areas, and in further glial subtypes, is an important next step. Second, FANS resolved major cortical lineages but not finer neuronal, glial, or activation-state subtypes; therefore, the isoform diversity reported here may represent a conservative estimate of subtype-specific transcript complexity. Third, isoform discovery remained unsaturated with sequencing depth, indicating that rare and lowly expressed transcripts are still underrepresented and that the observed median of two isoforms per gene likely underestimates true per-cell transcript diversity. Fourth, although most novel isoforms were predicted to be protein-coding and showed strong orthogonal multi-modal validation, direct proteomic confirmation of translation and protein function was not performed. Fifth, enrichment of disease-causing variants within novel splice features was statistically robust but remains correlative, and experimental validation will be required to establish direct causal effects on splicing, isoform usage, and cellular phenotypes. Finally, disease-variant analyses are also limited by the ascertainment and annotation biases of existing databases such as HGMD, which are enriched for known coding and clinically studied loci.

## Supporting information

Supplemental Results

## RESOURCE AVAILABILITY

### Lead Contact

Requests for further information should be directed to and will be fulfilled by the lead contact, Michael S. Breen.

### Materials Availability

This study did not generate new, unique reagents.

### Data and Code Availability

- Raw PacBio Iso-Seq data are available at the NCBI Gene Expression Omnibus under the accession number GSE33XYZ (will be made public following publication). UCSC genome browser tracks of our processed Iso-Seq data together with a visual database of cortical isoforms are available through the Zenodo DOI: 10.5281/zenodo.20301473 and https://github.com/AndyYaj/cortical-FANS-IsoSeq/blob/main/cell_type_resolved_isoforms.zip. To maximize accessibility and utility, we developed an interactive R Shiny application to enable exploration of isoform structures and splicing events across all cortical cell types and regions generated in this study: https://andyyang-isoseq.share.connect.posit.cloud/
- All code used for data processing, analysis, and figure generation is publicly available at: https://github.com/AndyYaj/cortical-FANS-IsoSeq
- Any additional information required to reanalyze the data reported in this paper is available from the lead contact upon request.

## ACKNOWLEDGEMENTS

We sincerely thank the families of the brain donors for their invaluable contributions to this research. This work was supported by grants to MSB (5R01AG087324-02) and SD (NIH R01DA04324, R01MH122590, VA Merit Award I01BX00664), the Samuel A. Seaver Foundation (MSB), and philanthropic contributions to the Breen Laboratory. Computational analyses were performed in part using the MINERVA High-Performance Computing System at the Icahn School of Medicine at Mount Sinai.

## AUTHOR CONTRIBUTIONS

Dr. Breen had full access to all data in the study and takes responsibility for the integrity of the data and the accuracy of the analyses. Author contributions are as follows: study concept and design (M.S.B, S.D.), data acquisition (M.S.B, S.D., A.K., R.V, Y.H.), formal methodology and analysis (A.Y., M.S.B), visualization (A.Y.), interpretation (A.Y., M.S.B, S.D., J.H.), manuscript drafting (A.Y., M.S.B) and manuscript revision for intellectual content by all authors. Statistical analyses were performed by A.Y. and M.S.B. Funding, administrative, technical, and material support were provided by M.S.B, Y.H., S.D. Supervision was provided by S.D. and M.S.B.

## METHODS

### Study Design

This study generated a cell type-resolved isoform atlas of the adult human cortex by integrating PacBio Iso-Seq long-read sequencing with multiple FANS-derived Illumina RNA-seq cohorts. Full-length transcriptomes were profiled across the dorsolateral prefrontal cortex (DLPFC; Brodmann area 9/46) and the orbitofrontal cortex (OFC; Brodmann area 11), using fluorescence-activated nuclei sorting (FANS) to isolate five major cortical lineages, including MGE-derived GABAergic interneurons, glutamatergic neurons, oligodendrocytes, astrocytes, and microglia. Complementary short-read RNA-seq datasets derived from matched FANS protocols provided orthogonal splice-junction support and quantitative transcript abundance estimates, enabling accurate isoform reconstruction and validation across regions and cell types. Because long-read and short-read cohorts were processed in independent experimental batches, all primary analyses were performed within-region; cross-region comparisons were used exclusively as orthogonal validation of shared biological patterns.

### Postmortem Human Brain Tissue

Postmortem specimens were obtained from a brain bank curated by Dr. Yasmin Hurd at the Icahn School of Medicine at Mount Sinai under institutionally approved protocols, with informed consent from next of kin (**Table S1**). Brains were recovered at autopsy within 24 hours of death, as previously described^70^, and tissue pH values fell within the accepted high-quality range (6.4-6.8). Specimens were coronally sectioned, flash-frozen, and stored at −80°C until dissection. The cohort comprised Caucasian individuals of confirmed ancestry by ancestry-informative marker analysis.

### FANS-derived Long-read Iso-Seq Datasets

Long-read transcriptomes were generated from FANS-purified nuclei isolated from two cortical regions: DLPFC (GABA, GLU, and OLIG nuclei; n = 3 donors) and OFC (GABA, GLU, OLIG, AST, and MG nuclei; n = 4 donors). AST and MG were profiled exclusively in the OFC cohort, where all five lineages were represented, providing an internally controlled framework for glial-neuronal comparisons within that region. All primary cell type comparisons were performed within-region; cross-region analyses using OLIG, the only glial lineage profiled in both regions, served to confirm the reproducibility of glial isoform diversity findings across independent cohorts. All donors were neurologically and psychiatrically unremarkable adults with no documented neurological, neurodevelopmental, or major psychiatric diagnoses, negative toxicology screens, and non-neurological, non-suicide causes of death.

### FANS-derived Short-Read RNA-Seq Datasets

Three independent FANS-derived Illumina short-read RNA-seq cohorts were integrated to provide splice-junction validation and quantitative backbone for all differential expression and transcript usage analyses. Together, these cohorts, comprising up to 9 donors per cell type in DLPFC and 58 donors in OFC, constitute the primary statistical engine for the DGE, DTE, and DTU analyses reported here.

- **DLPFC cohort (GABA, GLU, OLIG).** Twenty-seven paired-end (125 bp) RNA-seq libraries generated from DLPFC nuclei (n = 9 donors per cell type) were obtained from a previously published dataset (Synapse ID: syn12034263)^23,27^. Nuclei were isolated using the same NeuN/SOX6/SOX10 FANS strategy applied in the current study, providing directly comparable cell type-specific splice-junction support for GABA, GLU, and OLIG populations. All donors were neurologically and psychiatrically unaffected adults with no history of major neurodevelopmental, neurological, or psychiatric disorders, negative toxicology screens, and non-neurological, non-suicide causes of death.
- **OFC cohort (GABA, GLU, OLIG).** One hundred and eighteen short-read RNA-seq libraries from 58 OFC donors spanning GABAergic, glutamatergic, and oligodendrocyte nuclei were incorporated^28^. Nuclei were isolated using FANS approaches matching those applied to the Iso-Seq samples. This cohort provides the primary quantitative depth for validating splice junctions and performing differential analyses in OFC neuronal and oligodendrocyte lineages. The cohort comprised individuals who died of apparent heroin overdose (n = 30) and neurologically and psychiatrically unaffected controls (n = 28; confirmed by self-report and ancestry-informative marker analysis).
- **Multi-region NeuN⁺/NeuN⁻ cohort (five cortical areas).** To extend splice-junction validation to AST and MG isoforms, which lack matched cell type-pure short-read data, we integrated 100 RNA-seq libraries from 10 donors sampled across five cortical regions (BM10, BM17, BM22, BM36, BM44; Synapse ID: syn25716684)^29^. This dataset comprises NeuN⁺ neuronal and NeuN⁻ non-neuronal nuclei; although not cell type-pure, the deeply sequenced NeuN⁻ fraction is enriched for glial populations and provides robust splice-junction evidence for AST- and MG-enriched exons. This cohort comprised neurologically and psychiatrically unaffected control individuals, consistent with the long-read and DLPFC short-read cohorts.

### Isolation of FANS-Purified Cortical Nuclei

Cell type-specific cortical nuclei were isolated using an optimized FANS workflow adapted from our previously published protocols and related long-read and chromatin studies of human postmortem brain tissue^23,71–73^. Briefly, ∼300-750 mg of frozen cortical tissue was homogenized in ice-cold lysis buffer (320 mM sucrose, 5 mM CaCl₂, 3 mM Mg(Ac)₂, 0.1 mM EDTA, 0.1% Triton X-100, 1 mM DTT, RNase inhibitor, and 10 mM Tris-HCl, pH 8.0) and underlaid with a high-sucrose cushion (1.8 M sucrose, 3 mM Mg(Ac)₂, 1 mM DTT, RNase inhibitor, and 10 mM Tris-HCl, pH 8.0). Homogenates were centrifuged for 1 h at 98,600 × g (24,000 rpm, SW41Ti rotor), after which nuclear pellets were gently resuspended in antibody-incubation buffer (0.5% BSA, 3 mM MgCl₂, RNase inhibitor, 10 mM Tris-HCl, pH 8.0).

Nuclei were incubated with primary antibodies for 1-1.5 hours at 4°C, followed by secondary antibody incubation where required. The sorting strategy used NeuN (RBFOX3) to identify neuronal nuclei, SOX6 to distinguish MGE-derived GABAergic from glutamatergic neurons within the NeuN⁺ fraction, SOX10 to mark oligodendrocyte lineage nuclei, and IRF5 to separate microglia from astrocytes within the NeuN⁻/SOX10⁻ fraction. The antibody panel consisted of PE-conjugated anti-NeuN (Millipore, FCMAB317PE, 1:1000), guinea pig polyclonal anti-SOX6 (1:1500) detected with Alexa Fluor 647-conjugated donkey anti-guinea pig IgG (Jackson ImmunoResearch, 1:1500), goat polyclonal anti-SOX10 (R&D Systems, AF2864, 1:300) detected with Alexa Fluor 488-conjugated donkey anti-goat IgG (Life Technologies, A21447, 1:1500), and Alexa Fluor 488-conjugated anti-IRF5 (R&D Systems, IC4508G, 1:200). All nuclei were counterstained with DAPI to exclude debris and doublets.

FANS was performed on a BD FACSAria instrument. Sequential gating on DAPI⁺ singlets followed by NeuN, SOX6, SOX10, and IRF5 signal patterns yielded five purified populations: GLU (NeuN⁺/SOX6⁻), GABA (NeuN⁺/SOX6⁺), OLIG (NeuN⁻/SOX10⁺), MG (NeuN⁻/SOX10⁻/IRF5⁺), and AST (NeuN⁻/SOX10⁻/IRF5⁻). Approximately 40,000 nuclei per population were recovered per sample. Sorted nuclei were collected directly into PicoPure lysis buffer to preserve RNA integrity for downstream Iso-Seq library preparation.

### RNA Extraction and Quality Metrics

Total RNA was prepared from sorted nuclei using the PicoPure RNA Isolation Kit (Thermo Fisher Scientific) and quantified with Qubit RNA HS assays. RNA integrity was assessed with the Agilent RNA Pico Kit. Although bulk tissue from all donors exhibited high RNA quality (pH 6.4-6.8; RIN ≥ 6), RIN values measured after nuclei isolation were lower (DLPFC: 5.4-7.9; OFC: 3.5-6.6), reflecting reduced cytoplasmic rRNA content in nuclear fractions rather than RNA degradation, a well-established property of nuclear RNA preparations in which RIN is not a reliable quality metric^74^. RNA input amounts ranged from 33-240 ng across DLPFC populations and 17-280 ng across OFC populations, with neuronal fractions generally yielding the highest mass.

### Full-Length cDNA Synthesis and Quality Assessment

Full-length cDNA was synthesized using the PacBio Iso-Seq Express kit following manufacturer recommendations for low-input RNA. First-strand synthesis and template switching were performed on 30-240 ng of total RNA. cDNA length distributions were monitored by electrophoretic profiling prior to amplification to confirm representation of full-length transcripts. An initial 10-12 cycles of PCR yielded 15-77 ng of cDNA (size distribution ∼2.7-3.2 kb); a second amplification step produced 80-410 ng sufficient for SMRTbell construction. At each checkpoint, cDNA integrity was assessed by Femto Pulse or Bioanalyzer, and samples progressed only if they displayed clean full-length peaks without evidence of degradation or over-amplification.

### Long-Read Library Construction and Sequencing

SMRTbell libraries were prepared using the SMRTbell Express Template Prep Kit 2.0 (PacBio). Amplified cDNA (200-500 ng) underwent DNA damage repair, end polishing, A-tailing, and SMRTbell adapter ligation, followed by Exonuclease III/VII treatment and AMPure PB bead purification. Final libraries exhibited tight size distributions (∼2.8-3.2 kb) with no adapter-dimer contamination as assessed by High Sensitivity DNA or Femto Pulse analysis. Sequencing was performed on the PacBio Sequel II platform using Sequel II Binding Kit 2.2 and Sequencing Kit 2.0, with libraries annealed to sequencing primer v4. Each library was sequenced on a dedicated 8M SMRT Cell with movie times exceeding 24 hours, typically yielding 250-350 Gb of raw data and ∼3-4 million CCS reads per sample.

### Long-Read Data Processing

Raw PacBio Iso-Seq data (BAM format) were processed using the Iso-Seq3 pipeline (v3.8.2; https://github.com/ylipacbio/IsoSeq3). CCS reads were called using Iso-Seq3 CCS (v3.1) with default parameters^75^, which performs iterative multiple-sequence alignment across subreads for error correction. Five-prime and three-prime cDNA primers and SMRT adapters were removed using Iso-Seq3 Lima (v2.7.1) to generate full-length (FL) reads. PolyA tails were trimmed and artificial concatemers removed using Iso-Seq3 Refine to yield full-length non-concatemer (FLNC) reads. FLNC reads were clustered and polished using Iso-Seq3 Cluster/Polish, which applies hierarchical nlog(n) alignment and iterative cluster merging^76^ followed by multiple-alignment-based error correction. Polished reads were mapped to the human GRCh38/GENCODE v38 reference genome using *minimap2* (v2.24) with default parameters^77^ and resulting BAM files were sorted for downstream processing.

### Short-Read Data Processing

Raw short-read RNA-seq data were processed using an established pipeline (https://github.com/CommonMindConsortium/RAPiD-nf/)^78^ and aligned to the GRCh38 primary assembly with GENCODE v38 annotations using STAR (v2.7)^79^.

### Transcriptome Reconstruction

Polished, mapped long-read data were processed using the Transcriptome Annotation by Modular Algorithms (TAMA) pipeline (https://github.com/jkimlab/TAMA)^80^. Redundant reads were removed by genomic location using TAMA Collapse (parameters: -d merge_dup -sj sj_priority -x no_cap -log log_off -lde 1 -sjt 20 -a 100 -z 100). Cell type-specific transcriptomes were generated by merging samples within each cell type using TAMA Merge (-d merge_dup). Three TAMA Go error-correction scripts were then applied sequentially: tama_remove_single_read_models_levels.py to filter transcript models with low read support; tama_remove_polya_models_levels.py to remove single-exon models with 3′ genomic polyA stretches; and tama_remove_fragment_models.py to remove apparent transcript fragments. Cell type-specific transcriptomes were merged within each cortical region using TAMA Merge (parameter: ‘-d merge_dup’), and the same error-correction pipeline was reapplied to the region-level annotations.

### Isoform Classification and Quality Filtering

Transcriptomes were processed using SQANTI3^81–83^ to generate transcript- and junction-level quality metrics, correct indels, and classify full-length isoforms relative to GENCODE v38. External validation resources included FANTOM5 CAGE peaks (human, hg38)^34–35^, canonical polyA motifs and 3’-seq^36^, the Intropolis splice-junction compendium^84^, STAR-derived junction files from matched short-read RNA-seq, and full-length non-chimeric (FLNC) read counts. Indel-corrected transcript models were generated by realignment of full-length reads to GRCh38, and all downstream analyses were restricted to high-confidence isoforms.

Transcripts were retained only if all splice junctions were supported by at least two short-read junctions and the transcript contained two or more exons. Transcripts were flagged and removed if they showed evidence of poly(A) intra-priming, defined as: (i) location >25 bp from an annotated transcription termination site (TTS), (ii) >60% adenine content downstream of the 3′ end, and (iii) no overlap with the polyA site database or curated SQANTI3 poly(A) motifs (https://github.com/ConesaLab/SQANTI3). Transcripts with unreliable 5′ ends were also removed if they were located >25 bp from an annotated transcription start site (TSS) and lacked overlap with FANTOM5 CAGE peaks. Junctions flagged as noncanonical or potentially derived from reverse-transcriptase switching were excluded using both the Intropolis compendium and matched short-read STAR splice-junction support.

High-confidence isoforms were then classified as full-splice match (FSM), incomplete-splice match (ISM), novel in catalog (NIC), novel not in catalog (NNC), or other (including fusion, antisense, intergenic, and genic transcripts).

### Validation of Novel Isoform Transcript Boundaries

Analyses were performed using filtered transcript annotations generated by SQANTI3 (RulesFilter output). Transcript 5′ boundaries were validated by overlap with FANTOM5 CAGE peaks^34,35^ within ±200 bp of annotated TSS positions, and by chromatin accessibility signal from four independent human brain ATAC-seq datasets^30–33^ within −100 bp upstream to +1 kb downstream of the TSS. Transcript 3′ boundaries were validated using polyadenylation site annotations from the PolyASite database^36^ (Homo sapiens v2, GRCh38.96) and the presence of canonical polyadenylation sequence motifs identified by SQANTI3 within 50 bp upstream of the TTS. Isoforms were stratified into support tiers: Tier 1, support at either the 5′ or 3′ end; Tier 2, support at both the 5′ and 3′ ends (high-confidence full-length); Tier 3, support from all orthogonal datasets; and Tier 4, no end support from any dataset.

### Isoform Reproducibility Across Independent Long-Read Datasets

To assess isoform reproducibility, filtered transcripts were compared against four independent human brain long-read RNA sequencing datasets^21,22,36,37^, spanning bulk and single-cell data and both PacBio and Oxford Nanopore platforms (**Figure 4G**). Transcript concordance was assessed using Gffcompare^85^, and isoforms exhibiting exact intron-chain matches to transcripts identified in external datasets were considered reproducible.

### Evolutionary Conservation at Transcript Boundaries

Filtered SQANTI3 transcript annotations for each cell type in DLPFC and OFC were converted from GTF to BED format using UCSC utilities (gtfToGenePred and genePredToBed), and exonic intervals were extracted per isoform structural class (FSM, ISM, NIC, NNC, Other). Evolutionary conservation was quantified using PhastCons scores from the hg38 100-way vertebrate alignment^86^. Signal profiles were computed using deepTools computeMatrix (scale-regions mode, v3.5.6) spanning 3 kb upstream of the TSS, scaled gene bodies (5 kb), and 3 kb downstream, with 25 bp binning. Shuffled control regions were generated for each brain region using bedtools shuffle (v2.31.0), constrained to hg38 and excluding annotated transcripts (Ensembl release 102) and assembly gaps, to establish a genomic background for comparison.

### Gene and Isoform Quantification

Isoform-level expression was quantified by aligning short-read RNA-seq data to cell type-specific Iso-Seq-derived reference transcriptomes using Salmon (v2.0.1) ^87^ in quasi-mapping mode, generating transcript-level TPM and count estimates. Gene-level counts were derived by aggregating Salmon estimates using tximport (v1.34.0) ^88^ with transcript-to-gene mappings from Iso-Seq GTF annotations via txdbmaker. Quantification was performed independently for each cell type and region. GO enrichment was assessed using ToppGene^89^ across Molecular Function, Biological Process, and Cellular Component categories, with Benjamini-Hochberg correction applied; terms were considered significant at FDR < 5%.

### Identification of Shared Genes and Isoforms Across Cell Types and Regions

To identify genes and isoforms shared across cell types within each region, cell type-specific GTF files were merged using StringTie --merge^90^ to generate a unified regional reference, followed by comparison of each cell type-specific GTF against this reference using Gffcompare^85^. Transcripts assigned class code “=” were defined as exact isoform matches shared across cell types, and parent gene identifiers were used to determine shared genes. The same workflow was applied to identify genes and isoforms shared between DLPFC and OFC for GABA, GLU, and OLIG, the three lineages profiled in both regions.

### Protein-Coding Predictions

Coding potential was assessed using four independent frameworks: Coding-Potential Assessment Tool (CPAT, v1.2.2)^91^ using the human pre-trained logistic regression model (coding threshold ≥ 0.364), Coding Potential Calculator v2 (CPC2)^92^, Pfam domain scanning via pfam_scan against the Pfam database^93^, and GeneMarkS-T (GMST)^94^. CPAT and CPC2 predict coding potential from sequence features including open reading frame length, Fickett score, and hexamer usage bias; Pfam scanning identifies conserved protein domain matches within predicted peptide sequences; and GMST applies a self-training hidden Markov model to identify coding regions without requiring a reference annotation. Concordance across methods was used to evaluate robustness of coding predictions, while CPAT classifications were used as the primary coding potential metric for downstream analyses because it provides transcript-level probabilistic scoring optimized for human transcriptomes.

### RNA Secondary Structure Prediction

RNA secondary structure and minimum free energy (MFE) were predicted for transcripts using RNAfold from the ViennaRNA (v2.7.0) package^95,96^. This produced predicted secondary structures and corresponding MFE values for all input transcripts. RNAfold was executed with default parameters, which compute the MFE structure using thermodynamic folding rules from the ViennaRNA model^97^.

### Alternative Splicing Classification

Alternative splicing events were classified from Iso-Seq transcriptomes using SUPPA2 (v2.3)^98^ with the -f ioe flag to generate isoform-event association files. Seven event types were quantified: skipped exon (SE), mutually exclusive exons (MX), retained intron (RI), alternative first exon (AF), alternative last exon (AL), alternative 5′ splice site (A5), and alternative 3′ splice site (A3). NMD susceptibility was predicted based on established transcript features including premature termination codons, long 3′ UTRs, and upstream open reading frames.

### Region-Specific Transcriptome Construction for Differential Analyses

Prior to DGE, DTE, and DTU analyses, cell type-specific transcript annotations within each brain region were merged to generate unified region-specific reference transcriptomes. This approach enabled quantification against a shared regional reference containing all expressed isoforms while preserving cell type-specific differences in transcript abundance and usage for downstream comparisons. Short-read RNA-seq data for all samples within a given region were then aligned to the corresponding region-specific transcriptome using Salmon in quasi-mapping mode. For isoform-level quantification, transcript-level TPM and count estimates were generated for each sample.

For gene-level quantification, transcript-to-gene mappings were extracted from the merged GTF using txdbmaker and AnnotationDbi, and transcript-level estimates were aggregated using tximport with countsFromAbundance = “dtuScaledTPM”.

### Differential Gene and Transcript Expression

Expression matrices were filtered to retain features detected in at least one-third of samples per region and normalized using VOOM^99^ to model the mean-variance relationship. Principal component analysis confirmed the absence of sample outliers beyond two standard deviations from the mean. Differential expression was assessed using limma (v3.40.6)^100^ separately within each brain region, with cell type included as a fixed effect and donor modeled as a repeated measure where applicable. Comparisons were performed between (1) GABA versus GLU neurons and (2) neurons (GABA + GLU combined) versus OLIG. Significance thresholds were |log₂FC| ≥ 1.5 and FDR-adjusted p ≤ 0.05 (Benjamini-Hochberg). Gene set enrichment was assessed using CAMERA^101^ on Gene Ontology (GO) Biological Process gene sets using DGE and DTE summary statistics (DTE summarized at the gene level). Variance inflation was adjusted using default parameters (variance inflation factor = 0.01) and resulting p-values were corrected for multiple testing.

### Differential Transcript Usage

Differential transcript usage (DTU) was evaluated using DEXSeq (v1.52.0)^102^ implemented within IsoformSwitchAnalyzeR (v2.6.0)^103^. The same Salmon-derived isoform counts used for DTE analyses were supplied to the DTU workflow. Prior to testing, data were prefiltered to remove single-isoform genes and retain isoforms detected across all conditions; no minimum expression or isoform fraction thresholds were otherwise applied. DTU was assessed for GABA versus GLU neurons and neurons (GABA + GLU combined) versus OLIG separately in DLPFC and OFC. DEXSeq estimated differences in isoform fraction (ΔIF) between conditions while controlling for cell type and donor identity. Isoforms were considered significant at FDR-adjusted q < 0.05 and |ΔIF| > 10%.

GO enrichment analysis of significant DTU genes was performed using ToppGene^89^ for Biological Process (BP) categories. Isoform identifiers were mapped to parent genes using Ensembl gene annotations. P-values were corrected using the Benjamini-Hochberg procedure. To reduce redundancy and identify consensus functional groups, significant GO terms were summarized using REVIGO^104^ with normalized Resnik semantic similarity. Terms with dispensability < 0.15 were retained as representative non-redundant categories.

### Dominant Isoform Switching

Isoform usage was quantified as each isoform’s proportional contribution to total gene expression. The isoform with the highest proportional usage within a cell type was defined as the dominant isoform. Dominant isoform switching was defined as a change in dominant isoform identity between cell types accompanied by a difference in proportional usage exceeding 50%, assessed for significance by two-sample t-test (p < 0.05).

### Isoform Co-expression Network Analysis

Isoform co-expression networks were constructed using WGCNA (v1.68)^105^. Expression matrices were filtered to retain features present in at least one-quarter of samples and normalized using VOOM. Relevant covariates (brain region, cell type) were regressed from isoform expression matrices using linear mixed-effects models prior to network construction. Signed adjacency matrices were computed using soft-thresholding powers of β = 4 (DLPFC) and β = 5 (OFC) and converted to signed topological overlap matrices. Isoform clustering used hclust with average linkage; modules were defined by Dynamic Hybrid Tree Cut with a minimum cluster size of 50 isoforms. Module annotation used ToppGene^89^ for GO enrichment and REVIGO^104^ for term summarization, as described above.

Co-expression modules were summarized at the gene level and tested for enrichment of neurodevelopmental disorder-associated genes using GeneOverlap (v3.19)^106^, with gene lists derived from published transcriptomic and genetic studies^30–32^. Enrichment p-values were Benjamini-Hochberg corrected. Novel isoforms belonging to ASD-associated genes were intersected with de novo variants by overlapping variant genomic coordinates with isoform-specific exon structures using corresponding BAM alignments; variants were considered intersecting if their mapped position overlapped any exon or splice junction supported by isoform-aligned reads.

### Disease-Variant Intersection with Long-read Isoform Annotations

To determine whether previously unannotated transcript structures expose exonic sequence overlapping pathogenic genetic variation, full-length long-read isoform annotations from DLPFC and OFC were intersected with high-confidence disease-causing mutations (DM class) from the Human Gene Mutation Database (HGMD)^41^, represented as strand-aware hg38/GRCh38 BED6 intervals. Isoforms were grouped by structural category (FSM, NIC, NNC), and exon-level coordinates were intersected with HGMD DM variants using strand-aware bedtools overlap to quantify unique disease variants overlapping each transcript class.

NNC isoforms were further classified by splice-site architecture. Exon-level BED6 intervals were compared with GENCODE v38 annotations on the matched strand using strand-aware donor and acceptor site catalogs. NNC exons were assigned hierarchically as follows: exact GENCODE matches (known_exact); novel donor events, defined as use of a previously unannotated 5′ splice donor with retention of a known acceptor; novel acceptor events, defined as use of a previously unannotated 3′ splice acceptor with retention of a known donor; internal exon extensions (boundary shifts), defined as exons that overlapped annotated exonic sequence but exhibited noncanonical boundaries and did not meet criteria for isolated novel donor or novel acceptor events; and novel exons, defined as exons with no overlap with annotated exons and further subdivided into intronic or intergenic subclasses.

Disease-variant enrichment within each NNC feature class relative to GENCODE v38 annotated exons was assessed by Fisher’s exact test using hit/non-hit 2×2 contingency tables, with Benjamini-Hochberg correction. Splice-proximal variant clustering was evaluated by expanding novel splice boundaries by ±5 bp and intersecting with HGMD variants; distance distributions across ±50 bp were computed using bedtools closest with strand-aware signed distances.

For gene-level analyses, non-intergenic novel splice features were assigned to genes by stranded intersection with GENCODE v38 gene intervals. Per-gene novel splice-feature counts and HGMD variant counts were calculated with and without normalization for gene length. Representative loci with high splice complexity and disease-variant burden, including *PLP1* and *TARDBP*, were visualized with long-read isoform structures, NNC-derived splice features, and HGMD variant positions to assess overlap with exonic sequence exposed by non-reference splice-boundary usage. All interval operations used strand-aware bedtools; statistical analyses and visualization were performed in R.

### Quantifying Cell-type Specificity with Tau

To quantify the cell-type specificity of isoform usage, we adapted the Tau index, originally developed to measure tissue-specific gene expression^107,108^. The Tau index for each transcript was calculated as follows:

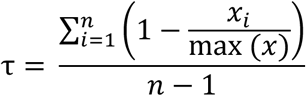

where *x*_i_ denotes isoform proportion in cell type *i, n* the total of cell types, and max (*x*) the maximum isoform proportion across cell types. Tau ranges from 0 to 1, with values approaching 1 indicating high cell-type specificity and values near 0 indicating uniform usage across cell lineages.

## SUPPLEMENTAL INFORMATION

**Document S1**. Supplemental Figures S1-S24.

**Table S1**. PacBio HiFi Iso-Seq CCS and FLNC Read Metrics by Donor, Sample, Cell Type, and Brain Region

**Table S2**. PacBio Iso-Seq Quality Control Metrics and Sample Metadata

**Table S3**. Illumina FANS RNA-Seq Quality Control Metrics and Sample Metadata

**Table S4**. Cell Type-Specific Genes in the DLPFC and OFC

**Table S5**. Genes with High Isoform Complexity (≥10 Isoforms per Gene) in the DLPFC and OFC

**Table S6**. Gene Ontology Biological Process Enrichment of Genes with High Isoform Complexity

**Table S7**. Highly Expressed Novel Isoforms in the DLPFC and OFC

**Table S8**. Iso-Seq Isoforms Mapped to Uncharacterized Gene Loci Across Cell Types and Brain Regions

**Table S9**. Cell Type- and Region-Resolved Protein-Coding Intron-Retaining Transcripts Predicted to Undergo Nonsense-Mediated Decay

**Table S10**. Genes Ranked by the Number of Intron Retention-Containing Transcript Events

**Table S11**. Differential Gene Expression Summary Statistics

**Table S12**. Differential Transcript Expression Summary Statistics

**Table S13**. Differential Transcript Usage Summary Statistics

**Table S14**. Gene Ontology Enrichment Summary for Differential Gene Expression (CAMERA)

**Table S15**. Gene Ontology Enrichment Summary for Differential Transcript Expression (CAMERA)

**Table S16**. Dominant Isoform Switching Events in the DLPFC and OFC

**Table S17**. WGCNA Module Summary Statistics

**Table S18**. Correlations Between WGCNA Module Eigengenes and Major Brain Cell Types

**Table S19**. Gene Ontology Enrichment of WGCNA Modules

**Table S20**. Novel NIC and NNC Isoforms Overlapping ASD Risk Genes

**Table S21**. ASD De Novo Variants Overlapping Novel NIC and NNC Isoforms

**Table S22**. Gene-level view of isoform specificity, splice burden and variant burden

