## Supplemental Figures 1-24 for "Long-read transcriptomics of purified human cortical cell types exposes glial isoform complexity and disease-relevant transcript architecture"

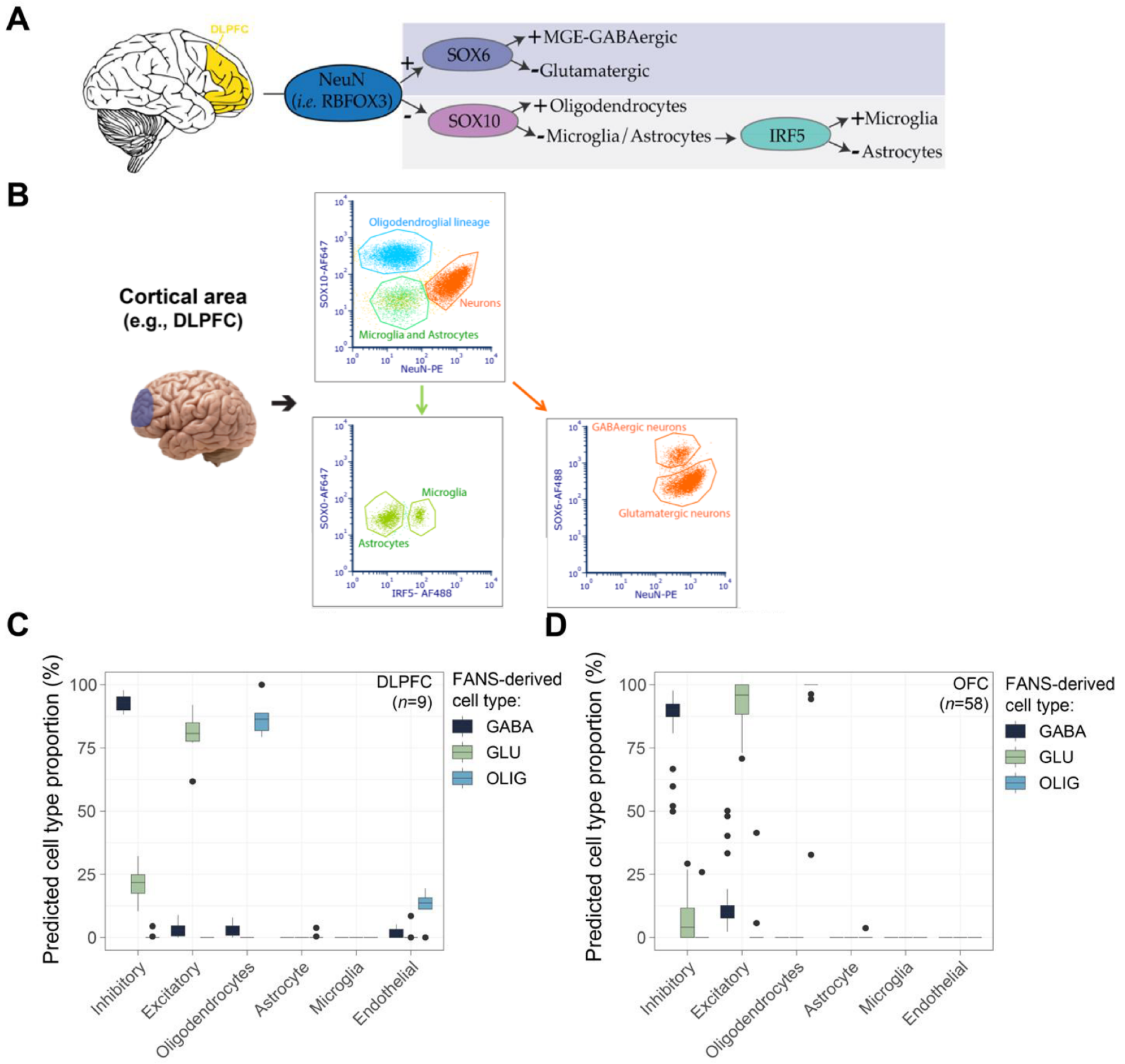


**Figure S1. FANS workflow, gating strategy, and short-read-based cell type validation. (A)** Overview of the fluorescence-activated nuclei sorting (FANS) strategy used to isolate five major cortical lineages from adult DLPFC and OFC tissue. NeuN (RBFOX3) was used to partition neuronal ($NeuN^{+}$) and non-neuronal ($NeuN^{-}$) nuclei, followed by SOX6 to enrich MGE-derived GABAergic neurons, SOX10 to isolate oligodendrocytes, and IRF5 to distinguish microglia from astrocytes within the non-neuronal pool. **(B)** Representative FANS gating plots illustrating the separation of neuronal and glial nuclei based on NeuN and SOX10, and the subsequent discrimination of OLIG, MG, and AST subpopulations using SOX10 and IRF5. Cell type deconvolution of matched short-read RNA-seq data for (**C**) DLPFC and (**D**) OFC nuclei demonstrates that each FANS-sorted population is strongly enriched for its expected transcriptional signature across six major cortical cell classes (inhibitory neurons, excitatory neurons, oligodendrocytes, astrocytes, microglia, endothelial cells), supporting the purity and specificity of the sorting strategy.

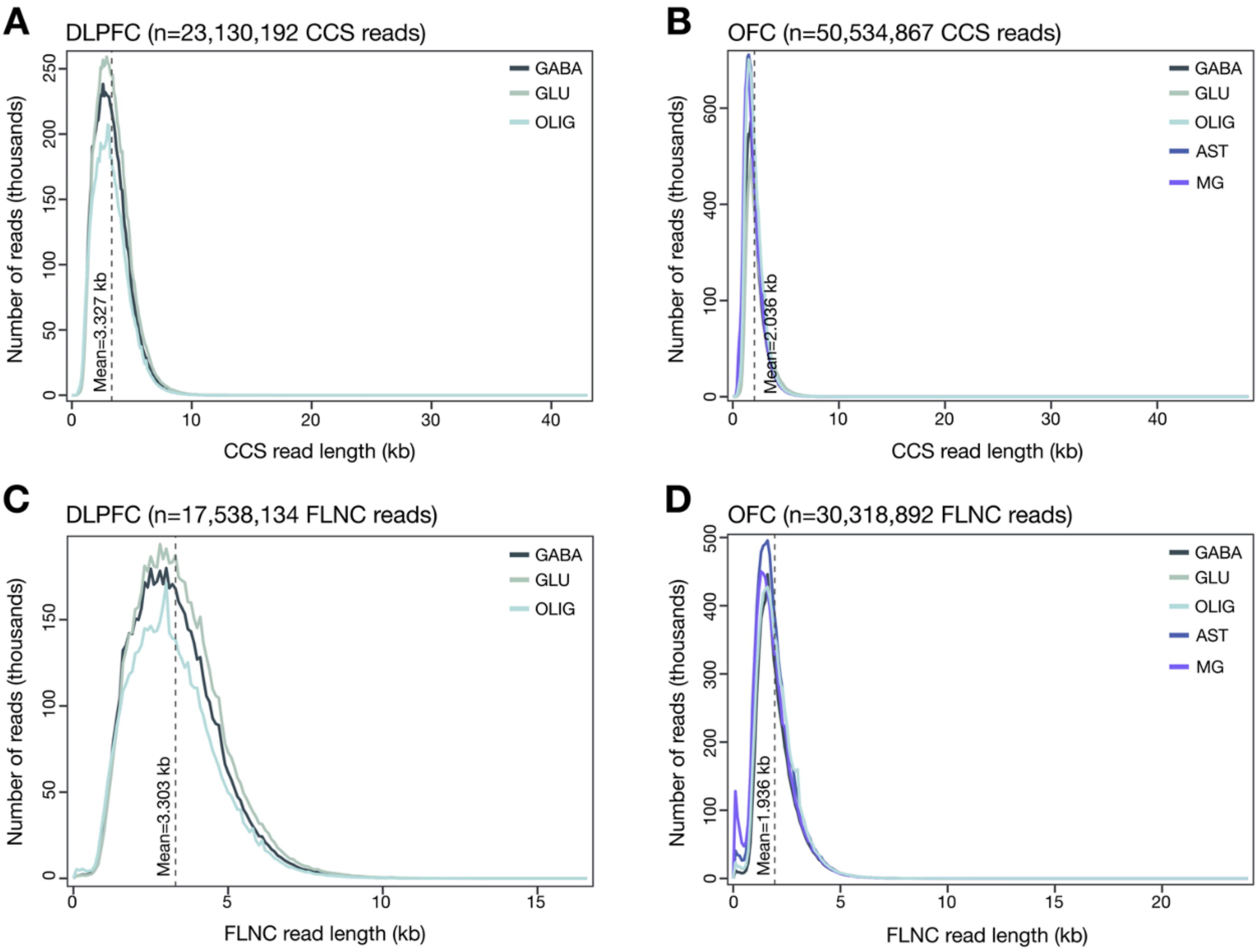


**Figure S2. Distribution of circular consensus (CCS) and full-length non-chimeric (FLNC) read lengths across cortical cell types.** Distribution of circular consensus (CCS) and full-length non-chimeric (FLNC) read lengths across major cortical cell types. (**A**, **B**) Aggregate mean CCS read length (kb) across cell types in the dorsolateral prefrontal cortex (DLPFC) and orbitofrontal cortex (OFC), respectively. (**C**, **D**) Aggregate mean FLNC read length across cell types in the DLPFC and OFC. For each region, aggregate means were calculated as the average of cell types in the DLPFC and OFC. GABA, MGE-derived GABAergic interneurons; GLU, glutamatergic neurons; OLIG, oligodendrocytes; AST, astrocytes; MG, microglia.

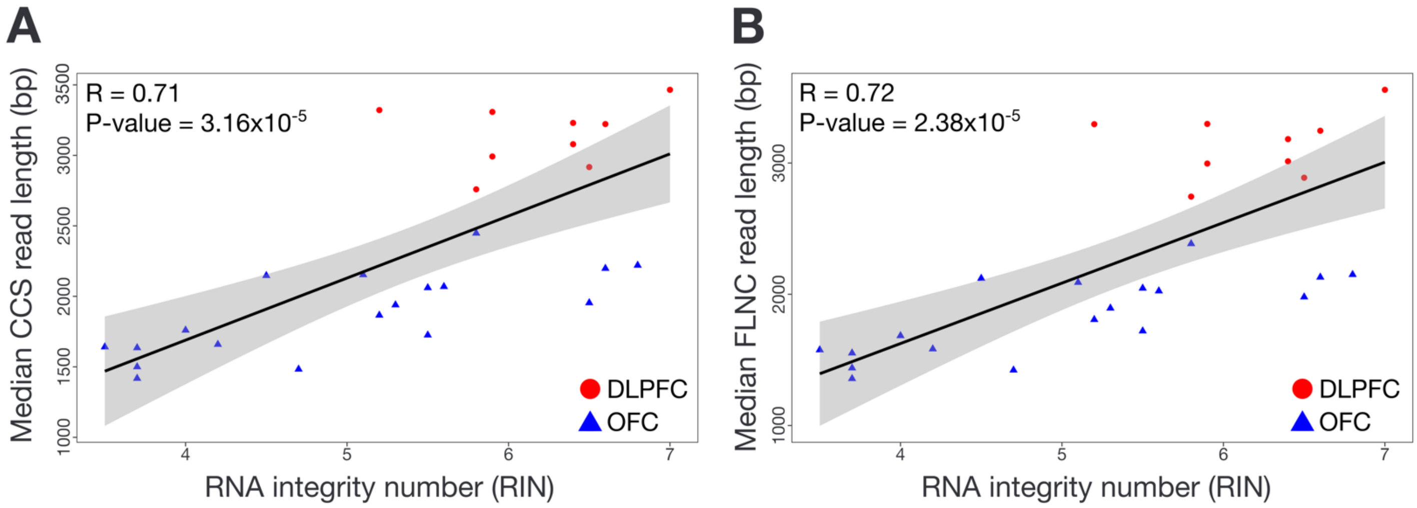


**Figure S3. RNA integrity (RIN) correlates with median long-read length in Iso-Seq libraries.** (**A**) Relationship between RNA integrity number (RIN) and median circular consensus sequence (CCS) read length across all DLPFC (red) and OFC (blue) Iso-Seq libraries. Each point represents a cell type-specific library from an individual donor. Higher RIN values were associated with longer CCS reads ($R = 0.71$, $P = 3.16 \times 10^{-5}$), consistent with expected effects of RNA quality on full-length transcript recovery in long-read sequencing. (**B**) Relationship between RIN and median full-length non-chimeric (FLNC) read length. RIN showed a similar positive correlation with FLNC read length ($R = 0.72$, $P = 2.38 \times 10^{-5}$). Despite this technical association, all samples yielded sufficient read lengths and depth for high-confidence splice-junction support, SQANTI3 classification, and downstream isoform reconstruction.

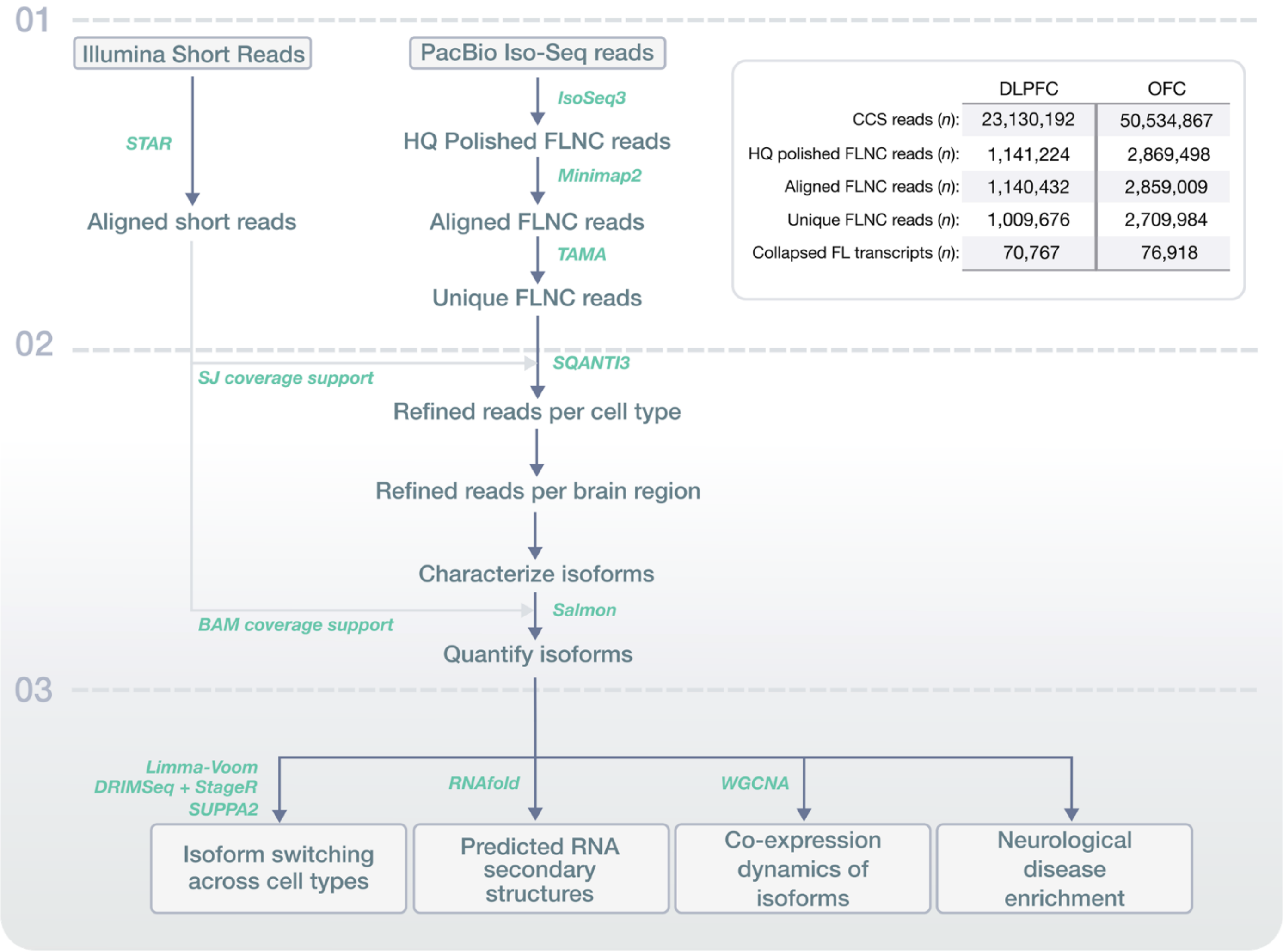


| | DLPFC | OFC |
|---|---|---|
| CCS reads (*n*): | 23,130,192 | 50,534,867 |
| HQ polished FLNC reads (*n*): | 1,141,224 | 2,869,498 |
| Aligned FLNC reads (*n*): | 1,140,432 | 2,859,009 |
| Unique FLNC reads (*n*): | 1,009,676 | 2,709,984 |
| Collapsed FL transcripts (*n*): | 70,767 | 76,918 |

**Figure S4. Data processing and analysis pipeline for full-length transcript annotation in DLPFC and OFC cell types.** Data analysis largely consisted of three main phases. (1) Data pre-processing: CCS reads from long-read sequencing were processed using IsoSeq3 and TAMA to generate high-quality full-length transcripts. (2) Quantifying high-confidence isoform annotations: SQANTI3 annotated individual isoforms, comparing them to short-read RNA-Seq datasets and reference annotations. (3) Downstream analyses: Downstream analyses characterized isoform expression patterns, including cell type-specific expression, isoform switching, and co-expression modules, revealing the transcriptomic landscape of DLPFC and OFC subregions.

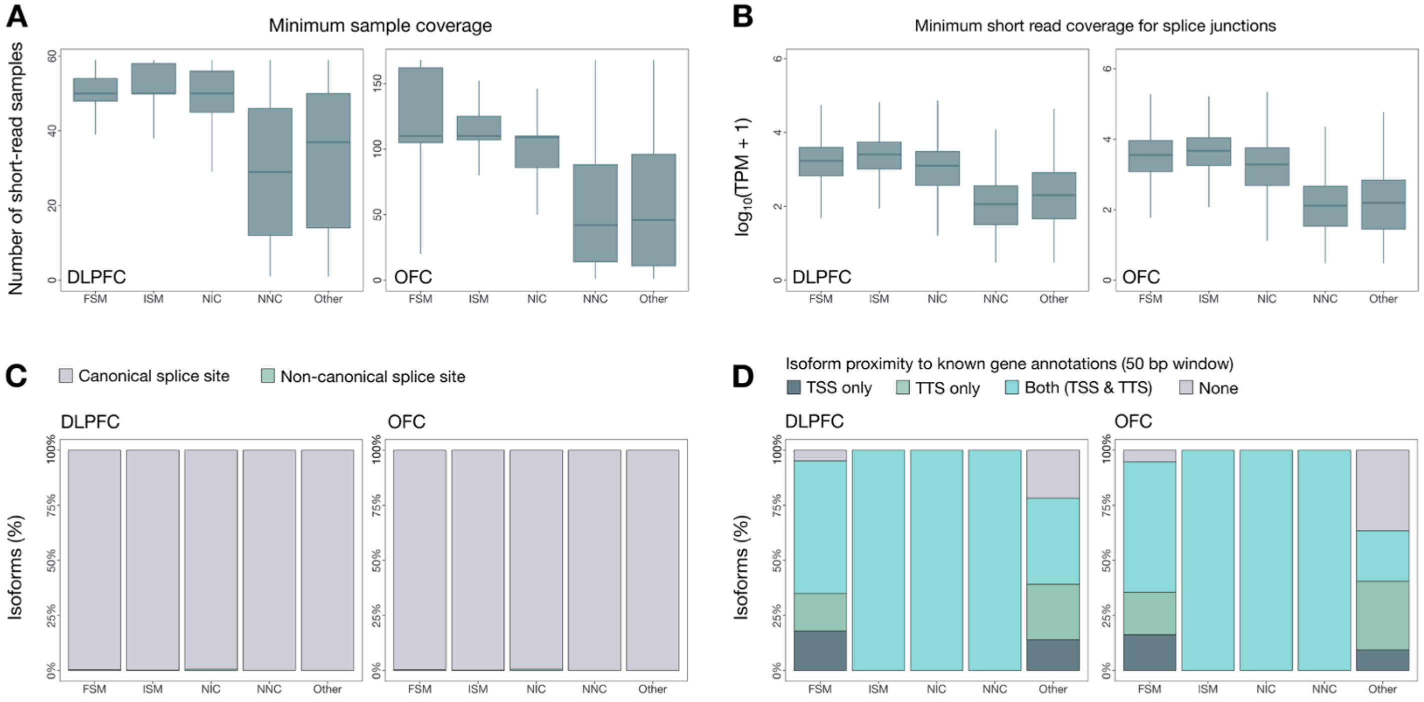


**Figure S5. SQANTI3 quality control metrics across cortical cell types.** (**A**) Minimum sample coverage across all isoforms, grouped by structural category, in the DLPFC and OFC. (**B**) Minimum short-read coverage for splice junctions across isoforms classified by structural category in the DLPFC and OFC. (**C**) Percentage of isoforms with canonical splice sites in each region. (**D**) Percentage of isoforms in each structural category with transcription start site (TSS), transcription termination site (TTS), or both TSS/TTS support relative to annotated gene ends. Isoform proximity at 5' and 3' ends was defined within a 50 bp window. FSM, full-splice match; ISM, incomplete-splice match; NIC, novel-in-catalog; NNC, novel-not-in-catalog.

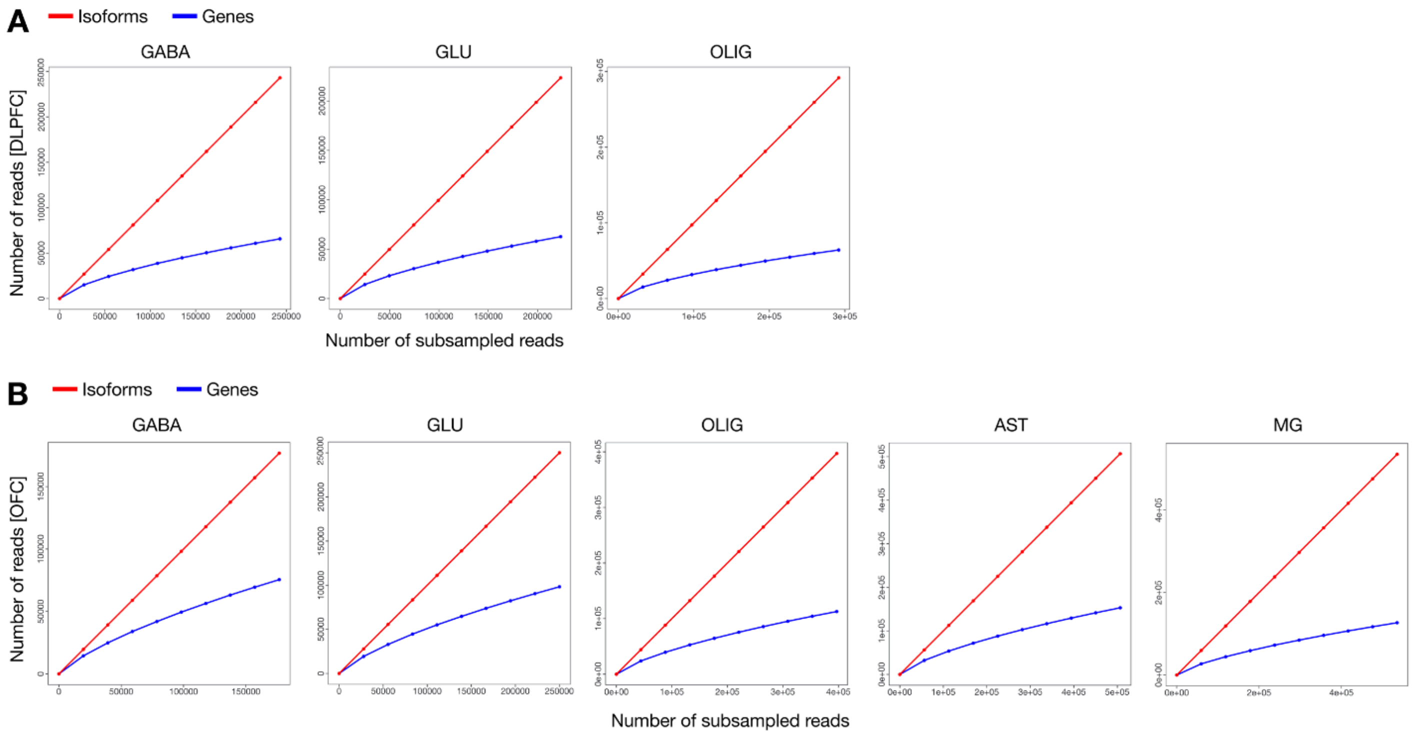


**Figure S6. Iso-Seq sequencing saturation curves across cortical cell types**. **(A-B)** Rarefaction curves showing the number of detected genes (blue) and isoforms (red) as a function of subsampled CCS read depth across GABA, GLU, and OLIG nuclei from the DLPFC (A) and across GABA, GLU, OLIG, AST, and MG nuclei from the OFC (B). Gene-level detection approaches saturation in all cell types, indicating sufficient depth to recover the majority of expressed genes. In contrast, isoform detection increases nearly linearly with sequencing depth and does not plateau, reflecting the substantially higher structural complexity of full-length transcripts.

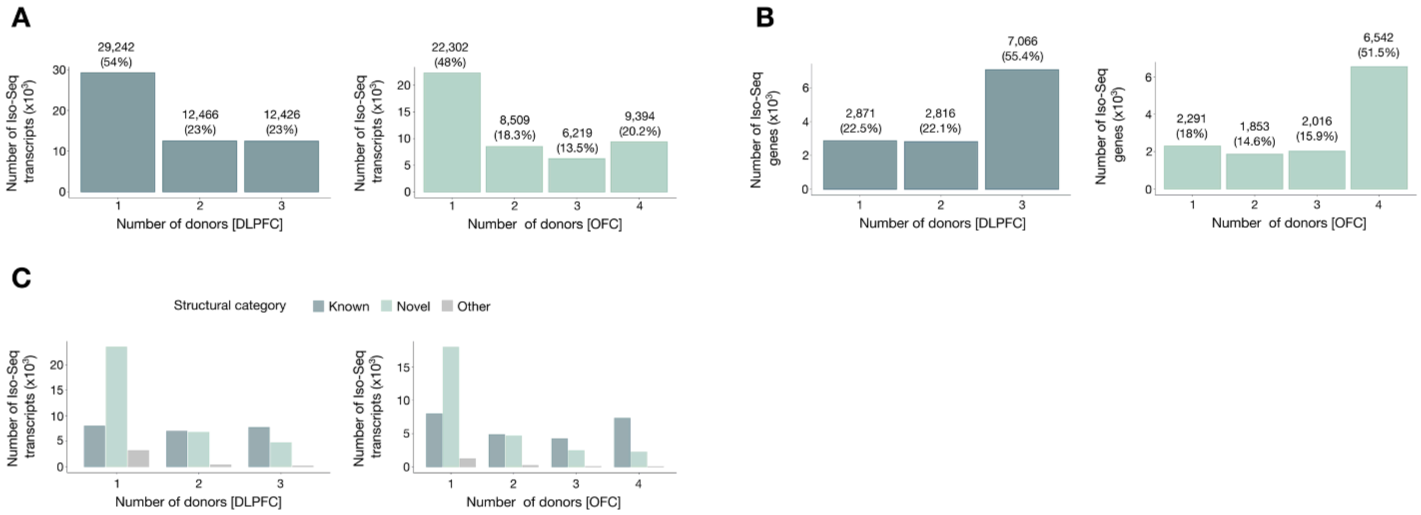


**Figure S7. Reproducibility of Iso-Seq gene and isoform detection across donors**. (A) Bars reflect the number of Iso-Seq transcripts detected in exactly 1, 2, 3, or all donors within each region. Most transcripts appear in only one donor, consistent with the expected rarity and shallow sampling of many low-abundance isoforms in long-read data. (B) In contrast to transcripts, most genes are detected in every donor (N = 3 for DLPFC; N = 4 for OFC), demonstrating strong cross-individual consistency at the gene level. (C) Counts of transcripts detected across donors, stratified by structural category (known FSM/ISM; novel NIC/NNC; other). Novel isoforms show the greatest donor-to-donor variability, as expected for low-abundance and cell-state-restricted transcript classes.

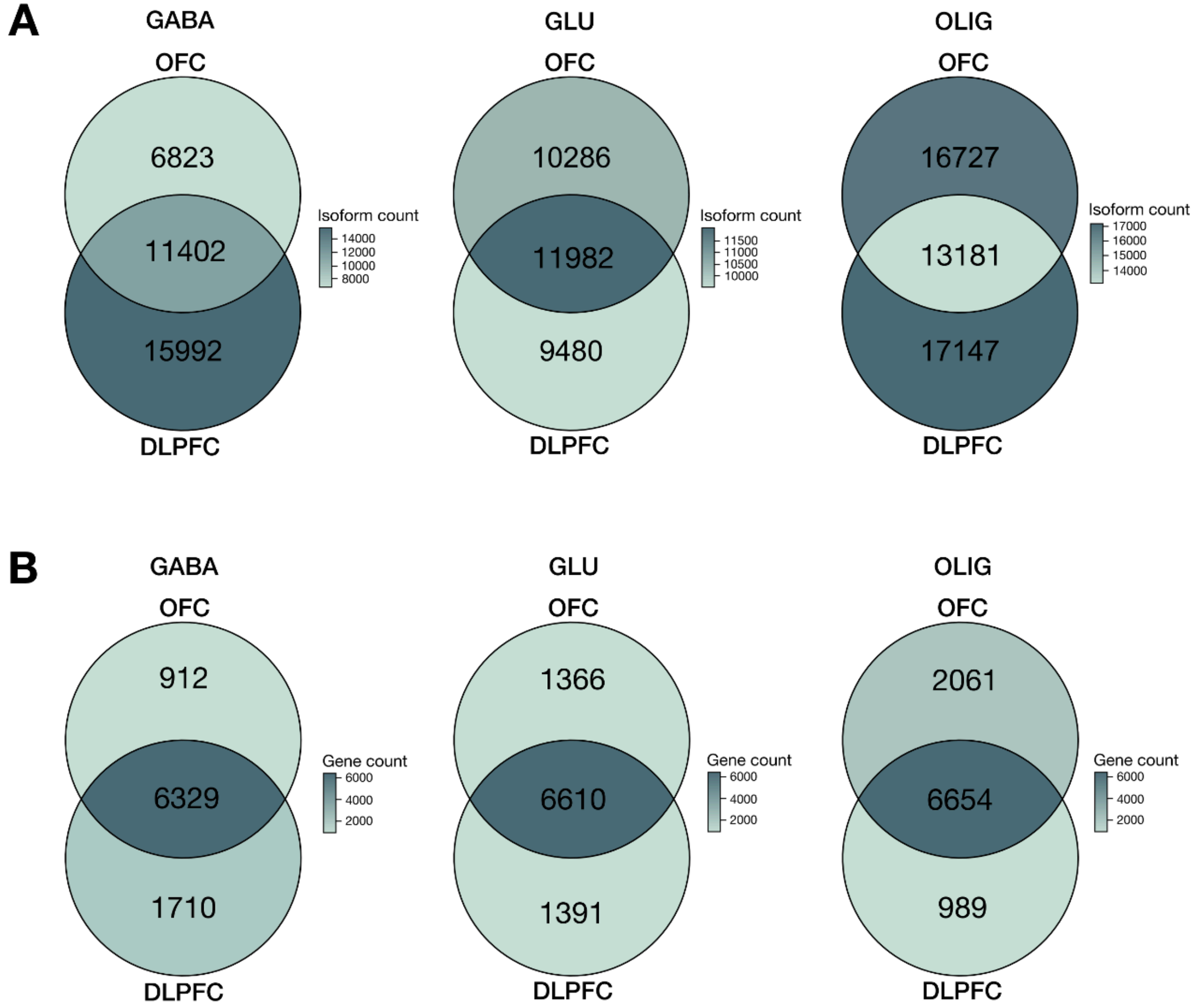


**Figure S8. Reproducibility of isoform and gene detection across cortical regions.** (A) Overlap of detected isoforms between the DLPFC and OFC within matched cell types (GABA, GLU, OLIG). Values indicate the number of isoforms detected uniquely in each region and those shared across both regions. (B) Overlap of detected genes between the same cell types in the DLPFC and OFC. Gene-level overlap was consistently higher than isoform-level overlap, reflecting the expected cell-type stability of gene expression and the added sensitivity of long-read sequencing for regionally specific isoform discovery.

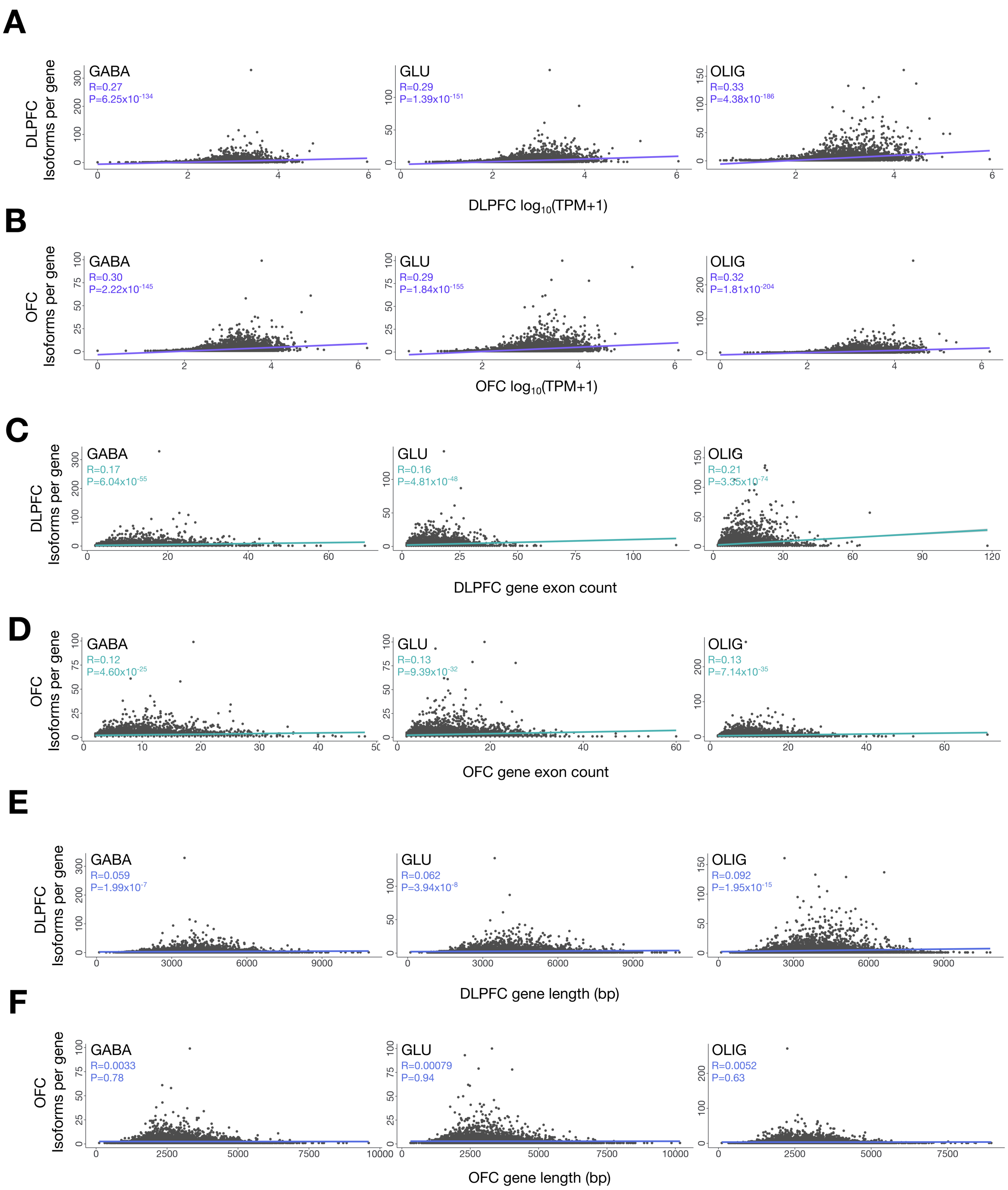


**Figure S9. Relationship between gene features and isoform diversity across cortical cell types**. Genes with higher expression or more exons exhibit more detected isoforms, whereas gene length shows no consistent association. (**A-B**) Pearson correlations between gene expression and the number of isoforms per gene in the DLPFC and OFC. (**C-D**) Correlations between exon count and isoform number in the DLPFC and OFC. (**E-F**) Correlations between mean gene length (averaged across isoforms) and isoform number in the DLPFC and OFC.

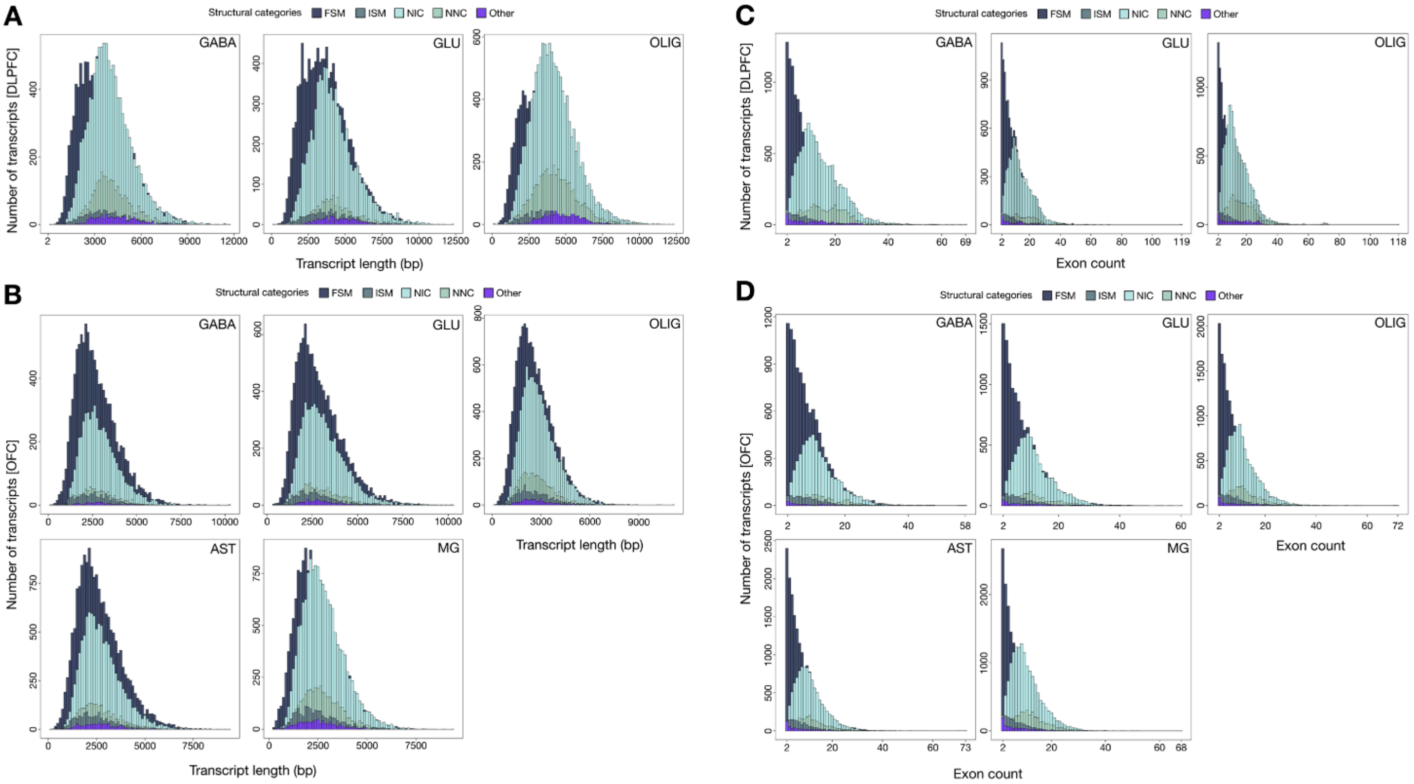


**Figure S10. Transcript length and exon-count distributions across cortical cell types.** (**A-B**) Density distributions of full-length transcript lengths for GABA, GLU, OLIG, AST, and MG isoforms in the DLPFC and OFC. (**C-D**) Corresponding exon-count distributions for the same cell types. Distributions highlight broad similarity in overall isoform length ranges across lineages but reveal marked cell-type-specific differences in the long-tail regions, particularly the enrichment of long, exon-rich transcripts in OLIG and MG populations. Structural categories (FSM, ISM, NIC, NNC, and Other) are shown to illustrate contributions of known and novel isoforms across the full complexity spectrum.

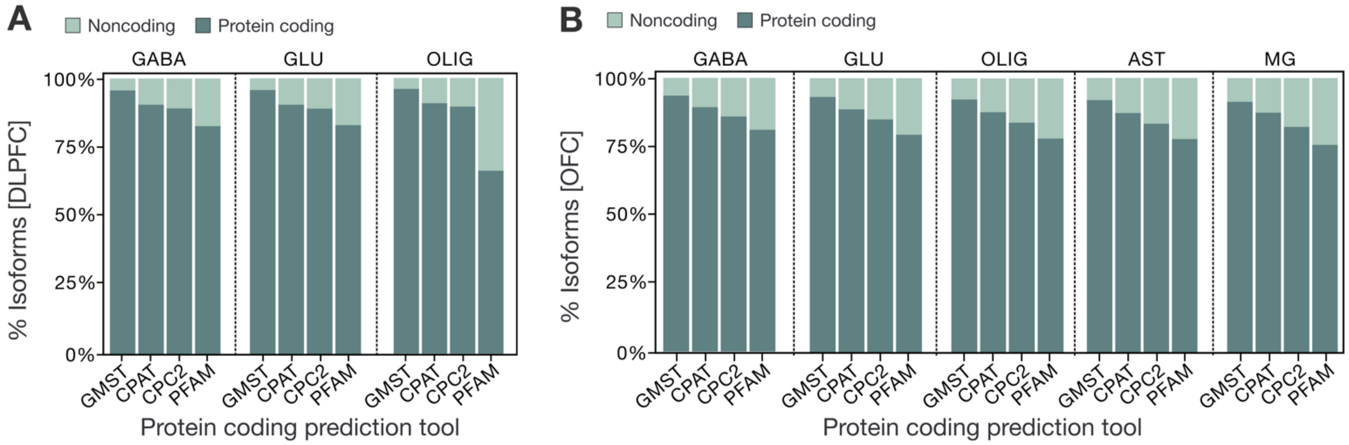


**Figure S11. Evaluation of protein-coding potential for known and novel isoforms across cortical cell types.** (A-B) Percentage of all isoforms predicted to be protein-coding or noncoding by four independent computational frameworks (GMST, CPAT, CPC2, and PFAM) in the DLPFC (A) and OFC (B). Each panel displays cell-type-resolved predictions (GABA, GLU, OLIG; and in OFC, AST and MG), illustrating the high concordance among tools and the predominance of protein-coding predictions across all cortical lineages.

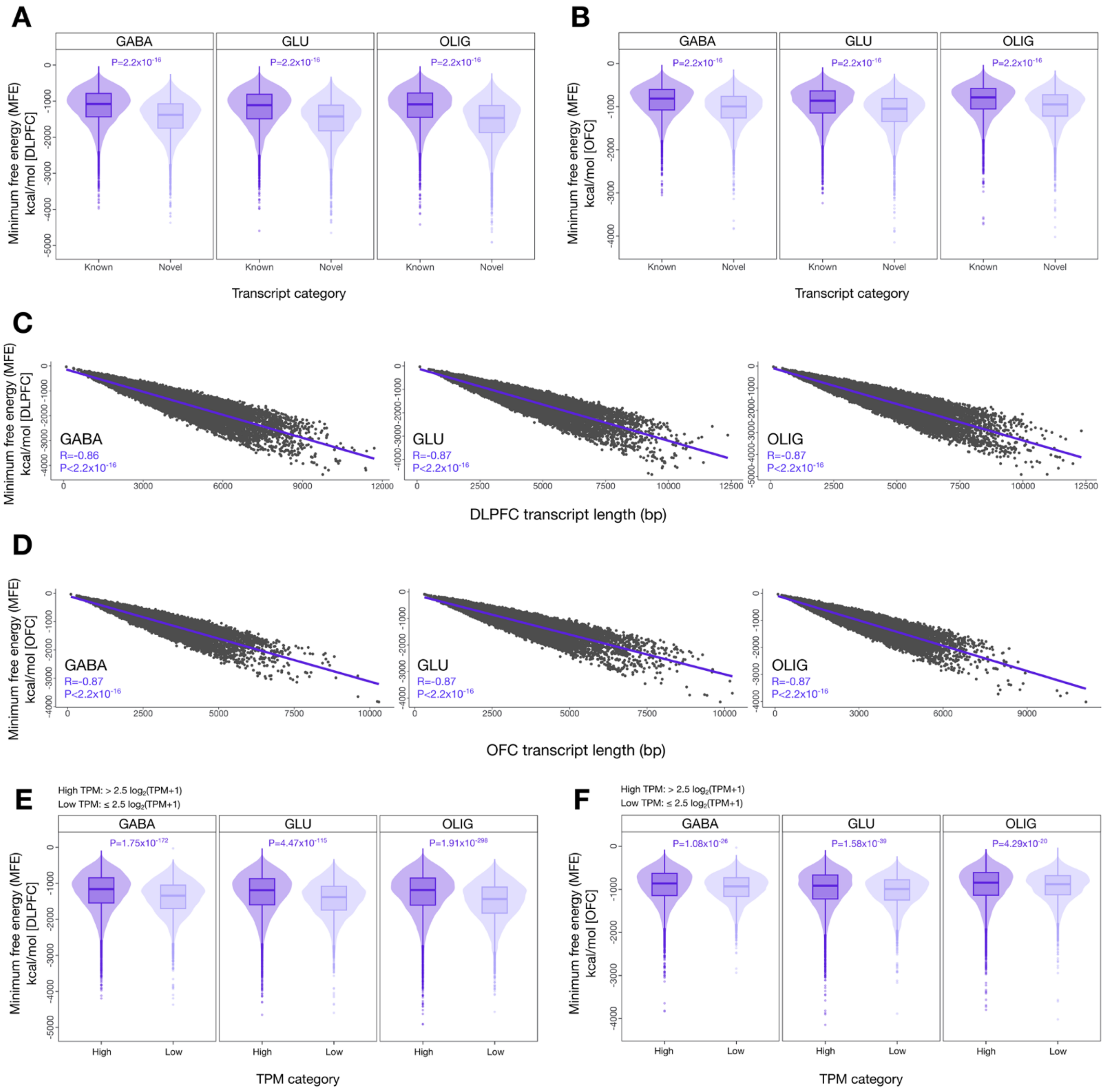


**Figure S12. RNA secondary structure and minimum free energy (MFE) analysis across transcripts**. (**A**, **B**) Comparison of MFE (kcal/mol) between known (FSM and ISM) and novel (NIC and NNC) transcripts across cortical cell types in the DLPFC and OFC. Statistical significance was assessed using a two-sided Mann-Whitney U test (all comparisons significant). (**C**, **D**) Pearson's correlation between MFE and transcript length (bp) across all cell types in the DLPFC and OFC. (**E**, **F**) Comparison of MFE between transcripts with high (>2.5 log2[TPM+1]) and low (≤2.5 log2[TPM+1]) expression levels across cell types in the DLPFC and OFC. Statistical significance was evaluated using a two-sided Mann-Whitney U test.

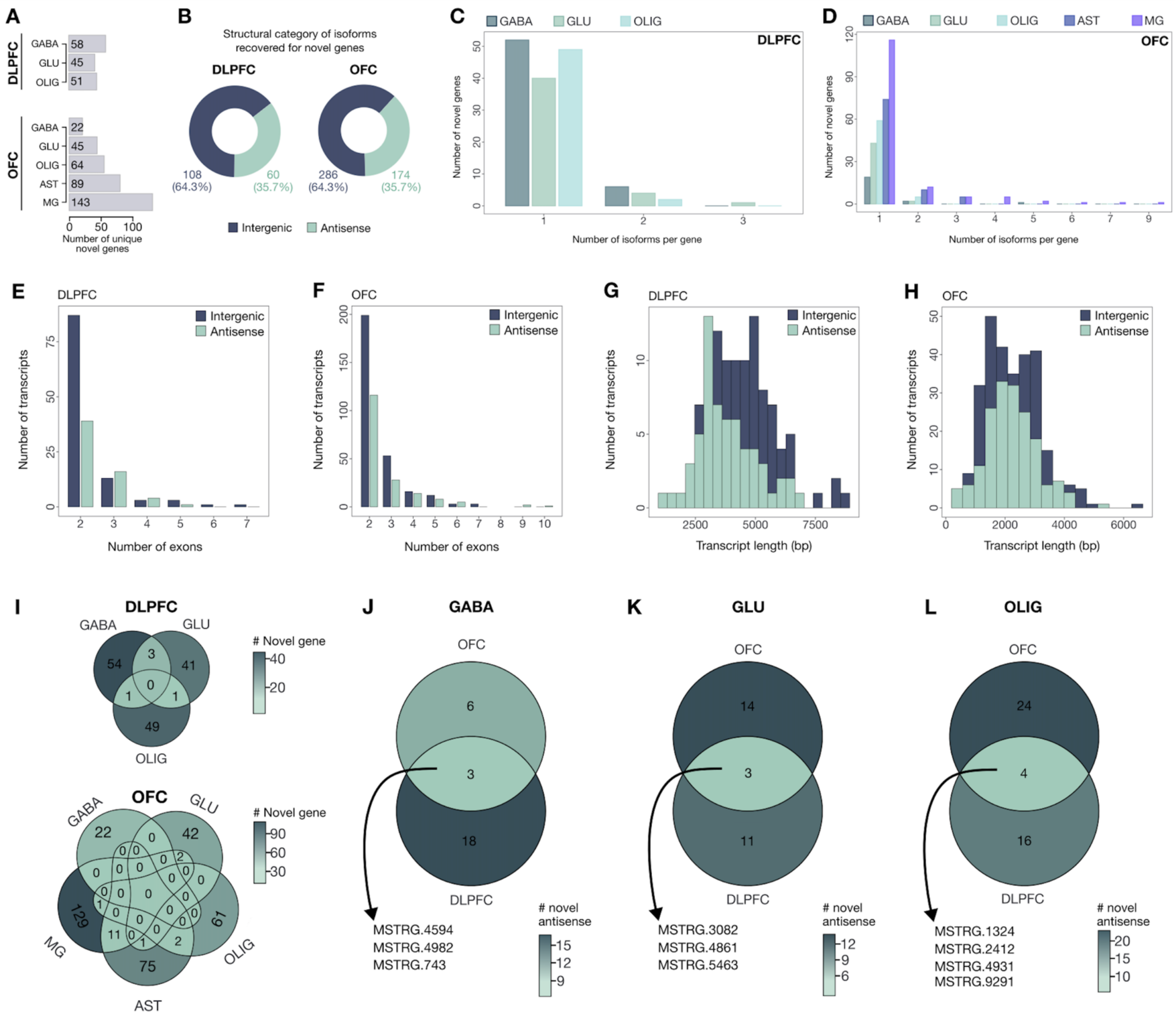

**Figure S13. Iso-Seq isoforms mapping to uncharacterized gene loci**. (**A**) Number of novel genes identified across cortical cell types in the DLPFC and OFC. (**B**) Number and percentage of isoforms corresponding to novel genes, stratified by structural category. (**C**, **D**) Number of isoforms per novel gene across cell types in the DLPFC and OFC, respectively. (**E**, **F**) Distribution of exon counts in isoforms mapping to novel genes across cell types in the DLPFC and OFC. (**G**, **H**) Transcript length (bp) of isoforms mapping to novel genes across cell types in the DLPFC and OFC. (**I**) Overlap of novel genes across cell types between the DLPFC and OFC. (**J-L**) Comparison of novel genes with antisense transcripts between matched cell types (GABAergic interneurons, glutamatergic neurons, and oligodendrocytes) in the DLPFC and OFC. Novel genes with antisense transcripts are listed in **Table S8**.

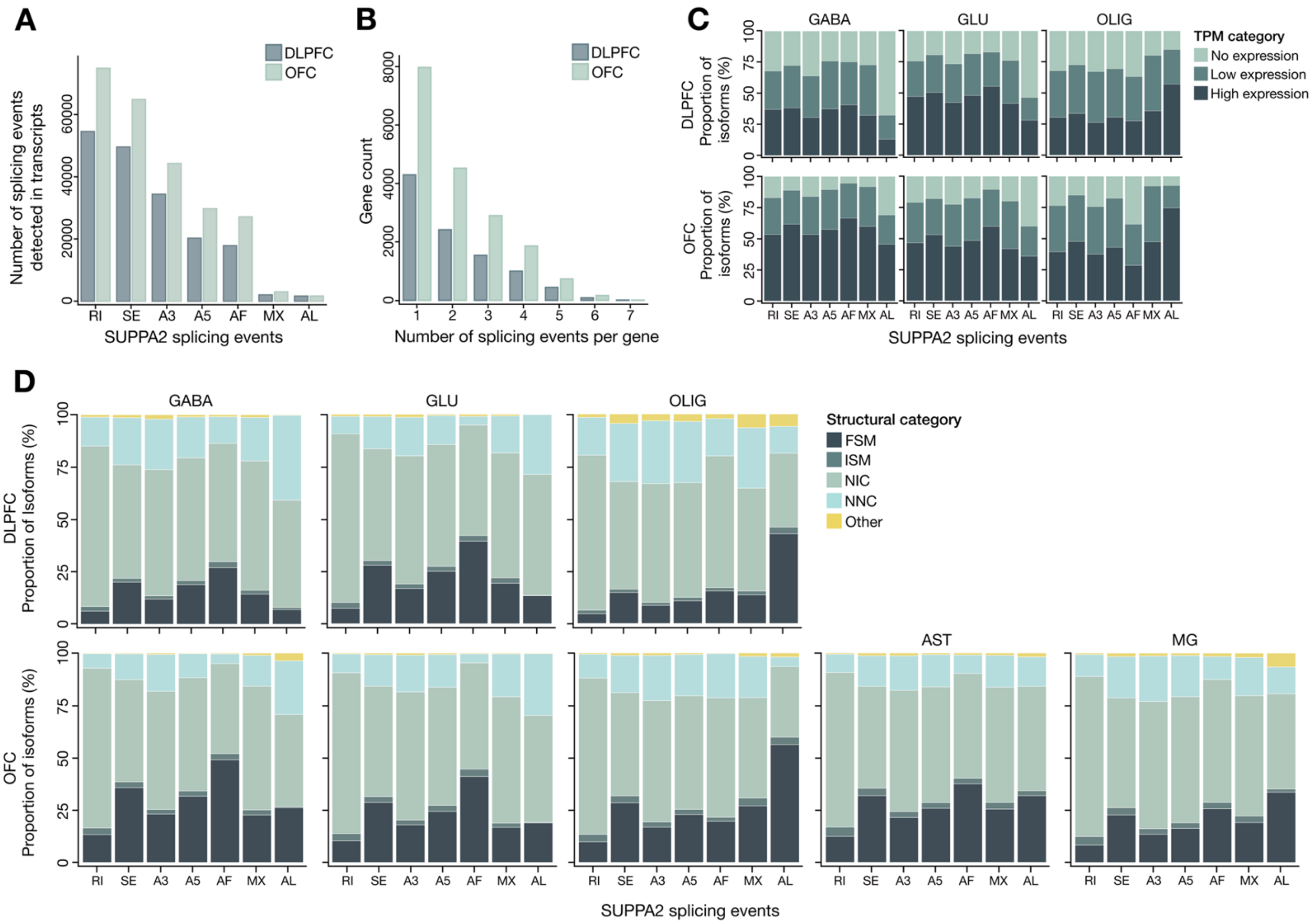


**Figure S14. Alternative splicing events across cortical cell types**. **(A)** Total number of alternative splicing events detected across transcripts in the DLPFC and OFC. **(B)** Number of splicing events per gene in the DLPFC and OFC. **(C)** Proportion of isoforms with high, low, or undetectable expression (based on $\log_2$[TPM]+1] thresholds) across SUPPA2 event types in DLPFC cell types (top row) and OFC cell types (bottom row). **(D)** Distribution of isoforms across structural categories for each SUPPA2 event type in DLPFC cell types (top row) and OFC cell types (bottom row).

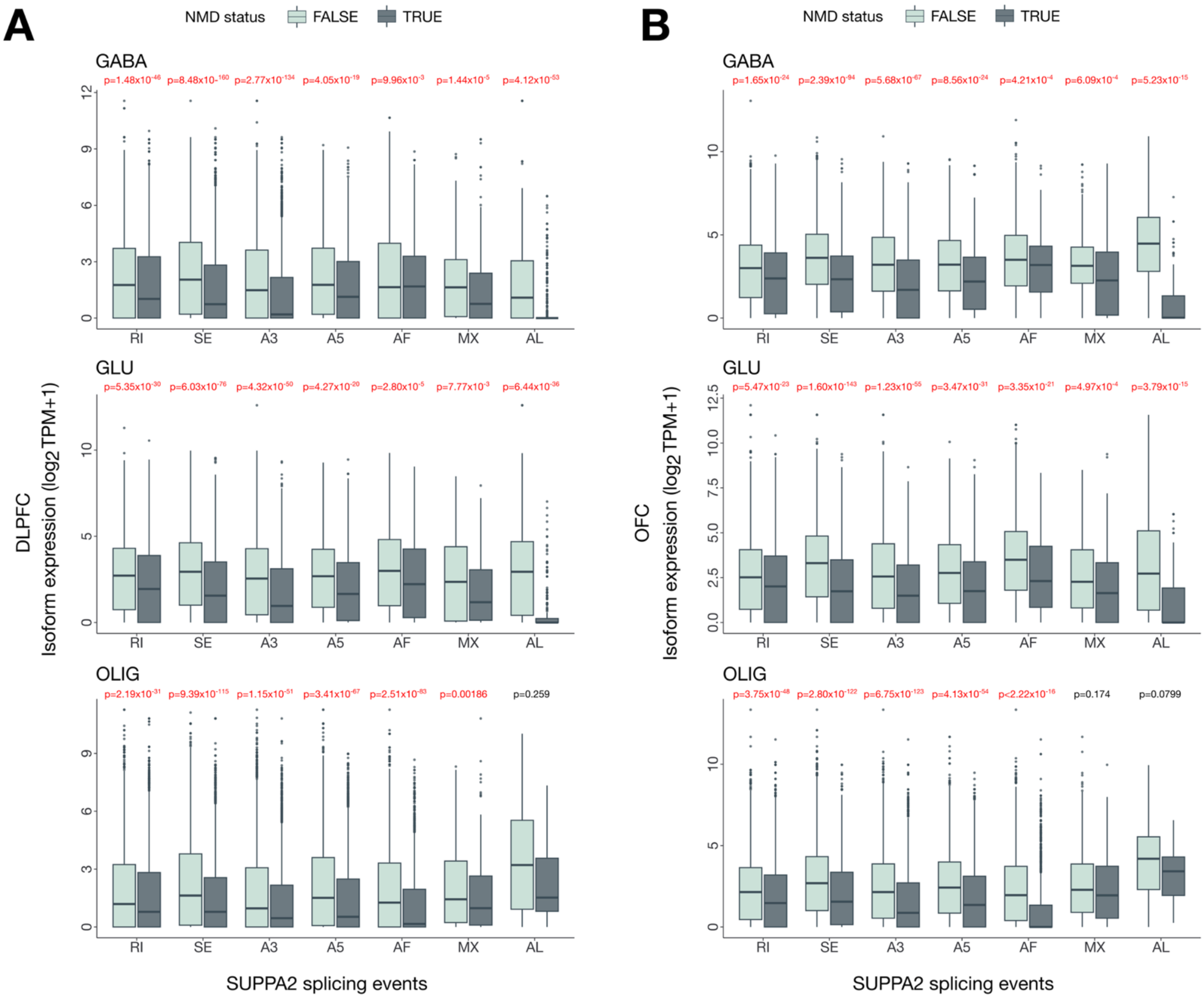


**Figure S15. Expression of protein-coding transcripts subject to nonsense-mediated decay (NMD) across cortical cell types**. (A-B) Distribution of expression levels ($\log_2$[TPM+1]) for protein-coding transcripts predicted to undergo nonsense-mediated decay (NMD; "TRUE") versus those not targeted by NMD ("FALSE") across SUPPA2 splicing event classes, retained intron (RI), skipped exon (SE), alternative 3′ splice site (A3), alternative 5′ splice site (A5), mutually exclusive exons (MX), alternative first exon (AF), and alternative last exon (AL), in the DLPFC (A) and OFC (B). Analyses are shown separately for each major cortical cell type (GABAergic neurons, glutamatergic neurons, and oligodendrocytes). For nearly every event type and lineage, NMD-predicted isoforms displayed significantly lower expression than their non-NMD counterparts (two-sided Mann-Whitney U test), consistent with active NMD-mediated repression of a substantial fraction of alternatively spliced transcripts.

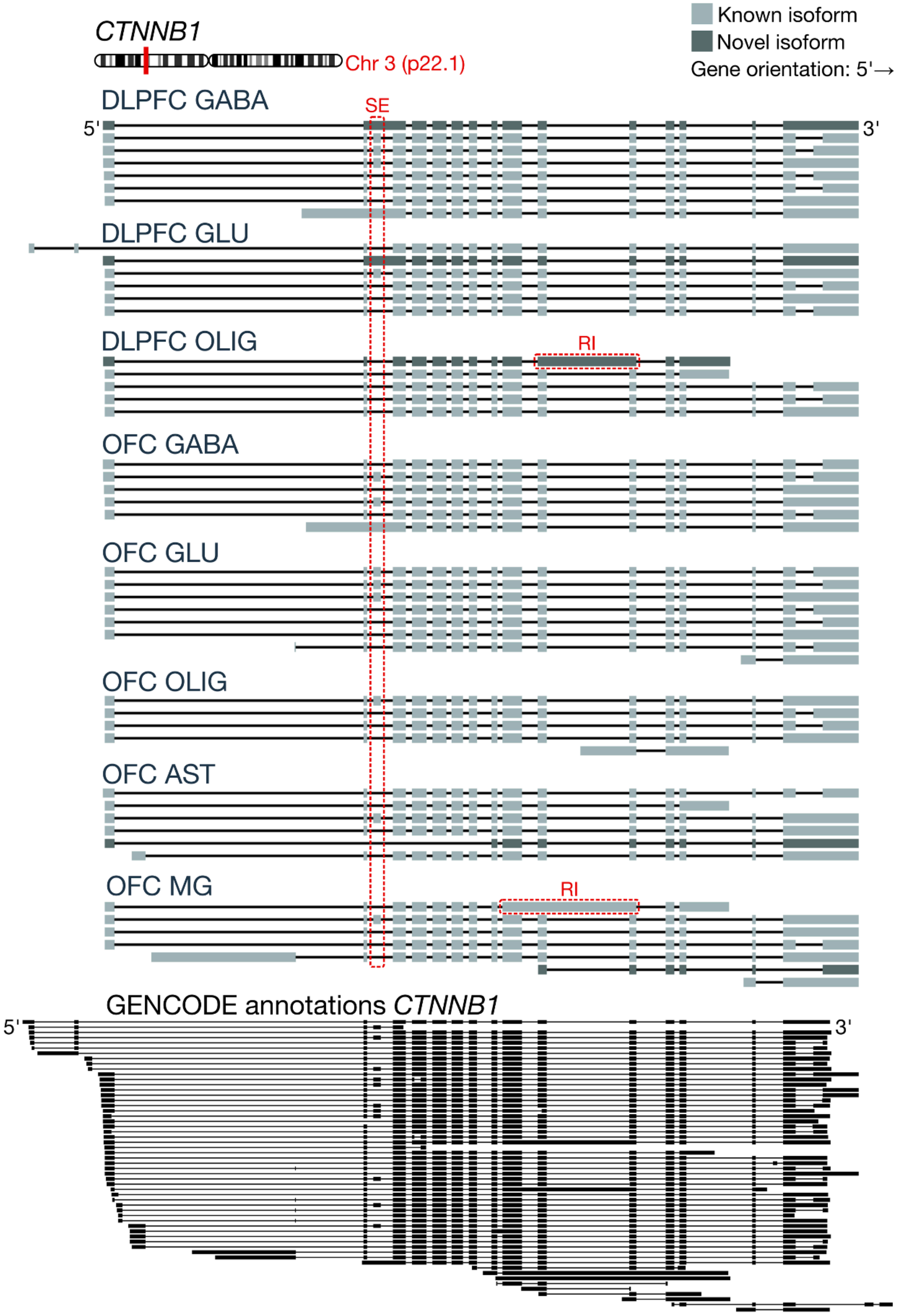

CTNNB1
Chr 3 (p22.1)
Known isoform
Novel isoform
Gene orientation: 5'→ 3'
DLPFC GABA
SE
5'
3'
DLPFC GLU
DLPFC OLIG
RI
OFC GABA
OFC GLU
OFC OLIG
OFC AST
OFC MG
RI
GENCODE annotations CTNNB1
5'
3'

**Figure S16**. ***CTNNB1* isoforms with RI and SE across cortical cell types**. Shown are full-length Iso-Seq reconstructions of *CTNNB1* across major cortical cell types in the DLPFC and OFC, illustrating extensive isoform heterogeneity at a single locus. Known (FSM/ISM) and novel (NIC/NNC) isoforms are displayed in gray and dark gray, respectively. Red dashed boxes highlight two recurrent alternative splicing features: (i) a shared exon-skipping (SE) event present across neuronal and glial lineages, and (ii) distinct retained-intron (RI) events that appear selectively in OLIG and MG populations.

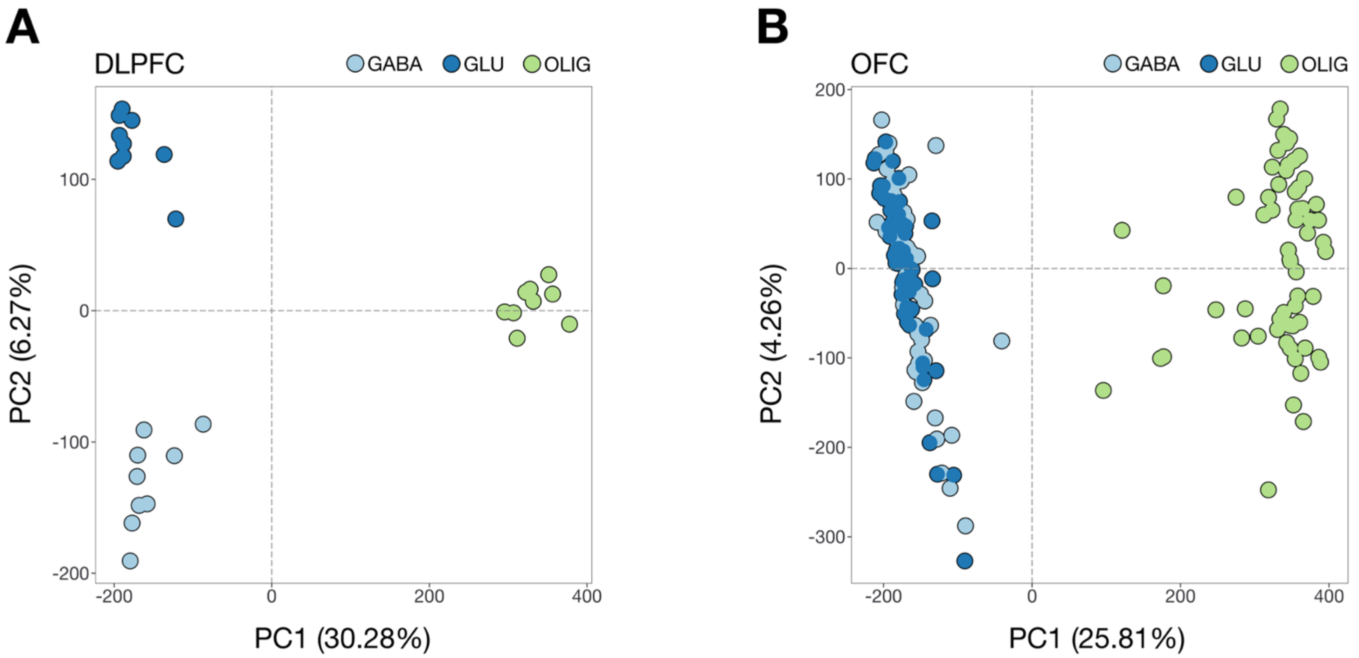


**Figure S17. Principal component analysis (PCA) of Illumina short-read RNA-seq data**. (**A**, **B**) Clustering of samples by cell type in the DLPFC and OFC, respectively. Principal component 1 (PC1) is shown on the x-axis and principal component 2 (PC2) on the y-axis.

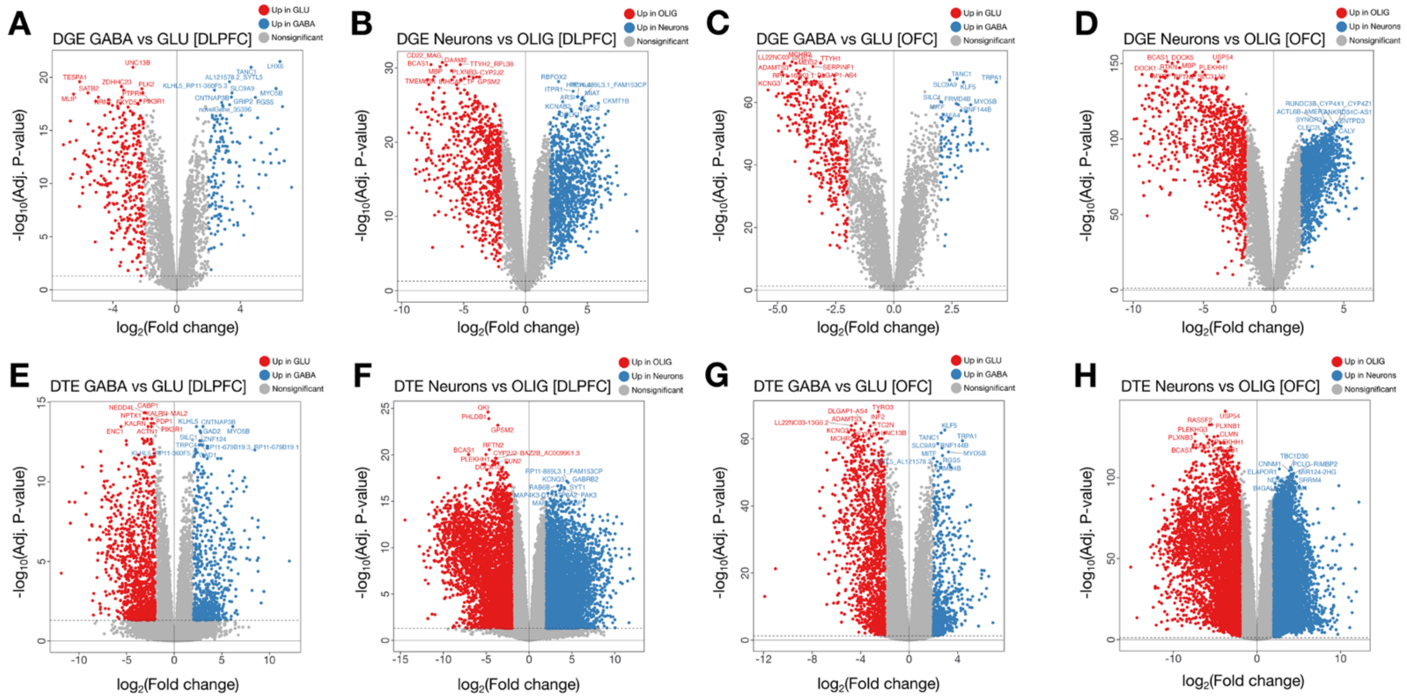


**Figure S18. Differential gene expression (DGE) and differential transcript expression (DTE) analysis**. (**A**, **C**) DGE between GABAergic and glutamatergic neurons in the DLPFC and OFC. (**B**, **D**) DGE between neurons (GABAergic and glutamatergic neurons combined) versus oligodendrocytes in the DLPFC and OFC. (**E**, **G**) DTE between GABAergic and glutamatergic neurons in the DLPFC and OFC. (**F**, **H**) DTE between neurons and oligodendrocytes in the DLPFC and OFC. Volcano plots show $\log_2$ fold change ($\log_2$FC) on the x-axis and $-\log_{10}$ adjusted p-value (Benjamini-Hochberg) on the y-axis. Significant genes or transcripts were defined as those with false discovery rate (FDR) < 5% and $|\log_2FC|>2$.

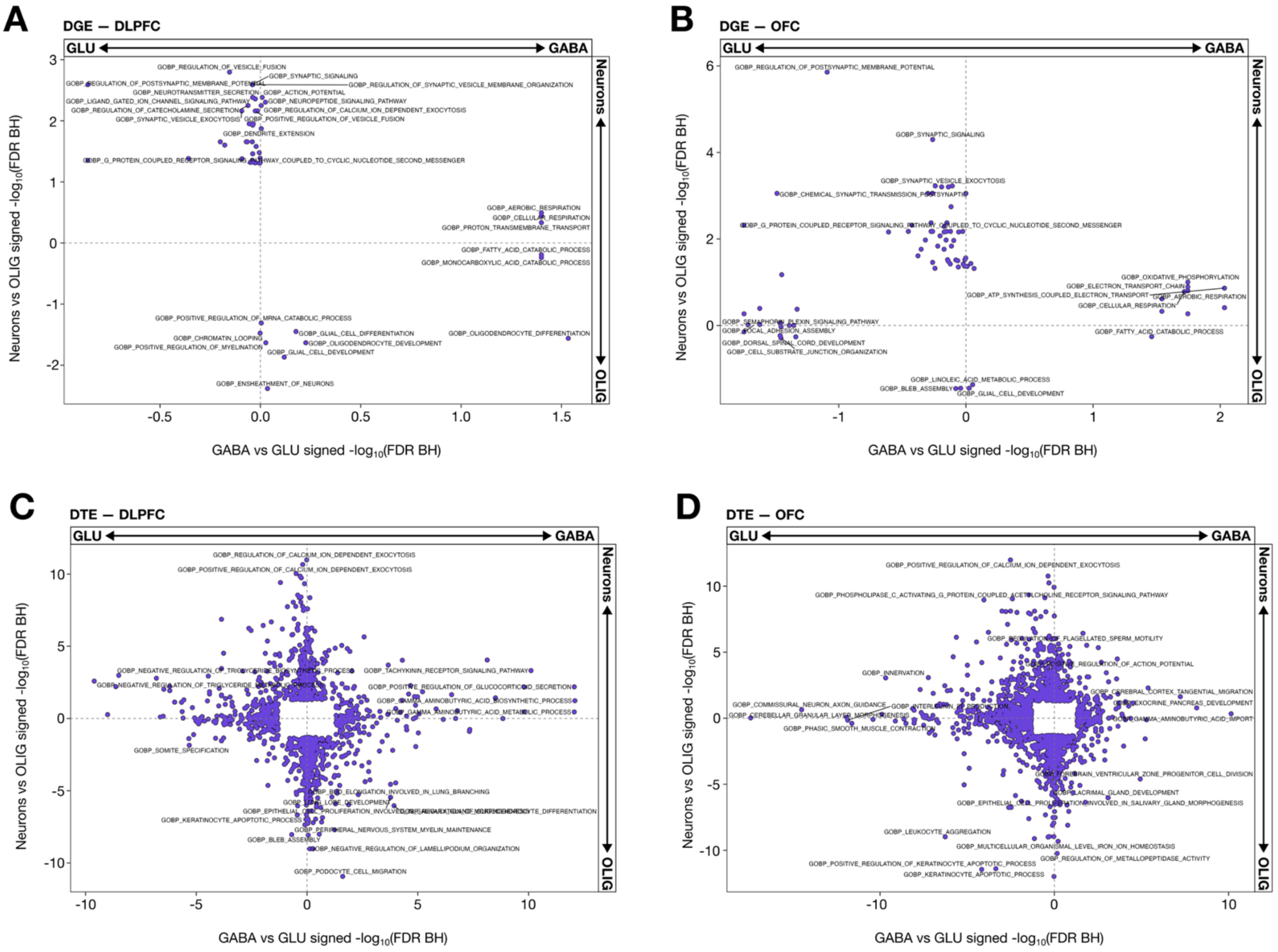


**Figure S19. DGE and DTE CAMERA gene ontology (GO) biological processes (BP) enrichment plots across cortical regions**. (**A**, **B**) DGE CAMERA GO BP enrichment plots for the DLPFC and OFC. (**C**, **D**) DTE CAMERA GO BP enrichment plots for the DLPFC and OFC. GO enrichment is plotted as signed -log10 false discovery rate (FDR, Benjamini-Hochberg). The y-axis represents the comparison of neurons versus oligodendrocytes, and the x-axis represents the comparison of GABAergic versus glutamatergic neurons. Positive and negative values indicate enrichment in the first and second group, respectively. Quadrants indicate bias: Q1, GABA-enriched; Q2, GLU-enriched; Q3-Q4, OLIG-enriched.

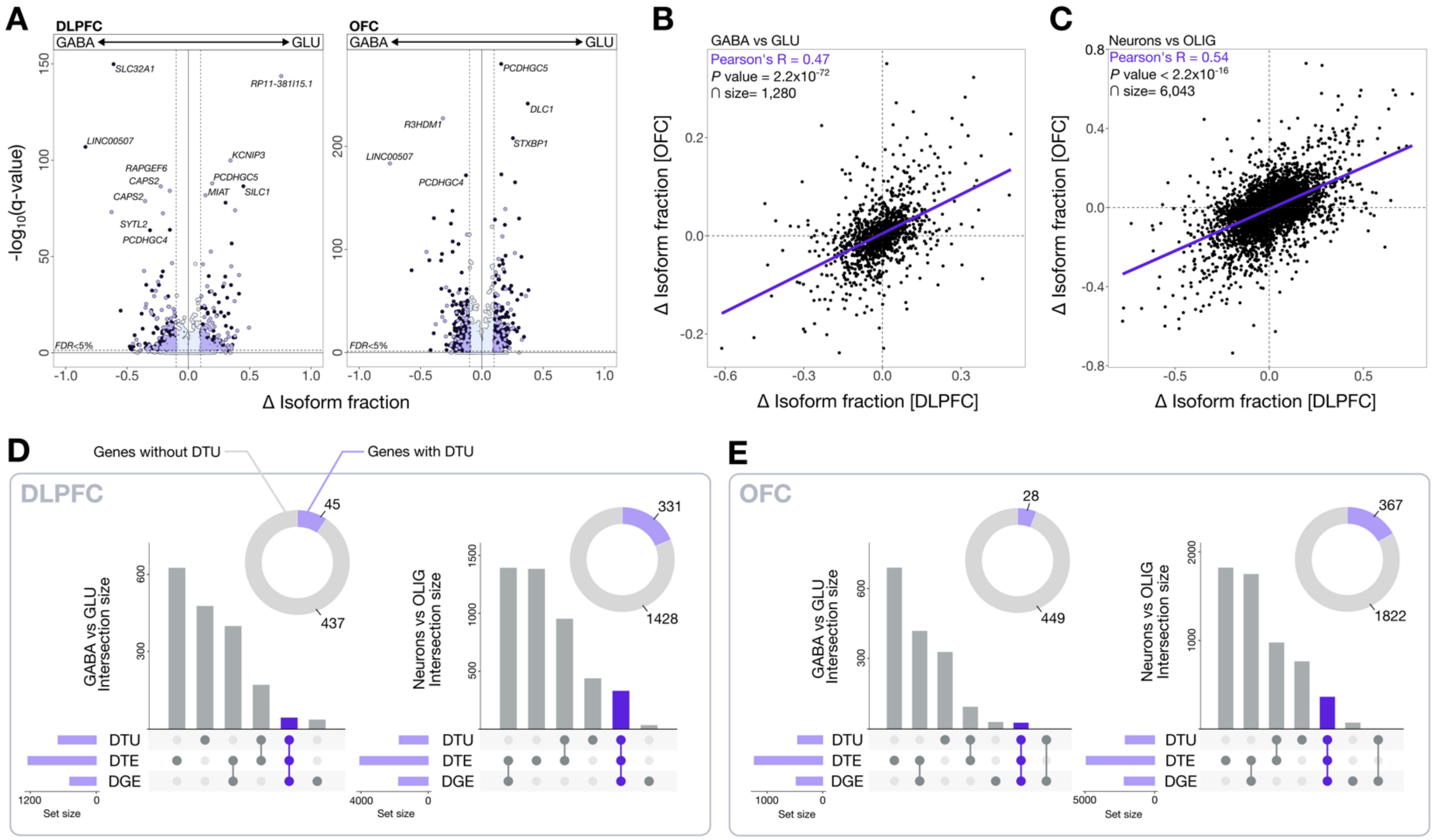


**Figure S20. Differential transcript usage (DTU) analysis across cortical cell types**. (**A**) Volcano plots showing differential isoform fraction on the x-axis and FDR (Benjamini-Hochberg adjustment) on the y-axis, comparing GABAergic and glutamatergic neurons in the DLPFC and OFC. Significant DTUs were defined as those with FDR < 5% and Δ isoform fraction > 10%. (**B**) Concordance of significant DTUs between GABAergic and glutamatergic neurons in the DLPFC and OFC. (**C**) Concordance of significant DTUs between neurons and oligodendrocytes in the DLPFC and OFC. (**D**) Upset plots showing overlap of significant DGE, DTE, and DTU at the gene level for GABAergic versus glutamatergic neurons and neurons versus oligodendrocytes in the DLPFC (left) and OFC (right).

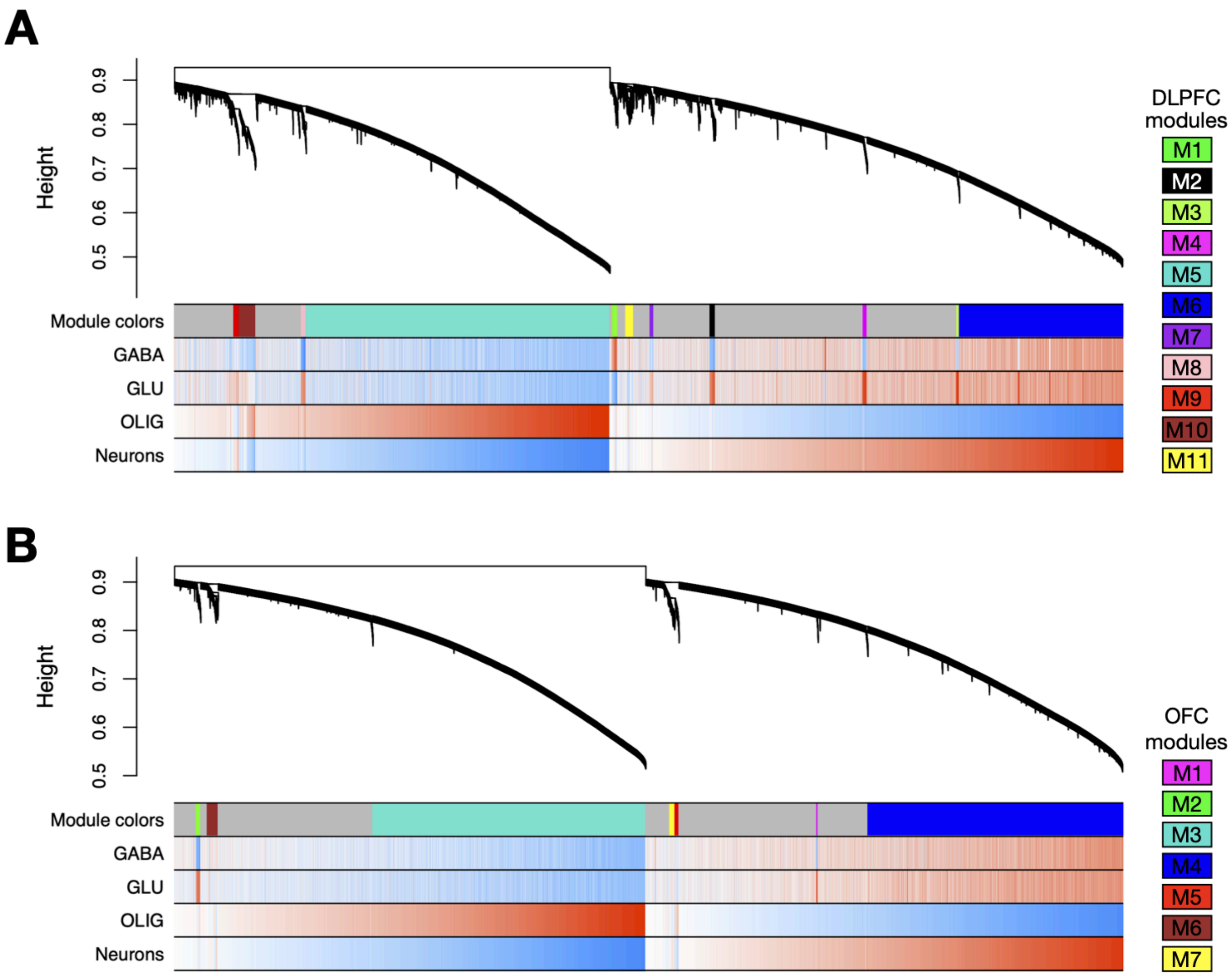


**Figure S21. Isoform-level weighted gene co-expression network analysis (WGCNA) across cortical cell types in the DLPFC and OFC.** Hierarchical clustering dendrograms of isoform expression used to define WGCNA co-expression modules in the (A) DLPFC and (B) OFC. Dynamic tree cutting identified 11 modules in the DLPFC and 7 in the OFC, each represented by a unique module color bar aligned beneath the dendrogram. Below each module color track, heatmaps display the correlation of module eigengenes with major cortical lineages, GABA, GLU, OLIG, and a combined "pan-neuronal" category, highlighting cell type-specific enrichment patterns. Warmer colors indicate positive correlations, and cooler colors indicate negative correlations.

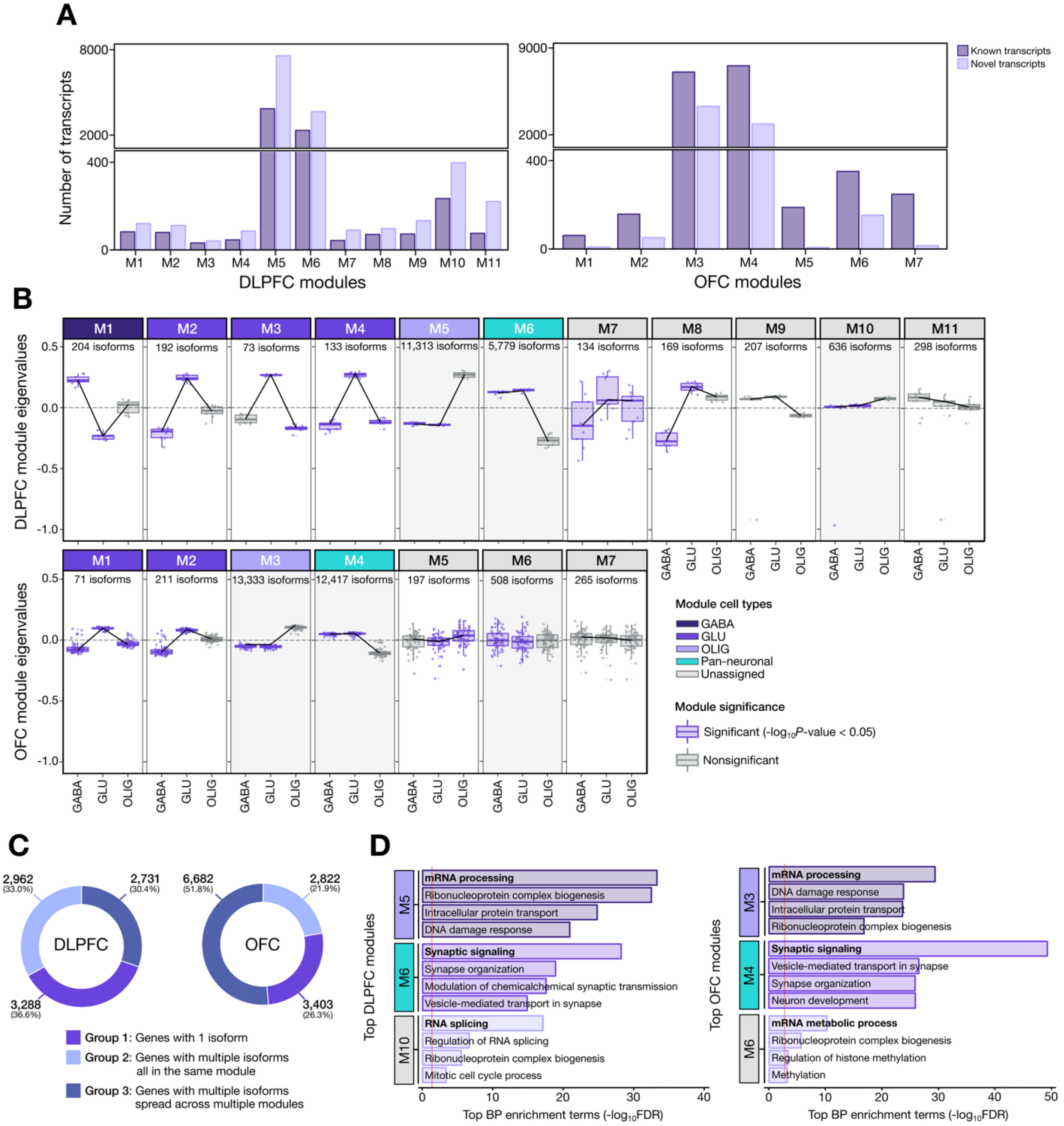


**Figure S22. Isoform-level co-expression modules across cortical cell types.** (**A**) Isoform co-expression modules identified by WGCNA in the DLPFC (11 modules) and OFC (7 modules). Each bar shows the total number of isoforms per module, stratified by known versus novel transcripts. Horizontal lines denote the median module size. (**B**) Module eigengene profiles across major cell types (GABA, GLU, OLIG, pan-neuronal, and unassigned) in each cortical region. Boxes indicate the distribution of eigengene values for each cell type; significant associations (FDR $p < 0.05$) are marked in purple. (**C**) Distribution of genes containing one isoform, multiple isoforms within a single module, or multiple isoforms spread across ≥2 modules. Approximately ~20% of multi-isoform genes distribute across multiple modules, indicating functional divergence at the isoform level. (**D**) Gene Ontology biological process enrichments for the most isoform-dense modules in each region.

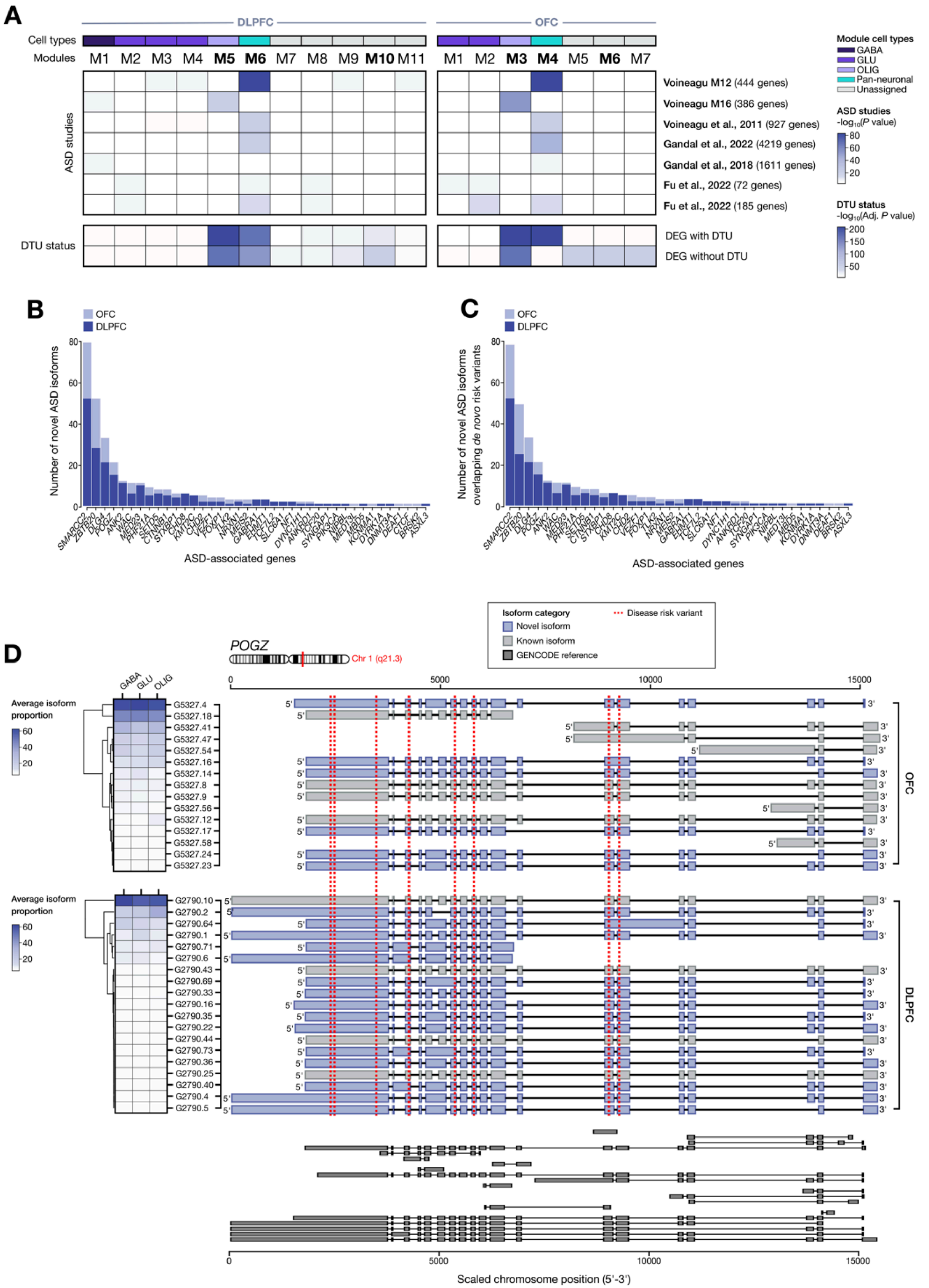
A
DLPFC
OFC
Cell types
Modules
M1 M2 M3 M4 M5 M6 M7 M8 M9 M10 M11
M1 M2 M3 M4 M5 M6 M7
ASD studies
Voineagu M12 (444 genes)
Voineagu M16 (386 genes)
Voineagu et al., 2011 (927 genes)
Gandal et al., 2022 (4219 genes)
Gandal et al., 2018 (1611 genes)
Fu et al., 2022 (72 genes)
Fu et al., 2022 (185 genes)
DTU status
DEG with DTU
DEG without DTU
Module cell types
GABA
GLU
OLIG
Pan-neuronal
Unassigned
ASD studies
-log10(P value)
80
60
40
20
DTU status
-log10(Adj. P value)
200
150
100
50
B
OFC
DLPFC
Number of novel ASD isoforms
ASD-associated genes
C
OFC
DLPFC
Number of novel ASD isoforms overlapping de novo risk variants
ASD-associated genes
D
POGZ
Chr 1 (q21.3)
Isoform category
Novel isoform
Known isoform
GENCODE reference
Disease risk variant
Average isoform proportion
GABA
GLU
OLIG
OFC
DLPFC
Scaled chromosome position (5'-3')

**Figure S23. Isoform-resolved convergence of cortical co-expression modules with ASD-associated genes.** (**A**) Heatmaps showing the overlap between DLPFC and OFC isoform co-expression modules and ASD-associated gene sets derived from major genetic and transcriptomic studies. Modules are annotated by their dominant cell type (GABA, GLU, pan-neuronal, OLIG, or unassigned). For each module, enrichment is shown separately for genes with DTU and those without DTU, highlighting isoform-specific regulatory contributions to ASD risk. (**B**) Number of novel isoforms detected within ASD-associated genes in the refined DLPFC and OFC transcriptomes, sorted by gene, illustrating extensive unannotated isoform diversity in ASD-relevant loci. (**C**) Novel isoforms from ASD-associated genes that overlap de novo variants from ASD exome-sequencing studies, highlighting isoform structures that directly intersect risk-bearing sites. (**D**) Isoform structures for *POGZ*, including both known and novel transcripts, shown alongside average isoform usage across GABA, GLU, and OLIG cell types in each cortical region. Red dashed lines mark ASD-associated de novo variants mapping to specific exons/isoforms

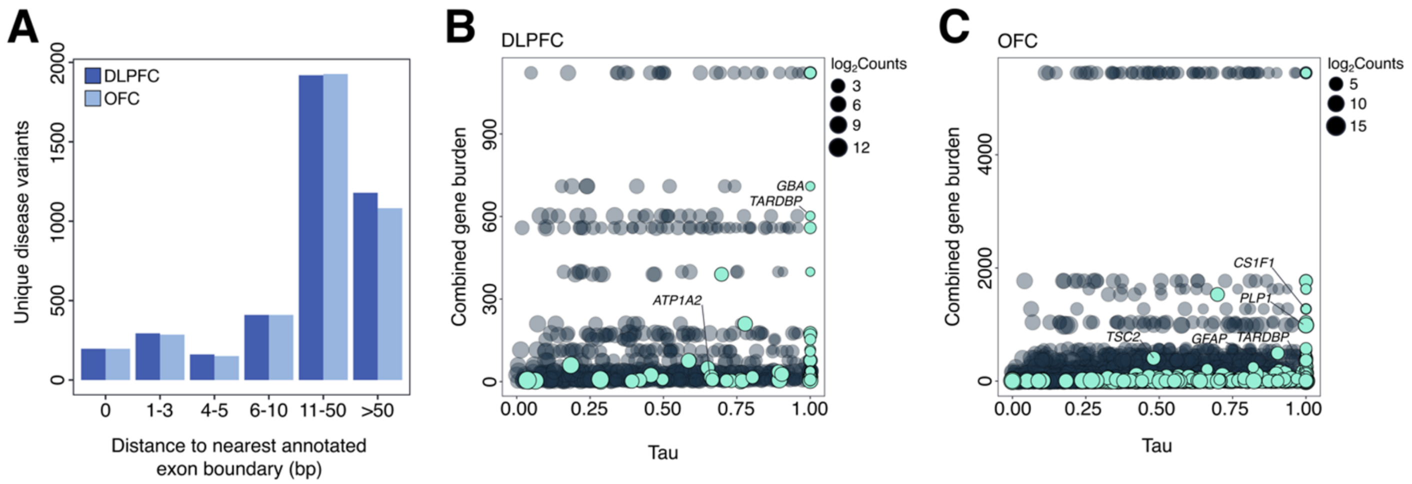


**Figure S24. Disease-associated variants are enriched near novel splice boundaries and concentrated in cell type–specific genes. (A)** Distribution of HGMD disease-causing variants relative to the nearest annotated exon boundary for novel splice-site events in DLPFC and OFC. Variants were binned by distance to the closest annotated exon boundary (0, 1–3, 4–5, 6–10, 11–50, and >50 bp). The majority of variants localized near splice boundaries, with the strongest enrichment observed within 11–50 bp of annotated exon boundaries, consistent with pathogenic variation being concentrated around novel donor, acceptor, and exon extension events. **(B-C)** Relationship between gene-level cell type specificity (tau) and combined disease variant burden for genes harboring novel splice-site events in DLPFC (B) and OFC (C). Tau values approaching 1 indicate strong cell type–restricted expression, whereas values near 0 indicate broad expression across lineages. Point size reflects total gene expression (log2 counts). Highlighted genes with high splice-site burden and strong cell type specificity include GBA, TARDBP, and ATP1A2 in DLPFC, and CSF1R, PLP1, TSC2, GFAP, and TARDBP in OFC. These results indicate that disease-associated splice architecture is frequently concentrated in lineage-restricted genes, supporting a model in which novel isoform structures contribute to selective cellular vulnerability in neurological disease.
