## Supplemental Results for "Long-read transcriptomics of purified human cortical cell types exposes glial isoform complexity and disease-relevant transcript architecture"

### Isoform-level co-expression networks reveal cell type-specific functional modules

To explore coordinated regulation of isoforms across cortical cell types, we performed weighted gene co-expression network analysis using isoform-level expression profiles separately for the DLPFC and OFC. This analysis resolved 11 co-expression modules in the DLPFC and 7 modules in the OFC, incorporating both annotated and newly discovered isoforms (**Figure S21-22, Table S19**). Module-cell type correlations revealed strong and reproducible cell type-specific organization. In the DLPFC, six modules showed clear lineage biases, including an OLIG-enriched module (M5; $r = 0.99$, $p = 4.9\times10^{-29}$) and a pan-neuronal module (M6; $r = 0.49$, $p = 0.01$) with coordinated enrichment across both GABA and GLU neurons (**Figure S22B**, **Table S20**). The OFC showed similar structure, with OLIG-linked M3 and pan-neuronal M4 exhibiting analogous patterns. Across multi-isoform genes, ~20% had transcripts partitioning across ≥2 modules (**Figure S22C**), indicating that isoforms from the same gene often participate in distinct regulatory programs.

We next evaluated the biological functions represented within these modules. The most isoform-dense modules displayed strikingly coherent enrichment patterns that were broadly conserved across regions. OLIG-enriched modules (DLPFC M5; OFC M3) were dominated by RNA processing, ribonucleoprotein complex assembly, and mRNA metabolic pathways (**Figure S22D**), functions consistent with the extensive splicing and RNA remodeling observed in oligodendrocytes. In contrast, neuronal modules (DLPFC M6; OFC M4) displayed strong enrichment for synaptic signaling, vesicle transport, cytoskeletal organization, and other processes central to neuronal communication (**Figure S22D**). Modules without clear lineage assignment also exhibited region-specific distinctions: the unassigned DLPFC module (M10) was enriched for RNA splicing-related processes (FDR < 5%), whereas the comparable OFC module (M6) was enriched for mRNA metabolic and methylation pathways, suggesting subtle regional tuning of post-transcriptional regulation. These isoform-level co-expression networks uncover a hierarchical and cell type-structured organization of transcript regulation across the human cortex. The strong convergence of module functions across regions underscores conserved mechanisms of isoform-mediated regulation, whereas region-specific enrichments point to fine-grained differences in post-transcriptional programs that tailor cellular specialization in distinct cortical areas.

### Isoform-level signatures of neurodevelopmental disorder risk

To assess whether isoform co-expression architecture captures neurodevelopmental disease biology, we next evaluated each module for enrichment of autism spectrum disorder (ASD) risk genes and ASD-linked differential expression signatures[30-32]. Significant enrichments were detected across multiple isoform co-expression modules (**Figure S23A**), demonstrating that ASD risk loci and transcriptomic perturbations preferentially converge within neuronally enriched isoform networks. Pan-neuronal modules M6 (DLPFC) and M4 (OFC) showed the strongest and most consistent enrichment across independent ASD gene lists (FDR ≤ 0.001), indicating shared, lineage-specific axes of isoform dysregulation reproducibly identified in both cortical regions. OLIG-enriched modules (M5 in the DLPFC; M3 in the OFC) also showed meaningful overlap with ASD-associated genes, suggesting that OLIG-specific splicing programs, though less prominently implicated than neuronal modules, may nonetheless contribute to ASD-relevant transcriptome remodeling.

Building from this, we integrated high-confidence genetic risk variants for autism spectrum disorder and identified 39 ASD-associated genes for which we uncovered novel isoforms (**Figure S23B**). Notably, 38 of these novel isoforms intersected *de novo* ASD variants within extended or isoform-specific exonic regions (**Figure S23C**), directly linking previously unannotated transcript structures to genetic risk. Several high-confidence ASD genes, including *SMARCC2*, *FOXP1*, *DYRK1A*, *CHD8*, and *POGZ* displayed extensive cell-type-specific isoform diversity (**Table S22**), underscoring the importance of isoform-level annotation for accurately interpreting disease-associated variants. Among these, *POGZ*, a recurrent ASD-risk gene encoding a chromatin regulator essential for neuronal differentiation, showed pronounced isoform heterogeneity across both the DLPFC and OFC (**Figure S23D**). Novel *POGZ* isoforms incorporated unique exon combinations that directly intersected de novo ASD variants (red dashed lines), highlighting regulatory complexity not visible from canonical annotations. The cell-type-resolved isoform profiles further suggest that these novel transcripts arise from distinct, lineage-specific splicing events, consistent with a model in which isoform usage modulates chromatin-regulatory functions in a context-dependent manner (**Table S23**).
